# 2-Deoxyglucose dendrimer-enabled niclosamide delivery to FRβ-expressing macrophages alleviates endometriosis progression and associated hyperalgesia

**DOI:** 10.64898/2026.02.06.704463

**Authors:** Anubhav Dhull, Mingxin Shi, Madeleine E. Harvey, Taylor M. Page, Rishi Sharma, Kenneth John Goody, Aqib Iqbal Dar, Nash S. Moawad, Vijay Kumar Sirohi, Paul S. Cooke, Kanako Hayashi, Anjali Sharma

**Author notes:** Correspondence: Anjali Sharma, Department of Chemistry, College of Arts and Sciences, Washington State University, Pullman, WA 99164, USA. Kanako Hayashi, School of Molecular Biosciences, Center for Reproductive Biology, Washington State University, Pullman, WA 99164, USA. These authors contributed equally to this work (Anubhav Dhull, Mingxin Shi, Madeleine E. Harvey).

## Abstract

Endometriosis is a chronic, incurable disease. Due to limited efficacy, high recurrence rates, and serious side effects of current treatments, development of new, targeted, non-hormonal therapies is urgently needed. We previously reported that niclosamide, an FDA-approved anthelmintic drug, attenuates endometriotic lesion growth. We further identified folate receptor-β (FRβ)-positive macrophages as contributors to disease progression. Significantly, niclosamide inhibits FRβ^+^ macrophages and reduces inflammation, innervation, and angiogenesis. To develop niclosamide as a non-hormonal and selective immune cell-targeted therapy for endometriosis, we engineered a folic acid-conjugated 2-deoxyglucose dendrimer (FA-2DG-D) using click chemistry to enable selective FRβ-mediated uptake. Conjugation of niclosamide to FA-2DG-D yielded a targeted nanotherapeutic (FA-2DG-D-Niclo) with enhanced aqueous solubility, controlled intracellular release, and excellent batch-to-batch reproducibility. In a mouse model of endometriosis, FA-2DG-D demonstrated lesion-specific accumulation and selective internalization by FRβ⁺ macrophages with minimal off-target organ retention. A single intraperitoneal dose of FA-2DG-D-Niclo (25 or 50 mg/kg/bw of niclosamide) significantly reduced FRβ⁺ macrophage burden, suppressed lesion number and volume, and markedly improved endometriosis-associated hyperalgesia at two weeks post-treatment. Together, these findings establish FRβ⁺ macrophages as a potential target in endometriosis and present FA-2DG-D-Niclo as a non-hormonal, macrophage-focused nanomedicine for precise and effective endometriosis treatment.

**Graphic Abstract:** 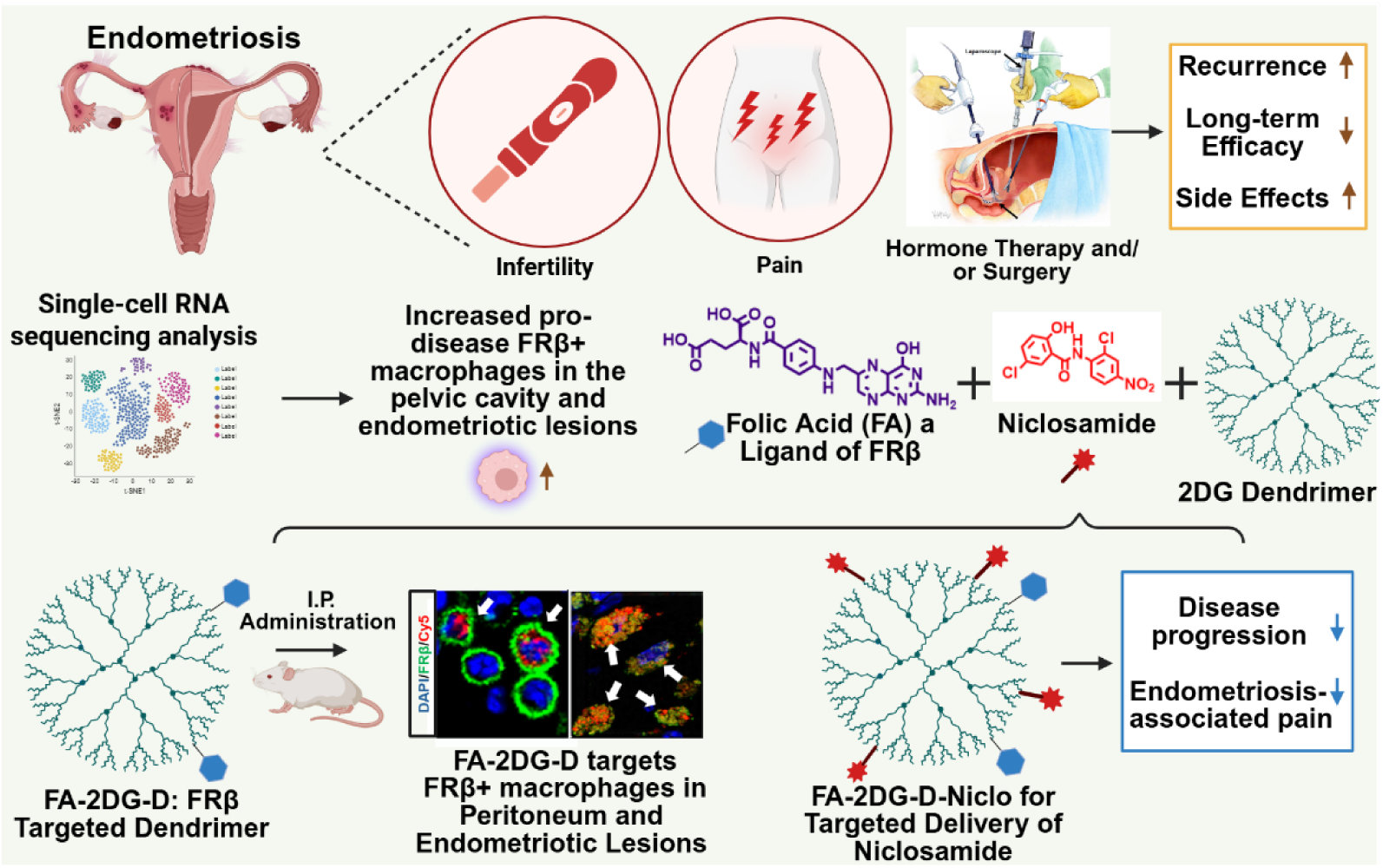

## 1. Introduction

Endometriosis affects an estimated 10% of reproductive-aged women, roughly 190 million individuals worldwide, and remains one of the most debilitating gynecological disorders [1]. The disease is characterized by chronic pelvic pain, dysmenorrhea, dyspareunia, and infertility, all of which severely diminish the quality of life for patients and their families [2–4]. In the United States alone, the annual economic burden is estimated at $78-119 billion, encompassing healthcare costs, productivity losses, and reduced workforce participation [5]. Despite its prevalence and impact, endometriosis remains incurable, and the therapeutic landscape has changed little over the past several decades. Current medical management relies on hormonal suppression of estrogen production or action [6, 7], such as oral contraceptives, progestins, and/or gonadotropin-releasing hormone (GnRH) agonists/antagonists [6, 8], as well as surgical excision of lesions. However, hormonal treatments are limited in efficacy, have high recurrence rates, and cause significant side effects. Surgical excision offers temporary relief of symptoms, yet nearly half of surgically treated women experience recurrence within five years, and ∼12% fail to achieve meaningful pain reduction after surgery [9, 10]. Thus, there is an urgent need to identify new therapeutic targets to develop novel non-hormonal treatments in endometriosis.

A growing body of evidence implicates immune dysfunction, particularly macrophage dysregulation, in driving lesion establishment, chronic inflammation, angiogenesis, and neurogenesis in endometriosis [11–13]. Recently, our single-cell transcriptomic analysis identified a unique FRβ-expressing macrophage population, differentiated from monocyte-derived proinflammatory macrophages and characterized by residential macrophage traits in mouse endometriosis [14]. The population of macrophages is also identified in the lesions of endometriosis patients [15]. The selective upregulation of FRβ^+^ on these macrophages can be used as an accessible and disease-restricted cellular target for ligand-guided targeted drug delivery.

Our prior studies demonstrated that niclosamide significantly suppressed endometriotic lesion growth, alleviated aberrant peritoneal inflammation, and decreased vascularization and innervation in lesions using two distinct mouse models of endometriosis without compromising fertility [13, 14, 16]. Niclosamide (5-chloro-*N*-(2-chloro-4-nitrophenyl)-2-hydroxybenzamide) is a chlorinated salicylanilide derivative that is FDA-approved as an anthelmintic drug. Currently, multiple clinical trials are ongoing to repurpose it for the treatment of other diseases, including cancer and metabolic diseases [17]. However, the clinical use of niclosamide is significantly limited by its poor aqueous solubility and correspondingly low oral bioavailability. To address these challenges, researchers are actively developing innovative formulation strategies such as water-soluble derivatives, solid dispersions, and nanocarrier-based delivery systems to enhance its solubility, pharmacokinetics and overall therapeutic potential [18]. Additionally, systemic niclosamide treatment inhibits several pathways, including STAT3, NFκB, and WNT signaling, across all cell types [11, 12, 16], including other macrophages [14]. Disrupting these essential pathways across all cell types would lead to unwanted side effects and this poses a significant barrier to therapeutic use of niclosamide. Importantly, niclosamide can also reduce elevated FRβ⁺ macrophages and restore the perturbed microenvironment in endometriosis by finely regulating macrophage dynamics [14], suggesting that FRβ⁺ macrophages can serve as a viable therapeutic target in endometriosis. Therefore, targeted delivery of niclosamide to FRβ⁺ macrophages offers a strategy to maximize therapeutic efficacy while minimizing systemic toxicity.

In this context, dendrimer nanotechnology has emerged as a powerful tool for targeted therapeutic delivery, enabling selective engagement of pathogenic cells while minimizing off-target toxicity [19–24]. Dendrimers are monodisperse, highly branched macromolecules whose multivalent surface architecture and tunable chemistry allow precise conjugation of targeting ligands, therapeutic agents, and imaging probes [25–27]. Building on our previously synthesized 2-deoxy-glucose (2DG) functionalized dendrimer (2DG-D) [28], which incorporates 2DG moieties to enhance solubility and macrophage interaction, we engineered a folic acid (FA), a ligand of FRβ, conjugated dendrimer (FA-2DG-D) by covalently coupling FA to the dendrimer surface through orthogonal click chemistry. This design exploits ligand-FRβ binding to selectively direct dendrimer uptake into FRβ⁺ macrophages within endometriotic lesions and peritoneum [29, 30]. To achieve targeted delivery, we further developed the niclosamide conjugated FA-2DG-D (FA-2DG-D-Niclo). Here, we report the design, synthesis, and in vivo evaluation of FA-2DG-D and FA-2DG-D-Niclo in a mouse model of endometriosis. Our results demonstrate that FA-2DG-D has FRβ⁺ macrophage-specific targeting in the lesions and peritoneum and delivers niclosamide into FRβ⁺ macrophages with ligand-receptor internalization. This targeted dendrimer platform offers a promising strategy for therapeutic delivery in endometriosis, addressing a critical unmet need for more precise and effective interventions.

## 2. Methods

### 2.1. Human Sample Collection

All procedures involving human subjects were performed in accordance with ethical approval by the Institutional Review Board (IRB) guidelines for the University of Florida under IRB code NO. IRB202003131. All patients were recruited from the Division of Minimally Invasive Gynecologic Surgery of the University of Florida Health Shands Hospital, Gainesville, Florida. Participants were either previously diagnosed with endometriosis or suspected of having endometriosis, with symptoms including ovarian cysts, chronic pelvic pain, dysmenorrhea, dyschezia, dyspareunia, uterine bleeding, and myofascial pain syndrome. Endometriosis is further confirmed by pathological analysis of the excised lesions obtained during surgery. Informed written consent was obtained from each participant before the procedure. In this study, peritoneal washings from 4 patients aged 19 to 37 years (**Supplementary Table S1**) were collected at the beginning of the procedure. Briefly, the pelvic cavity was irrigated using approximately 100 mL of sterile saline, and approximately 80 mL of peritoneal washings containing pelvic exudate cells were collected from each patient and placed on ice immediately. Red blood cells were removed by resuspending cells with 500 μL Red Blood Cell Lysis Buffer (BioLegend, Cat.#420301). Cells were then counted and cryopreserved with freezing media containing 90% fetal bovine serum (FBS) and 10% dimethyl sulfoxide, and stored at −80 °C for further analysis.

### 2.2. Single-cell RNA-Sequencing and Analysis

#### 2.2.1. Library Preparation and Sequencing

To perform single-cell RNA sequencing (scRNA-seq), frozen pelvic exudate cells were thawed in a 37 °C water bath and prepared following the 10x Genomics protocol named “Fresh frozen human peripheral blood mononuclear cells for single-cell RNA sequencing” (document #CG00039 Rev E). Briefly, thawed cells were slowly transferred into a 50 mL conical tube and recovered by sequential 1:1 dilution with warmed FBS for a total of 5 times. Cell viability was assessed to ensure samples with viability greater than 80%. After centrifuging at 300g for 5 min at room temperature, the cells were resuspended in phosphate-buffered saline (PBS) with 0.04% bovine serum albumin (BSA) at a concentration of 900-1,200 cells/μl. Cells were then placed on ice and immediately proceeded to library construction using a 10x Genomics Chromium Next GEM Single Cell 3’ Kit v3.1, following the standard kit user guide (document #CG000204) for a target of 6000 cells. Libraries were sequenced on Illumina NovaSeq 6000. FASTQ files were aligned to the human reference genome GRCh38 via Cell Ranger v7.1.0 [31], and an average of 107,903 reads per cell was achieved for each RNA library (**Supplementary Table S2**).

#### 2.2.2. External Human Endometrial and Pelvic Exudate Cell scRNA-seq Datasets

We integrated publicly available human scRNA-seq datasets from eutopic endometrium with or without endometriosis, ectopic endometrium, and pelvic exudate cells. Processed Cell Ranger output files or raw FASTQ files were downloaded from the Gene Expression Omnibus (GEO), ArrayExpress, or the Sequence Read Archive (SRA). These datasets included: (1) Wang et al. (GSE111976) [32], (2) Lai et al. (GSE183837) [33], (3) Garcia-Alonso et al. (E-MTAB-10287) [34], (4) Tan et al. (GSE179640) [15], (5) Fonseca et al. (GSE213216) [35], (6) Huang et al. (GSE214411) [36], (7) Marečková et al. (E-MTAB-14039) [37], (8) Shin et al. (PRJNA932195) [38], (9) Han et al. (GSE228030) [39], and (10) Zou et al. (PRJNA713993) [40], as shown in **Figure 1A and B**. For all datasets, only samples collected from women of reproductive age were included. For endometrial scRNA-seq datasets from (1) to (8), patients labeled as undergoing hormonal treatments during sample collection were also excluded. Samples used in this study are summarized in **Supplementary Table S3**.

**Figure 1.**
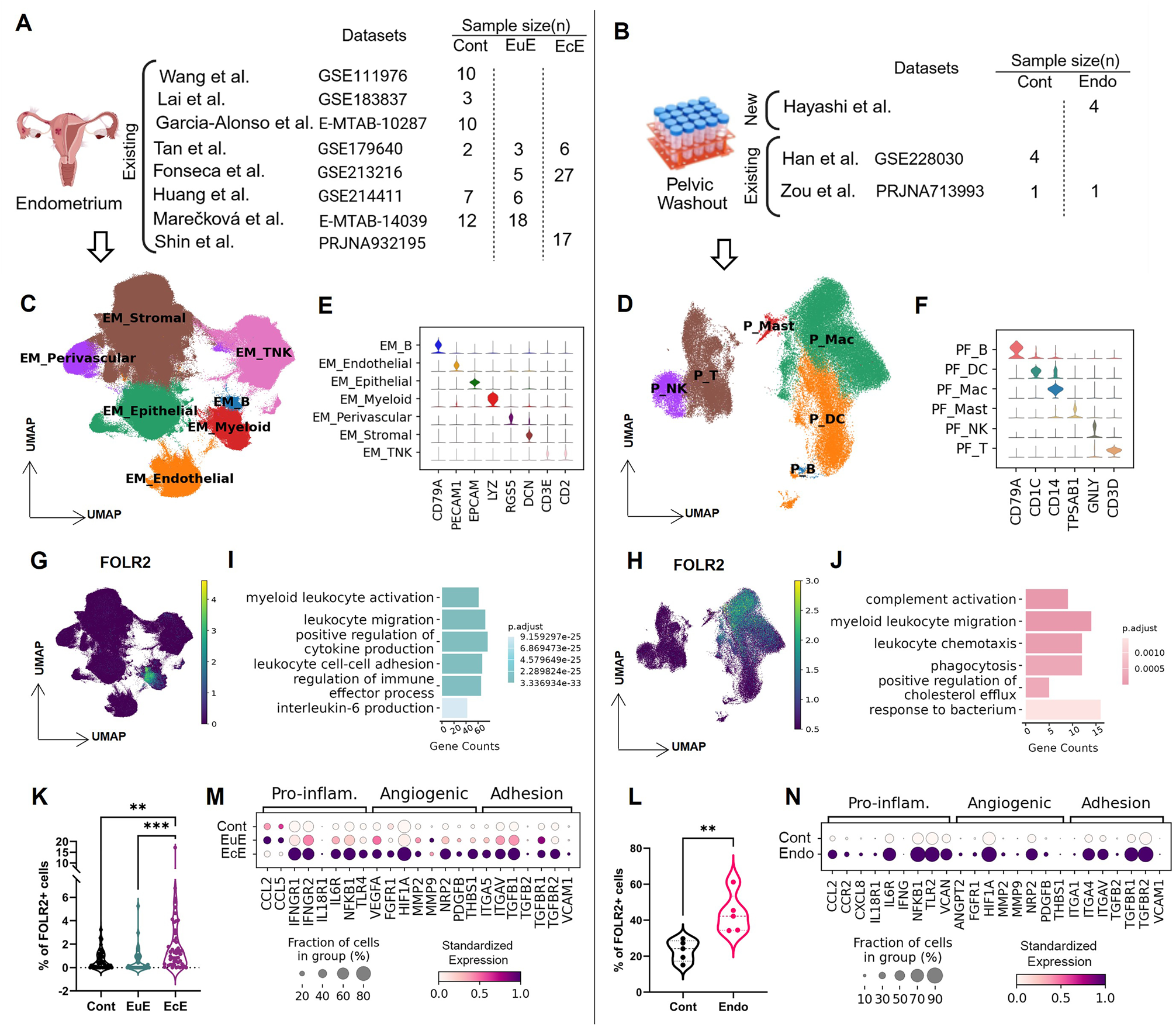
scRNA-seq analysis of FOLR2^+^ macrophages in endometrium and pelvic exudate cells in endometriosis. (**A, B**) Lists of human scRNA-seq datasets analyzed and sample size per dataset in the groups of (**A**) eutopic endometrium with (EuE) and without (Cont) endometriosis, ectopic endometrium (lesion, EcE), and (**B**) pelvic exudate cells from subjects with (Endo) and without (Cont) endometriosis. (**C, D**) UMAP plots showing major cell types identified in the endometrium (EM) and pelvic cells (P). (**E, F**) Violin plot representing marker gene expression for each major cell type identified in endometrial and pelvic cell datasets. (**G, H**) UMAP projections showing expression patterns of FOLR2 in endometrial and pelvic cells. (**I, J**) Bar plots showing enriched gene ontology (GO) biological processes in FOLR2^+^ macrophages from endometrium and pelvic cells. (**K, L**) Proportions of endometrial and pelvic FOLR2-expressing cells to total cells within each tissue type. Differences among groups were analyzed with one-way ANOVA followed by the Tukey multiple comparisons test. **P<0.01, ***P<0.001. (**M, N**) Dot plot showing representative significantly differentially expressed genes (DEGs) involved in proinflammatory, angiogenesis, and adhesion processes among groups of endometrium and pelvic cells (adjusted p-value <0.05).

#### 2.2.3. Single-cell Data Pre-processing and Clustering

RNA matrices from both newly generated and existing public datasets were individually imported to Scanpy v.1.8.1 for further analysis [41]. Low-quality cells were excluded using the following criteria: (1) genes that were detected in fewer than 3 cells, (2) cells with fewer than 300 genes, (3) a maximum of 100,000 UMIs, and (4) a maximum of 20% mitochondrial content. Doublets were detected and removed using Scrublet [42]. Filtered matrices from endometrial or pelvic cell scRNA-seq datasets were combined separately and analyzed independently. For each combined endometrial or pelvic cell scRNA-seq matrix, gene expression values were normalized with a global scaling factor of 10, 000 and log-transformed. Top 3,000, or 2,000 highly variable genes for the endometrial, or pelvic cell datasets, respectively, were identified by Scanpy. The mitochondrial genes, haemoglobin genes, and ribosomal genes were excluded from the highly variable gene set [43]. Principal component analysis (PCA) was performed using the highly variable gene set. After Harmony (v.0.0.10) [44] batch effect correction, harmony-corrected components were used to generate the neighborhood graph and perform dimensionality reduction using uniform manifold approximation and projection (UMAP). The Leiden algorithm was then used for clustering analysis [45]. Marker genes of each cluster were identified using the Wilcoxon rank-sum test [46] with the default setting (**Supplementary Table S4**). Major cell types were determined by manually annotating cell clusters with previously established marker genes from the literature [15, 39, 47, 48].

#### 2.2.4. Identification of Enriched Genes and Gene Ontology Analysis in FOLR2^+^ Endometrial and Pelvic Cells

To understand the transcriptional profiles of folate receptor beta-positive (FOLR2^+^) macrophages, cells with specific FOLR2 expression (FOLR2>0) were labeled and extracted from endometrial and pelvic cell matrices. Enriched genes in FOLR2^+^ cells were identified by the Wilcoxon rank-sum test in a one-versus-rest fashion, with a minimum log_2_(fold change) of 2 (**Supplementary Table S5**). Gene Ontology (GO) was performed on the identified enriched genes in FOLR2^+^ endometrial or pelvic cells using clusterProfiler (v.4.2.2) [49]. The enriched GO list was further filtered with the *simplify* function (cutoff = 0.7) to remove redundant terms (**Supplementary Table S6**). Differentially expressed genes (DEGs) among eutopic endometrium without (Control; Cont) or with endometriosis (EuE), and ectopic endometrium (EcE) in the endometrial FOLR2^+^ macrophages, and DEGs between control (Cont) and endometriotic (Endo) pelvic exudate cells, were identified by Wilcoxon rank-sum test, with a minimum log_2_(fold change) of 0.5 (**Supplementary Table S7**).

### 2.3. Chemistry Experimental Section

Experimental methods related to materials, instrumentation, synthesis protocols, characterization of compounds (**Supplementary Figures S1-S15 and S17-S34**), in vitro drug release studies, and FA-2DG-D-Niclo formulation stability studies are presented in the **Supplementary Information**.

### 2.4. Animals

C57BL/6 mice were purchased from Inotiv and housed in an environment-controlled animal facility (20-22 °C, ∼40% relative humidity, and 12:12 light-dark cycle) with *ad libitum* access to food and water. All animal experiments were conducted at Washington State University in accordance with NIH guidelines for care and use of laboratory animals (protocol #6751).

### 2.5. Mouse Model of Endometriosis

An experimental mouse model of endometriosis was employed, using procedures previously described and adopted by us, with minor modifications [50]. Briefly, to induce endometriosis-like lesions (ELL), female donor mice were injected subcutaneously with pregnant mare serum gonadotropin (PMSG; 5 IU; Sigma, St. Louis, MO) to stimulate an estrogenic response within the uterus. Uteri were harvested from these donors 41 hours after PMSG injection. The endometrium was then scraped from myometrium and dissected into fragments (1-2 mm per side), and introduced as the source of syngeneic mouse endometrium (50 mg of fragments) via injection (in 0.2 mL of PBS) into the peritoneal cavity in the ovary-intact recipient females under anesthesia via inhaled isoflurane.

### 2.6. Animal Study Design

#### 2.6.1. In Vivo Dendrimer Biodistribution

Adult female mice (n=36, ∼3 months old) were randomly divided into three groups (n=3/group/timepoint): 1) sham controls administered with Cy5-labeled dendrimer FA-2DG-D-Cy5; 2) ELL-induced mice administered with 2DG-D-Cy5; 3) ELL-induced mice administered with FA-2DG-D-Cy5. The animals in the ELL groups received donor endometrium to induce endometriotic lesions, and the sham control mice received PBS. One week following lesion induction, all mice received FA-2DG-D-Cy5 or 2DG-D-Cy5 injected into the tail vein, as listed in the above group description, at a dose of 50 mg/kg (300 μL). Mice were then euthanized at 1, 6, 24, and 72 hours after injection. The whole skin was gently peeled off to acquire the whole-body Cy5 fluorescence distribution using an in vivo imaging system (IVIS; Revvity). Lesions and/or major organs, including the brain, heart, spleen, lungs, liver and kidneys, were dissected to obtain ex vivo fluorescence images and stored at -80 °C. The fluorescence intensity of the region of interest (ROI) was calculated with the Living Image software (4.7.4) accompanying IVIS. Amounts of dendrimer in major organs and lesions were further examined ex vivo using fluorescence intensity-based quantification.

#### 2.6.2. Ex Vivo Dendrimer Quantification

Frozen organs (lesions, liver, spleen, heart, kidneys, brain and lungs) were slowly thawed on ice and weighed. Tissue samples were then dissected to obtain known masses from each organ. Samples were homogenized in methanol (1 mL per 100 mg tissue) with stainless steel beads using a tissue homogenizer. The homogenized samples were centrifuged at 4°C, and the clear supernatants were transferred to protein LoBind Eppendorf tubes and stored at -80°C. For fluorescence quantification, thawed supernatants were centrifuged again before analysis. Fluorescence intensity was measured on a Varioskan Lux Multimode Microplate Reader (ThermoFisher), using λ_ex = 650 nm and λ_em = 670 nm for Cy5. Background fluorescence from control solvent was subtracted, and corrected fluorescence values were converted to 2DG-D-Cy5 and FA-2DG-D-Cy5 concentrations using calibration curves generated at 5-10 slit widths.

#### 2.6.3. In Vivo Cellular Uptake of Dendrimer

To evaluate the specificity of cellular uptake of the dendrimer, endometriotic lesions were induced in 6 female mice. On Day 7 following lesion induction, mice received an intraperitoneal (i.p.) injection of FA-2DG-D-Cy5 or 2DG-D-Cy5 (50 mg/kg/bw, n=3/group). Peritoneal cells, lesions, kidney, liver and spleen were collected at 2 weeks after dendrimer injection. Specific cellular uptake of the dendrimer was then assessed by immunostaining.

#### 2.6.4. In Vivo Efficacy of Dendrimer-Niclosamide

The efficacy of targeted delivery of niclosamide with FA-conjugated dendrimer (FA-2DG-D-Niclo) was examined using the mouse model of endometriosis described above. On Day 7, ELL-induced mice were administered a single i.p. injection of control (FA-2DG-D), or 25 mg or 50 mg/kg body weight (bw) of FA-2DG-D-Niclo. Behavioral tests were performed on Day -1 (a day before lesion induction), Day 7 (before dosing), and Day 14 or 21 (1 or 2 weeks after dosing, n=16 at Day -1 and Day 7, n=8 at Day 14 and Day 21). Mice were then euthanized on Day 14 or 21 after the last behavioral test. Peritoneal fluid was recovered by lavage (4 mL x 2 of ice-cold PBS with 3% FBS) for immune profiling. The number and volume of lesions were recorded. Lesions were fixed in 4% paraformaldehyde for further analysis.

### 2.7. von Frey Test

A standard behavioral (mechanical sensitivity) test was performed before sample collection, as described by our laboratory previously [51, 52]. Mice (n=8/group/time point) were allowed to acclimate quietly in the testing room for 30 minutes under red light. Then, von Frey filaments (BIO-VF-M, Bioseb) were applied 10 times to the skin perpendicular to the lower abdomen and bilateral hind paws to evaluate local or systemic sensitivity, respectively. The force in grams (g) of the filament evoking a withdrawal response (50% response count as sensitive) was recorded. Three behaviors were considered positive responses to filament stimulation: 1) sharp retraction of the abdomen, 2) immediate licking and/or scratching of the area of filament stimulation, or 3) jumping. All behavioral tests were performed with investigators blinded to treatment groups. These data were then subsequently analyzed by another blinded investigator.

### 2.8. Flow Cytometry

Single-cell suspensions of peritoneal exudate cells were used for immune cell profiling by flow cytometry as described previously [51, 52]. Briefly, peritoneal exudate cells were lysed using Red Blood Cell Lysis Buffer (BioLegend) and incubated at room temperature for 20 minutes with Zombie Aqua™ Fixable Viability dye (BioLegend). Cells were blocked on ice for 20 minutes with Fc Block anti-CD16/CD32 (ThermoFisher) and stained with fluorochrome-conjugated monoclonal antibodies for 1 hour (**Supplementary Table S8**). Samples (n=8/group) were acquired with the Attune NxT Acoustic Focusing Cytometer using Attune NxT software (ThermoFisher), and data were analyzed with FlowJo v10.4 software (FLOWJO).

### 2.9. Immunohistochemistry and Confocal Imaging

The peritoneal exudate cell suspension was first fixed with 4% paraformaldehyde at RT for 15 min, and blocked with 10% donkey serum at RT for 30 min. Cells were then incubated with FRβ primary antibody (1:200, Abcam, Cat.#ab302532) at 4°C overnight and AlexaFluor 488-conjugated F(ab’) secondary antibody (Molecular Probe) at RT for 1 hour. Following DAPI staining, the cells were resuspended with Fluoromount-G (SouthernBiotech) and mounted on slides. The mounted slides were imaged using a Leica SP8-X Confocal Microscope.

Immunohistochemistry of FRβ on cross-sections (5 µm) of paraffin-embedded tissues, including lesion, kidney, and liver, was performed using the FRβ primary antibody (1:400) and AlexaFluor 488-conjugated F(ab’) secondary antibody, or VECTASTAIN ABC kit (Vector Lab). Immunostaining images were acquired by Leica SP8-X Confocal Microscope or Leica DM4 B microscopy. FRβ-positive cells in lesions were counted by Image J in an area of 0.07244 mm^2^, and the number of FRβ^+^ cells per 1 mm^2^ was shown.

### 2.10. Statistical Analysis

Statistical analyses were performed using GraphPad Prism (version 9.5). To compare differences in dendrimer organ uptake among groups, one-way ANOVA followed by the Tukey multiple comparison tests was used. To analyze behavioral test results, time-dependent differences within a single group were compared using one-way ANOVA followed by the Dunnett’s multiple-comparison test. To compare effects of different niclosamide dosages of niclosamide at each time point, one-way ANOVA was used with Tukey multiple comparison tests. Differences in lesion number, volume, and FRβ⁺ or CD45⁺ cell counts between groups at each time point were analyzed using one-way ANOVA followed by the Tukey multiple-comparison test. In all cases, data were tested for normal distribution using the Shapiro-Wilk normality test. If data were normally distributed, one-way ANOVA was used to analyze differences among groups. If data were not normally distributed, the Kruskal-Wallis test was performed, as stated in the figure caption. Significance was defined as p < 0.05 (*), p < 0.01 (**), p < 0.001 (***), or p < 0.0001 (****).

## 3. Results and Discussion

### 3.1. FRβ-expressing Macrophages in Eutopic and Ectopic Endometrium, and Pelvic Exudate Cells in Endometriosis

We performed an integrated analysis with our newly generated pelvic exudate cell scRNA-seq dataset from women with endometriosis (**Supplementary Tables S1 and S2**), along with publicly available human datasets, including eutopic endometrium without or with endometriosis, ectopic endometrium (lesion), and pelvic exudate cells (**Figure 1A-B, Supplementary Table S3**). After data merging and quality control filtering, 748,509, and 76,187 total high-quality endometrial and pelvic exudate cells were retained, respectively. Clustering analysis identified 7 major cell types in the endometrium (EM), including B lymphocytes (CD79A^+^), endothelial cells (PECAM1^+^), epithelial cells (EPCAM^+^), myeloid cells (LYZ^+^), perivascular cells (RGS5^+^), stromal cells (DCN^+^), and T lymphocytes and natural killer (NK) cells (CD3E^+^ and CD2^+^) (**Figure 1C and E, Supplementary Table S4**). In the pelvic cavity (P), B lymphocytes (CD79A^+^), dendritic cells (CD1C^+^), macrophages (Mac, CD14^+^), mast cells (TPSAB1^+^), NK cells (GNLY^+^), and T lymphocytes (CD3D^+^) were assigned as overarching cell types (**Figure 1D and F, Supplementary Table S4**). As expected, UMAP projection confirmed that FOLR2^+^ cells are localized within the myeloid cell clusters of endometrial and pelvic exudate cells (**Figure 1G-H**). Gene Ontology (GO) enrichment analysis indicated that both endometrial and pelvic FOLR2^+^ macrophages were primarily associated with myeloid migration and activation and immune regulation (**Figure 1I-J, Supplementary Table S6**). In particular, biological processes associated with cytokine production, leukocyte cell-cell adhesion, and interleukin-6 (IL-6) production were predominantly enriched in the FOLR2^+^ macrophages of the endometrium. Stronger association with phagocytosis, cholesterol regulation, and response to bacteria was detected in the pelvic FOLR2^+^ macrophages, consistent with the primary roles of residential FOLR2^+^ macrophages in host defense and metabolic processes [53–55]. Comparative analysis showed a significantly higher abundance of FOLR2^+^ macrophages in ectopic endometrium (EcE) than in eutopic endometrium with (EuE) and without (Cont) endometriosis (**Figure 1K**). Similarly, in pelvic exudate cells, significantly elevated numbers of FOLR2^+^ macrophages were observed in patients with endometriosis (**Figure 1L**). Furthermore, DEG analysis of endometrial and pelvic FOLR2^+^ macrophages demonstrated that significant upregulation of proinflammatory (CCL2, IL-6R, and NFKB1), angiogenesis (FGFR1, HIF1A, and MMP2), and adhesion-related genes (ITGAV, TGFBR1, and TGFBR2) in endometriosis-associated endometrium (EuE and/or EcE) and pelvic cells (Endo) (**Figure 1M-N, Supplementary Table S7**). Collectively, these results indicate an expansion of a pro-disease FOLR2^+^ macrophage subpopulation in the ectopic endometrium and pelvic cavity in endometriosis patients with pro-disease functions.

### 3.2. Synthesis and Characterization of FA-2DG-D Using Click Chemistry

A major challenge in developing nanomaterial-based drug delivery systems lies in the complexity and limited reproducibility of synthetic procedures, scalability constraints, and poor batch-to-batch consistency. To overcome these limitations, we used efficient copper(I) catalyzed alkyne-azide cycloaddition (CuAAC) click reaction to design and synthesize the FA-2DG-D conjugate capable of selectively targeting FRβ⁺ macrophages for treating endometriosis. This click chemistry-based approach offers a robust and versatile strategy for achieving precise, reproducible, and site-specific conjugation of targeting ligands or therapeutic molecules to polymeric nanocarriers, including dendrimers [28, 56–58].

Folic acid (**1**) was functionalized with an azide group to enable its participation in the CuAAC reaction. The folate receptor recognizes folic acid through its pterin and *p*-aminobenzoate moieties via multiple hydrogen bonds and electrostatic interactions within the receptor pocket [59]. To preserve these recognition elements, conjugation with the azide linker was performed through the α-carboxylic acid group of the glutamic acid residue, forming a stable amide linkage with the terminal amine of the linker. The carboxylic acid in the ligand was activated using dicyclohexylcarbodiimide (DCC) and *N*-Hydroxysuccinimide (NHS), and the resulting NHS-activated intermediate was reacted with azido-PEG5-amine (**2**) in the presence of DIPEA to yield the desired FA-Azide (**3**). Successful formation of the azide-functionalized folic acid was confirmed by ^1^H NMR spectroscopy, which showed characteristic PEG proton signals from the azide linker at δ 3.0 (2H), 3.4 (2H), and 3.5-3.7 (20H) ppm, along with a new amide proton signal at δ 7.3 ppm (**Figures 2B and Supplementary Figure S1**). HPLC analysis confirmed the purity of the product, exhibiting a single peak with a retention time of 13.3 minutes with more than 98% purity (**Supplementary Figure S3**). To enable the subsequent CuAAC coupling, terminal alkyne groups were introduced on the surface of 2DG-D. The primary hydroxyl groups of the surface-exposed 2DG moieties were utilized for this modification. 2DG-D (**4**) was reacted with 5-hexynoic acid (**5**) in the presence of 1-ethyl-3-(3-dimethylaminopropyl)carbodiimide hydrochloride (EDC) and 4-(dimethylamino)pyridine (DMAP) to afford 2DG-D-Hexyne (**6**) (**Figure 2A**). ^1^H NMR spectrum showed new methylene proton signals from the hexynoic linker in the aliphatic region at δ 1.75-1.68 (8H), 2.46-2.34, and 2.74-2.58 (8H) ppm (**Figures 2B and Supplementary Figure S5**), confirming the attachment of 4 hexyne arms. HPLC analysis showed a shift in retention time from 14.0 minutes for 2DG-D (**4**) to 19.7 minutes for 2DG-D-Hexyne (**6**), confirming successful surface modification (**Supplementary Figures S7 and S15**). Finally, the CuAAC reaction between FA-Azide (**3**) and 2DG-D-Hexyne (**6**) yielded the FRβ⁺ macrophage-targeted platform FA-2DG-D (**7**). Previous studies have demonstrated that dendrimers bearing a single folic acid ligand exhibit monovalent, reversible binding with folate-binding protein (FBP), whereas dendrimers conjugated with two or more folic acid molecules engage in multivalent interactions, resulting in stronger receptor binding [60–62]. Given the high hydrophobicity of folic acid, conjugating more than 2 ligand molecules could potentially reduce the dendrimer’s aqueous solubility. Our CuAAC click chemistry approach ensured desired loading of folic acid on the dendrimer. The number of attached molecules was confirmed through the comparative integration of aromatic protons from the dendrimer at δ7.2 ppm (16H), and the aromatic and amide protons of folic acid at δ8.7 ppm (2H), 7.7 ppm (4H), 7.0 ppm (4H) and at 6.7 ppm (4H). The HPLC chromatogram showed a retention time shift from 19.7 minutes for 2DG-D-Hexyne (**6**) to 16.1 minutes for FA-2DG-D (**7**) (**Supplementary Figures S7** and **S10)**. The purity of FA-2DG-D (**7**) was determined to be more than 96% by HPLC (**Supplementary Figure S10**). Dynamic light scattering (DLS) analysis revealed a hydrodynamic diameter of 6.6 ± 0.4 nm and a zeta potential of –5.8 ± 0.3 mV (**Supplementary Figures S11 and S12**).

**Figure 2.**
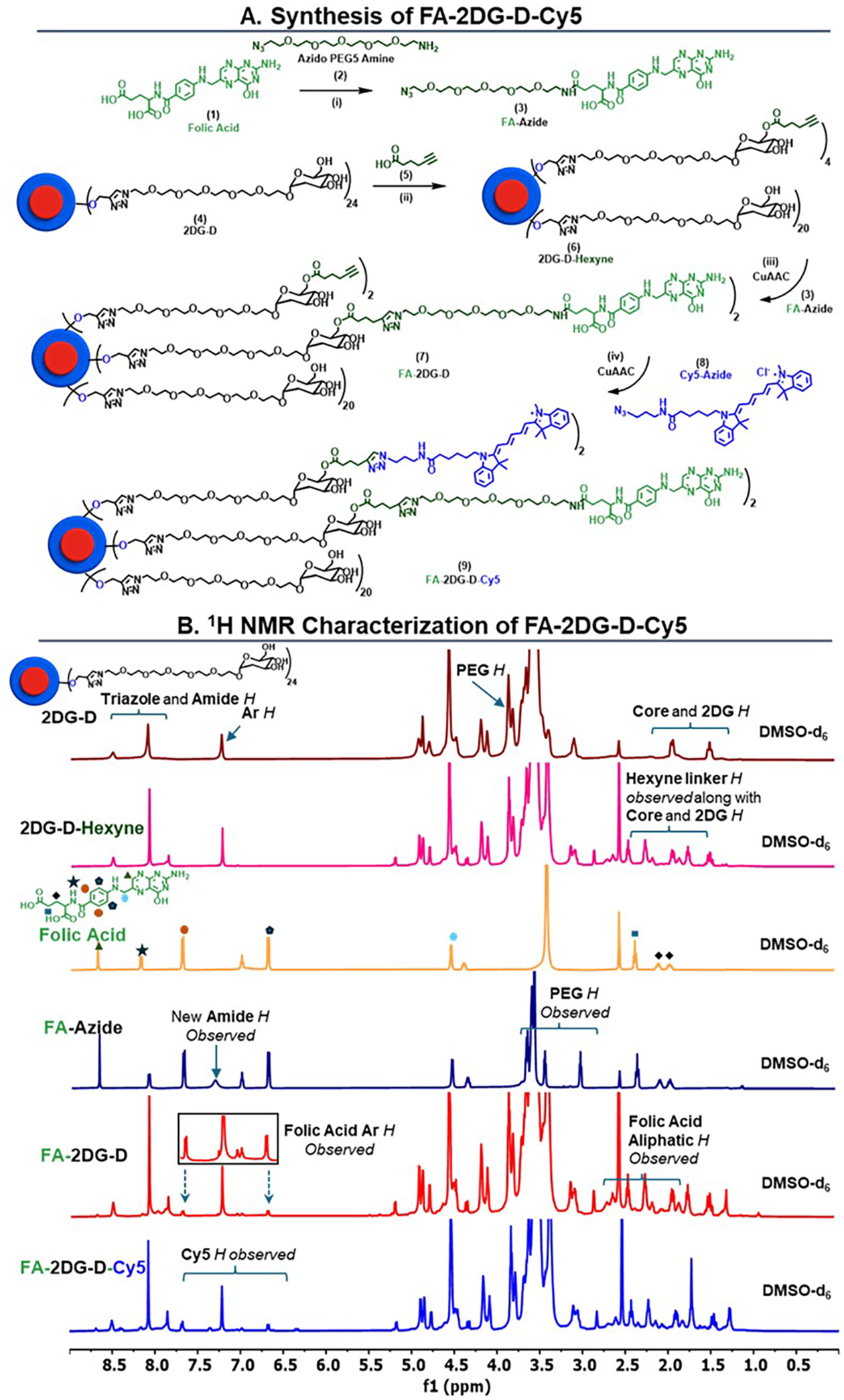
Preparation and structural elucidation of FA-2DG-D-Cy5 conjugate. (**A**) Schematic representation for the synthesis of FA-2DG-D-Cy5 using CuAAC reaction. Reagents and conditions: (i) DMSO, DCC, NHS, DIPEA, RT, 12 hours, 70% (ii) DMF, EDC.HCl, DMAP, RT, 12 hours, 80% (iii) CuSO_4_.5H_2_O, Sodium ascorbate, DMF, DI Water, 60 °C, 10 hours, 84% (iv) CuSO_4_.5H_2_O, Sodium ascorbate, DMF, DI Water, 40 °C, 6 hours, 88% (**B**) ^1^H NMR spectra showing characteristic proton signals of 2DG-D, 2DG-D-Hexyne, Folic Acid, FA-Azide, FA-2DG-D and FA-2DG-D-Cy5 at each step of the synthesis.

To enable visualization and assessment of uptake of the dendrimer in FRβ⁺ macrophages in endometriosis, the dendrimer was further functionalized with a near-infrared dye, cyanine 5 (Cy5). The FA-2DG-D **(7)** containing approximately two terminal alkynes on its surface was conjugated with Cy5-azide (**8**) via a CuAAC reaction, yielding the fluorescently labeled FA-2DG-D-Cy5 (**9**). Successful conjugation was confirmed by the appearance of characteristic Cy5 proton signals in the ^1^H NMR spectrum (**Figure 2B**). The number of Cy5 molecules attached to the dendrimer surface was estimated by proton integration. Comparison of the dendrimer aromatic protons at δ 7.2 ppm (16H) with the Cy5 aromatic protons at δ 8.4 (4H), 8.2 (4H), 7.4 (4H), 6.6 (2H), and 6.4 ppm (2H) indicated the attachment of approximately two Cy5 molecules (**Figures 2B and Supplementary Figure S13**). The HPLC chromatogram of FA-2DG-D-Cy5 (**9**) exhibited a retention time shift from 16.1 to 17.7 minutes following Cy5 conjugation, with a purity exceeding 97% (**Supplementary Figures S14** and **S15**). In addition, the Cy5-labeled dendrimer displayed a characteristic UV absorbance at 650 nm, corresponding to Cy5, which further confirmed successful conjugation. A Cy5-labeled 2DG-D dendrimer (2DG-D-Cy5), lacking the folic acid targeting ligand, was prepared as a non-targeted control for uptake studies following our previously reported procedure [27].

### 3.3. Fluorescence Imaging and Quantitative Analysis of the Biodistribution of Systemically Administered 2DG-D-Cy5 and FA-2DG-D-Cy5 in a Mouse Model of Endometriosis

We first evaluated the selective FRβ^+^ macrophages targeting capability of FA-2DG-D-Cy5 in ELL-induced mice at 1, 6, 24, and 72 hours (n=3) and compared it with sham controls (n=3). To evaluate the contribution of folic acid (FA) toward lesion targeting, biodistribution studies of non-targeted 2DG-D-Cy5 in ELL-induced mice (n=3) were also performed under identical conditions (**Figure 3**). One week after lesion induction, 50 mg/kg bw of fluorescently labeled (FA-2DG-D-Cy5) or non-targeted (2DG-D-Cy5) was injected into the tail vein. Cy5 fluorescent signals were calculated via the IVIS system at 1, 6, 24, and 72 hours. FRβ^+^ macrophages in the lesions showed a pronounced elevation in fluorescent (Cy5) signal starting at 1 hour post-injection (green arrows), which persisted above that of the 2DG-D-Cy5 group over the 72-hour observation period (**Figure 3A**). The ex vivo fluorescent results at 72 hours showed specific uptake of FA-2DG-D-Cy5 in the lesion, although some nonspecific uptake of FA-2DG-D-Cy5 and 2DG-D-Cy5 was observed primarily in the liver and kidneys (**Figure 3B-C**). For more precise quantification, frozen tissue samples from the 1, 6, 24, and 72-hour groups were homogenized, and fluorescence intensity was measured to assess quantitative dendrimer uptake in lesions and major organs. Both FA-2DG-D-Cy5 and 2DG-D-Cy5 showed strong, persistent accumulation within endometriosis lesions beginning at 1-hour post-injection and sustained through 72 hours **(Figures 3D and S16)**. By 72 hours, lesion uptake of FA-2DG-D-Cy5 was significantly higher than that of 2DG-D-Cy5, suggesting its targeted accumulation in FRβ^+^ macrophages that are abundant in endometriotic lesions (**Figure 3D-E and Supplementary Figure S16**). Moreover, FA-2DG-D-Cy5 demonstrated minimal accumulation (<5% Injected Dose (% I.D.)) in other major organs, indicating rapid clearance from off-target tissues **(Figure 3E)**. Importantly, unlike many nanoparticle systems that exhibit up to ∼80% hepatic accumulation, FA-2DG-D-Cy5 showed only ∼4% liver accumulation, comparable to the previously characterized 2DG-D-Cy5 platform, demonstrating its favorable pharmacokinetic profile [27].

**Figure 3.**
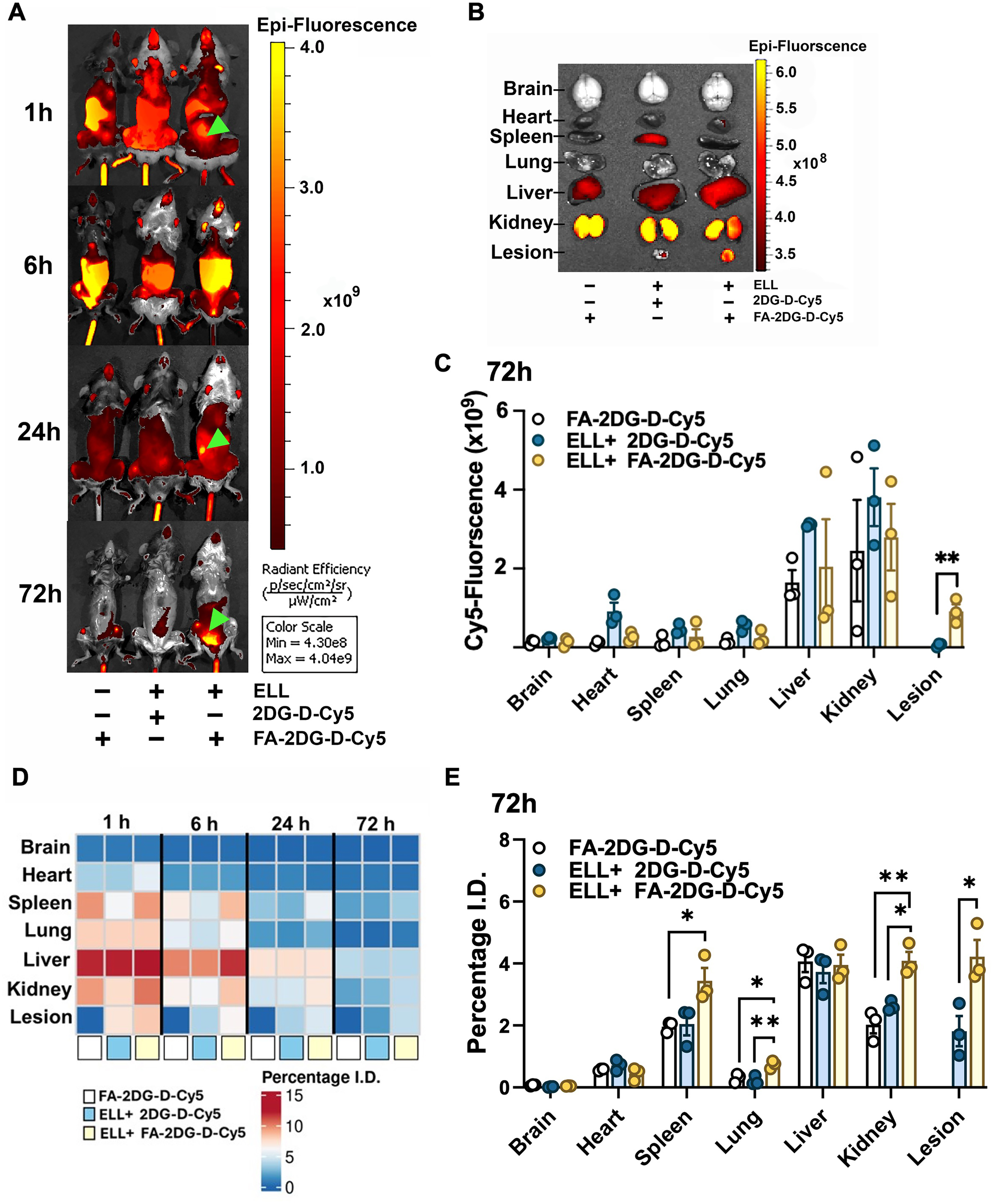
In vivo and ex vivo fluorescence imaging and whole-body biodistribution of FA-2DG-D-Cy5. (**A**) Whole-body in vivo fluorescence images of mice following intravenous (tail vein) administration of FA-2DG-D-Cy5 in sham controls (left) and ELL-induced mice (right), or 2DG-D-Cy5 administered to ELL-induced mice (center) at 1, 6, 24, and 72 hours (n = 3/group/timepoint). (**B**) Ex vivo fluorescence images of major organs collected at 72 hours post-injection. Organs are arranged from top to bottom as: brain, heart, spleen, lungs, liver, kidneys, and lesion. (**C**) Quantitative analysis of in vivo fluorescence signals in major organs at 72 hours (n = 3). (**D**) Heat map showing the mean percentage of injected dose (I.D.) in lesions and major organs at 1, 6, 24, and 72 hours after FA-2DG-D-Cy5 administration in sham controls and ELL-induced mice, or ELL-induced mice treated with 2DG-D-Cy5 (n=3/group). Heatmap was generated with the ComplexHeatmap package of R. (**E**) Quantitative biodistribution of FA-2DG-D-Cy5 and 2DG-D-Cy5 in lesions and major organs at 72 hours in sham and ELL-induced mice (n = 3). Data for (D) and (E) were obtained by fluorescence spectroscopy of homogenized tissues and are shown as the percentage of the injected dose per organ. Differences in each tissue among groups for data in (C) and (E) were analyzed by one-way ANOVA followed by the Tukey multiple-comparison test. p < 0.05 (*), p < 0.01 (**). ELL, endometriosis-like lesion.

### 3.4. Confocal Imaging of Cellular Uptake of Intraperitoneally Administered FA-2DG-D-Cy5 and 2DG-D-Cy5 in a Mouse Model of Endometriosis

We next examined the targeting ability, including selective cellular uptake, localization, and retention of FA-2DG-D-Cy5 or 2DG-D-Cy5 in ELL-induced mice 2 weeks after dendrimer administration (**Figure 4**). Confocal imaging showed that FA-2DGD-Cy5 (red) was selectively targeted and taken up by FRβ^+^ macrophages (green), not only in lesions but also in peritoneal exudate cells (**Figure 4A-B, white arrows**), where FRβ^+^ macrophages were elevated in ELL-induced mice. In contrast, cellular uptake of 2DG-D-Cy5 without FA ligand was limited in the lesion and peritoneal cells (**Figure 4A-B**). Given the potential uptake of non-targeted 2DG-D-Cy5 by activated macrophages [28], we further quantified nonspecific FA-2DG-D-Cy5 internalization in peritoneal exudate cells. The uptake was an average of 0.59%, indicating minimal nonspecific internalization. No dendrimer retention was observed in the kidneys, liver, or spleen in ELL-induced mice 2 weeks following dendrimer administration **(Figure 4C)**, indicating that FA-2DG-D-Cy5 underwent complete clearance, and off-target accumulation in major organs was negligible. These results suggest that FA-2DGD-Cy5 dendrimer is internalized only by ligand-receptor endocytosis in FRβ^+^ macrophages in the lesions and peritoneal cells. As it remains stable within cells for at least 2 weeks, it functions as an excellent nanocarrier for drug delivery.

**Figure 4.**
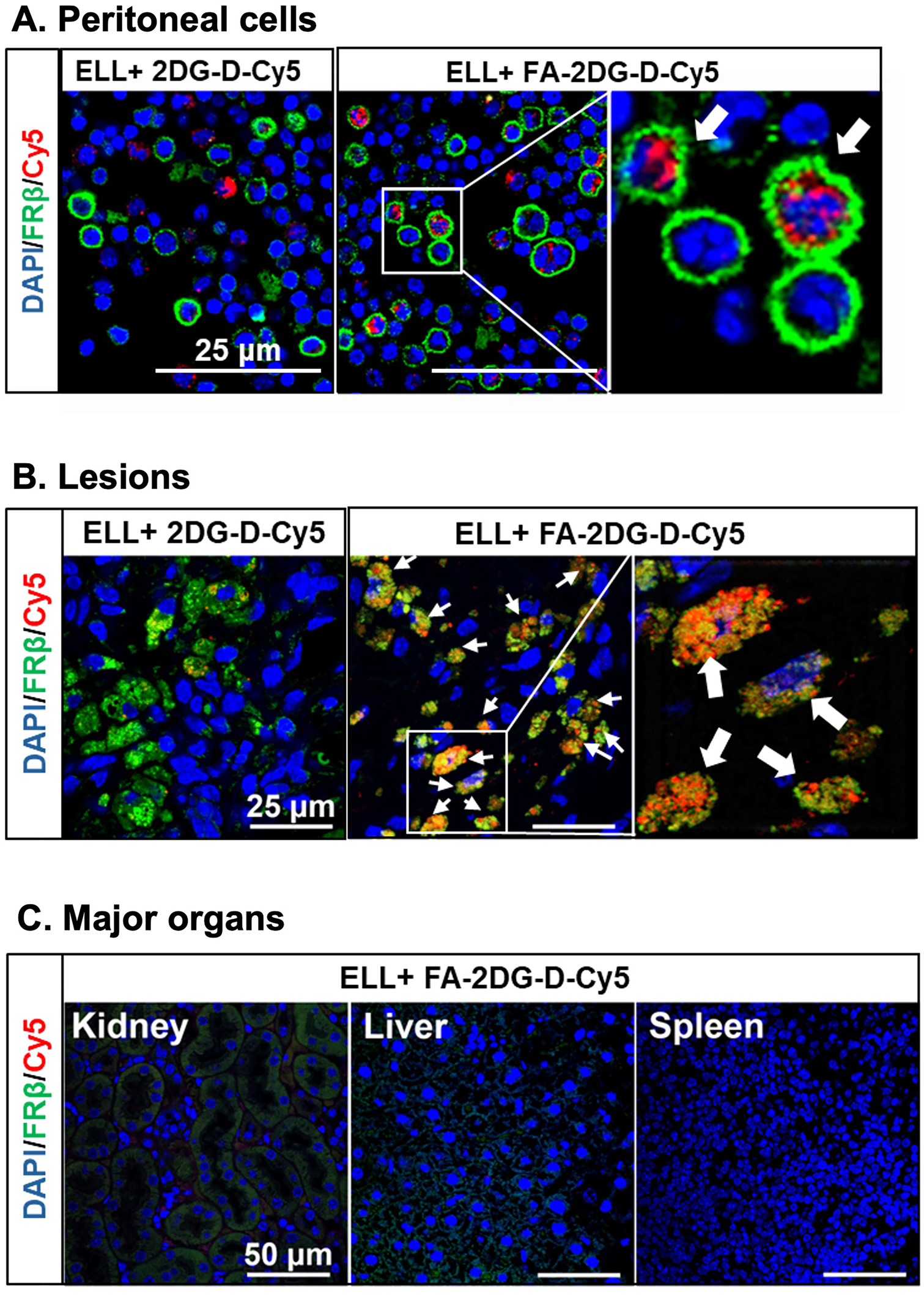
Cellular uptake of FA-2DG-D-Cy5 and 2DG-D-Cy5 at 2 weeks following intraperitoneal administration. Representative images showing selective uptake of FA-2DG-D-Cy5 or 2DG-D-Cy5 (**A**) in peritoneal cells, (**B**) in endometriosis-like lesions (ELL), and (**C**) kidney, liver, and spleen from ELL-induced mice (n=3). Peritoneal cells were co-stained with FRβ (green) and DAPI (blue). The Cy5 signal (red) was directly acquired using a confocal microscope. White arrows denote Cy5-positive signal in FRβ^+^ cells.

### 3.5. Synthesis and Characterization of FA-2DG-D-Niclo Conjugate Using Click Chemistry

We developed a niclosamide-labeled dendrimer platform, FA-2DG-D-Niclo, by conjugating niclosamide to the FA-2DG-D, to enable targeted delivery of niclosamide to disease-driving FRβ⁺ macrophages as a therapy for endometriosis. To achieve high conjugation efficiency and precise drug loading, CuAAC was employed. A niclosamide bearing an azide linker (Niclo-Azide) was first synthesized by functionalizing the phenolic -OH group of niclosamide (**Figure 5A**). Several linker architectures were explored to identify a construct that combined physiological stability with controlled intracellular release. Initially, aliphatic and PEG-based carboxylic acids were coupled to niclosamide through EDC/DMAP-mediated esterification, aiming to generate ester linkages that are labile in the intracellular conditions and stable at physiological pH. However, due to the proximity of the ester group to the aromatic ring, the resulting phenolic esters exhibited instability and underwent rapid cleavage during subsequent CuAAC reactions [63]. To circumvent this issue, a previous report on a disulfide-linker-based niclosamide prodrug was followed with some modifications to design and synthesize niclosamide-azide (niclo-azide) containing a disulfide linker [64]. It was reported that the disulfide linker remains stable under extracellular conditions but undergoes selective cleavage in the intracellular environment, thereby releasing free niclosamide [64].

**Figure 5.**
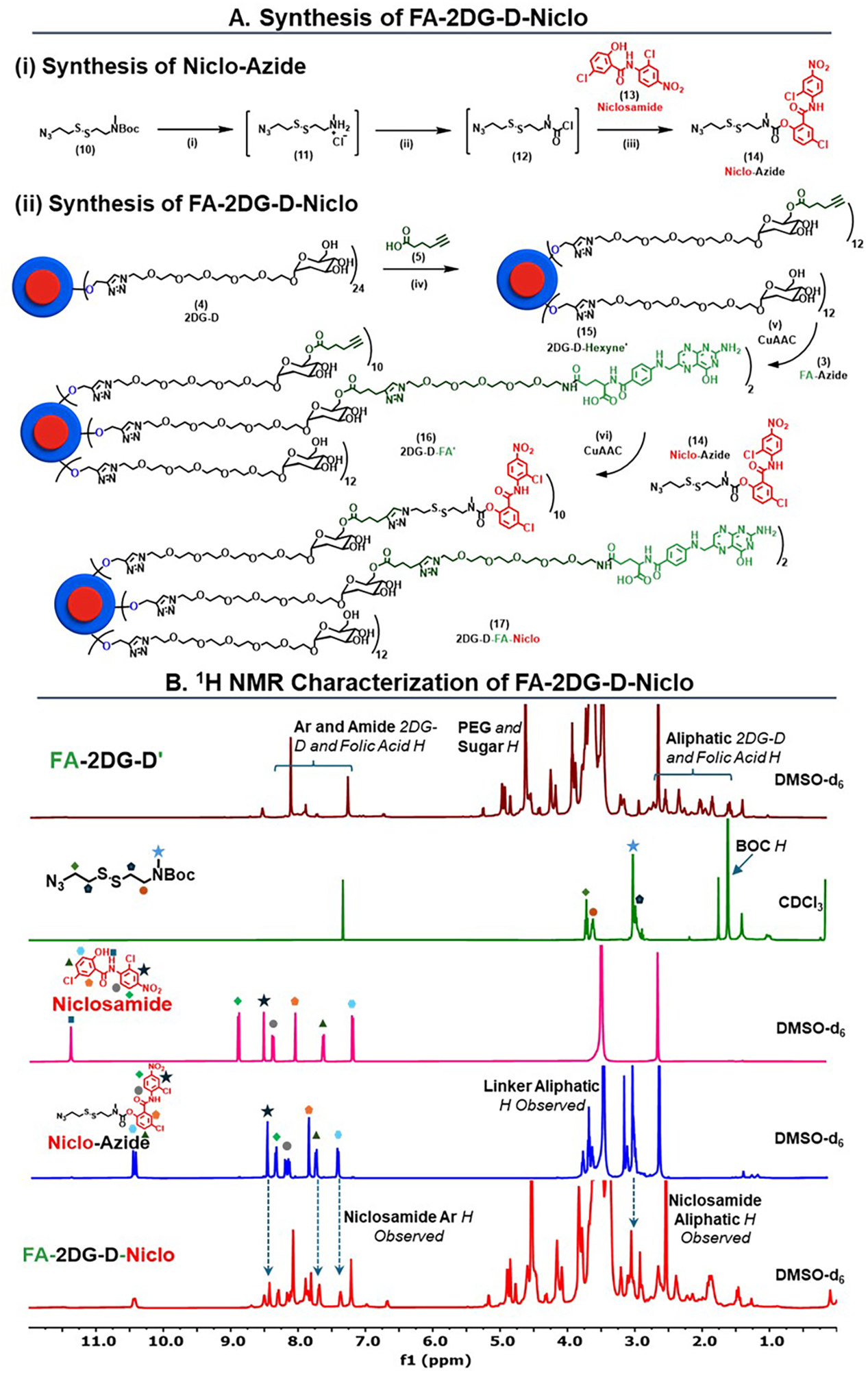
Preparation and structural elucidation of FA-2DG-D-Niclo conjugate. (**A**) Schematic representation for the synthesis of **(i)** Niclo-Azide and **(ii)** FA-2DG-D-Niclo using CuAAC reaction. Reagents and conditions: (i) DCM, HCl, Dioxane, RT, 3 hours (ii) DCM, TEA, Triphosgene, -25 °C, 1 hour (iii) DCM, TEA, DMAP, RT, 16 hours, 49% (iv) DMF, EDC.HCl, DMAP, RT, 12 hours, 82% (v) CuSO_4_.5H_2_O, Sodium ascorbate, DMF, DI Water, 60 °C, 10 hours, 85% (vi) CuSO_4_.5H_2_O, Sodium ascorbate, DMF, DI Water, 40 °C, 10 hours, 75% (**B**) ^1^H NMR spectra showing characteristic proton signals of FA-2DG-D’, Compound 10, Niclosamide, Niclo-Azide and FA-2DG-D-Niclo at each step of the synthesis.

The synthesis of niclo-azide was carried out through a multistep route (**Figure 5A**). The Boc-protected disulfide derivative **(10)** was synthesized using a published procedure [64]. It was then deprotected using 4 N HCl in dioxane to afford compound **11** as the amine hydrochloride salt, which was directly converted to the corresponding carbamoyl chloride intermediate **12** using triphosgene, following coupling with niclosamide **(13)** under mild basic conditions to afford the desired Niclo-Azide **(14) (Figure 5A)**. Successful formation of the product was confirmed by comparative integration in ^1^H NMR spectroscopy, which showed characteristic aromatic protons from the niclosamide scaffold δ 8.4-7.3 ppm (6H) and the aliphatic protons from the disulfide linker between δ 3.7-2.7 ppm (11H) **(Figures 5B and Supplementary Figure S19)**. The product purity (∼ 95.9%), was verified by HPLC, which showed a chromatogram with a retention time of 14.7 minutes **(Figures 6A and Supplementary Figure S21).** To enable subsequent CuAAC coupling, 2DG-D **(4)** was partially modified with 5-hexynoic acid. Approximately 12 hydroxyl groups per dendrimer were modified to yield 2DG-D-Hexyne’ **(15) (Figure 5A)**. The formation of the required product was confirmed by the appearance of characteristic methylene proton signals from the hexynoic linker in the aliphatic region of the ^1^H NMR spectrum at δ 1.7 (24H), 2.4 (24H) and at 2.2 ppm **(Figures 5B and Supplementary Figure S23)** which confirmed the presence of 12 hexyne arms. HPLC analysis of 2DG-D-Hexyne’ **(15)** showed the chromatogram at 10.4 minutes with a purity exceeding 97% **(Figure 6A)**. Subsequently, a CuAAC reaction between FA-Azide **(3)** and 2DG-D-Hexyne’ **(15)** yielded FA-2DG-D’ **(16)**. The ^1^H NMR spectrum of FA-2DG-D’ **(16)** showed characteristic aromatic resonances corresponding to folic acid, confirming successful conjugation of 2 molecules per dendrimer **(Supplementary Figure S26).** The resulting construct retained approximately ten free terminal alkyne groups per dendrimer. The HPLC chromatogram exhibited a retention time shift from 10.4 minutes for 2DG-D-Hexyne’ to 9.5 minutes for FA-2DG-D’ **(16)**, with a purity of ∼99% **(Figures 6A and Supplementary Figure S28)**. Lastly, FA-2DG-D’ **(16)** was conjugated with Niclo-Azide **(14)** through CuAAC to obtain FA-2DG-D-Niclo **(17)**. Consistent with our previous studies [56, 58], we aimed to maintain the drug loading between 10 and 15% to ensure aqueous solubility of the dendrimer conjugate. Drug loading was calculated by ^1^H NMR spectroscopy through comparison of the dendrimer aromatic protons at δ 7.2 ppm (16H) with the newly aromatic signals at δ 8.4 (10H), 8.3 (10H), and 8.2-7.8 ppm, along with distinct amide protons of niclosamide at δ 10.43 ppm (10H), confirmed the conjugation of approximately ten niclosamide molecules per dendrimer, corresponding to ∼12% drug loading by weight **(Figures 5B and Supplementary Figure S29)**. HPLC analysis revealed a shift in retention time from 9.5 minutes for FA-2DG-D’ **(17)** to 11.2 minutes for FA-2DG-D-Niclo **(17)**, with a final purity of more than 98% **(Supplementary Figure S31)**. DLS analysis indicated a hydrodynamic diameter of 7.6 ± 0.6 nm and a zeta potential of -7.7 ± 0.6 mV **(Figure 6C and Supplementary Figures S32 and S33)**.

**Figure 6.**
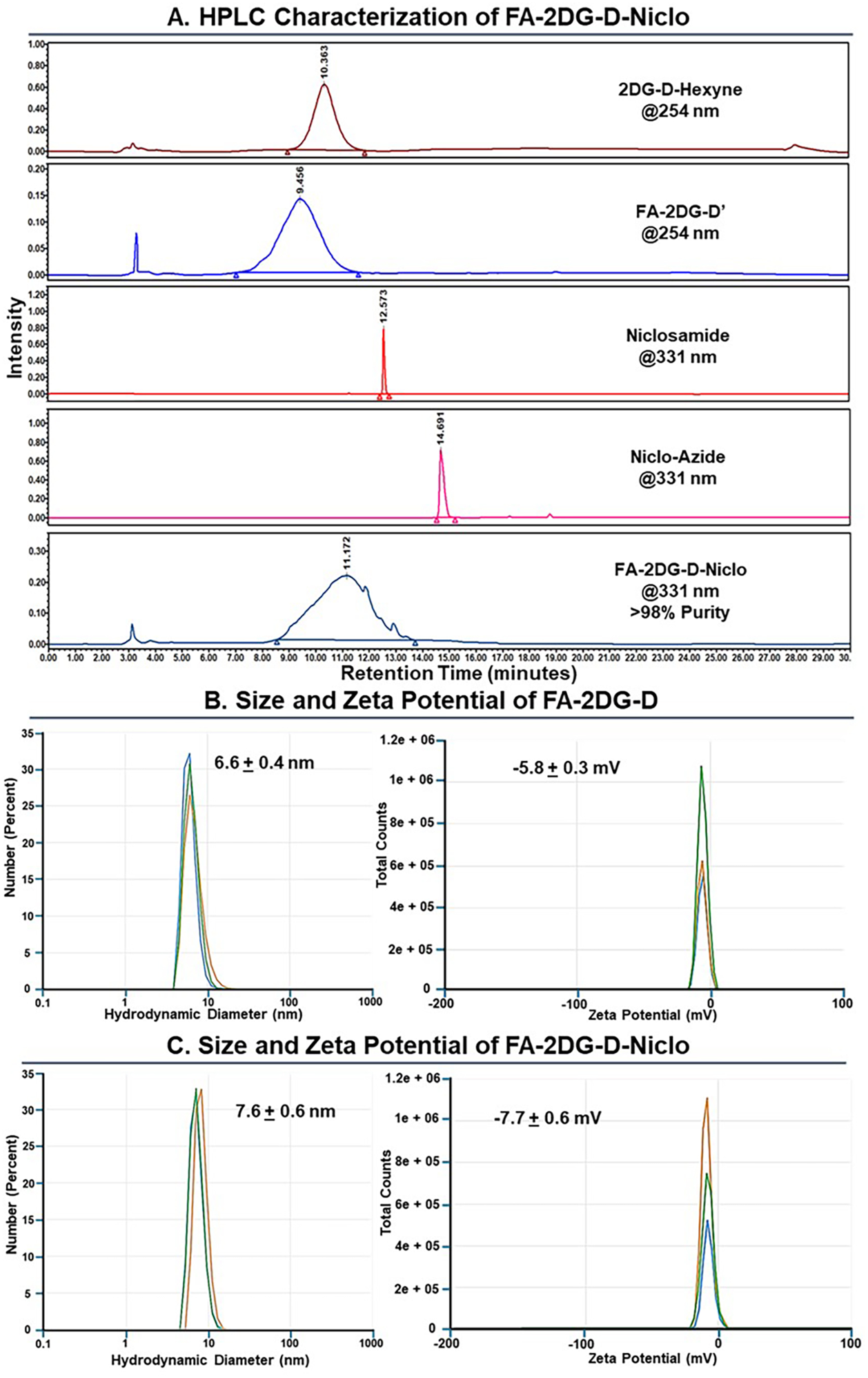
Characterization of FA-2DG-D-Niclo conjugate and intermediates: (**A**) HPLC chromatograms of intermediates at various synthesis steps and percentage purity of FA-2DG-D-Niclo. (**B**) (i) Hydrodynamic diameter, and (ii) Zeta Potential of FA-2DG-D analyzed by dynamic light scattering (DLS) in triplicate. (**C**) (i) Hydrodynamic diameter, and (ii) zeta potential of FA-2DG-D-Niclo analyzed by DLS in triplicate.

The physicochemical properties of FA-2DG-D-Niclo **(23)** are presented in **Figure 7A**. While niclosamide is practically insoluble in water (∼5 µg/mL) [65], the dendrimer-drug conjugate exhibited excellent aqueous solubility (∼150 mg/mL). At a drug loading of 12%, this corresponds to an effective niclosamide solubility of approximately 18 mg/mL in water, representing a ∼3,600 fold increase in solubility compared to free niclosamide **(Figure 7A)**.

**Figure 7.**
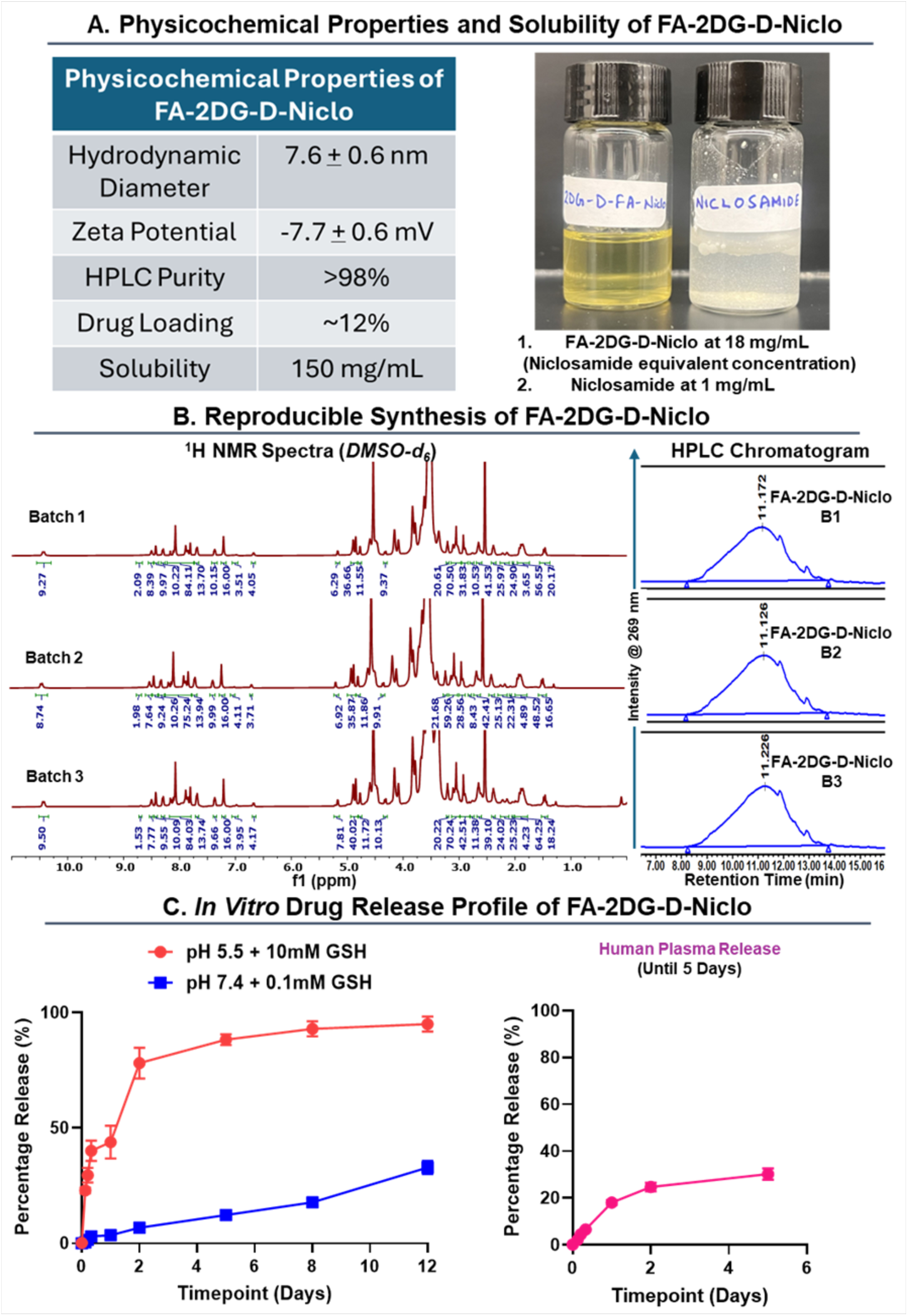
Characterization of FA-2DG-D-Niclo conjugate and intermediates: (**A**) Table showing the physicochemical properties of FA-2DG-D-Niclo and image showing the increase in solubility of niclosamide after conjugation with FA-2DG-D scaffold (FA-2DG-D-Niclo). (**B**) The figure shows the reproducibility in the synthesis of multiple 400 mg-scale batches of FA-2DG-D-Niclo via ^1^H NMR and HPLC. The ^1^H NMR spectra of 3 batches (400 mg scale) show characteristic protons from FA-2DG-D-Niclo. The HPLC chromatographs show reproducibility in the purity of these batches with all these batches showing >99% purity. (**C**) In vitro drug release profile of FA-2DG-D-Niclo conjugate in PBS and 0.1mM GSH, intracellular conditions at 37°C after 15 days, and in (100%) human plasma after 120 hours.

The shelf stability of FA-2DG-D-Niclo **(17)** (1 mg/mL) was assessed in PBS at 4 °C, room temperature (25 °C), and 40 °C over a 30-day period. The formulation remained stable throughout the study, retaining more than 96% purity by HPLC at all tested temperatures **(Supplementary Figure S34)**. No shifts in retention time were observed over 30 days, confirming the chemical stability of the formulation under these conditions.

### 3.6. Scalable and Reproducible Synthesis of FA-2DG-D-Niclo

Ensuring synthetic reproducibility and scalability remains a central barrier in advancing nanoparticle-based therapeutics toward clinical applications. To overcome these challenges, we established a streamlined and scalable synthetic workflow for FA-2DG-D-Niclo **(17)** that enables rigorous characterization of intermediates at each stage. To assess batch-to-batch consistency, three independent 400 mg-scale batches were synthesized and evaluated by ^1^H NMR and HPLC. The ^1^H NMR spectra of all three batches **(Figure 7B, left)** showed identical chemical profiles and consistent drug and ligand loading. Similarly, HPLC analysis revealed overlapping chromatogram peaks with retention times of ∼11.2 minutes and purities exceeding 98%, confirming robust batch-to-batch reproducibility **(Figure 7B, right)**.

### 3.7. In vitro drug release studies for FA-2DG-D-Niclo

We further assessed the in vitro drug-release profile of the conjugate under conditions mimicking both the extracellular environment (PBS + 0.1 mM GSH and 100% human plasma) and the intracellular microenvironment (pH 5.5 citrate buffer + 10mM GSH) **(Figure 7C)**. Since niclosamide is attached to the dendrimer through a disulfide linkage, it enables glutathione (GSH) responsive cleavage [66]. Upon cleavage, the generated free thiol is known to undergo an intramolecular nucleophilic attack on the tertiary carbamate, ultimately yielding free niclosamide [64]. When conjugated to the targeted FA-2DG-D, this design facilitates controlled release within the intracellular environments while limiting premature systemic exposure outside the lesion sites. In PBS and 0.1mM GSH, the conjugate was found to be relatively stable. Less than 40% of the drug was observed to be released in around 2 weeks, with <10% drug release detected for the first 5 days. Notably less than 5% of the drug was released within 8 hours. When incubated in human plasma, less than 35% of the drug was released over 5 days, indicating stability in physiological conditions **(Figure 7C).** Under intracellular conditions (pH 5.5 citrate buffer + 10mM GSH), the conjugate showed a sustained release profile, with ∼45% released in the first 24 hours, ∼80% by two days, and reaching ∼95% over 12 days **(Figure 7C)**. The targeted and sustained intracellular drug release profile is important to maintain the efficacy of the drug for a long time and to avoid off-target effects.

### 3.8. FA-2DG-D-Niclo attenuates lesion progression and endometriosis-associated hyperalgesia

Lastly, we assessed the efficacy of FA-2DG-D-Niclo in endometriosis disease progression and endometriosis-associated hyperalgesia. After the lesion induction, a single dose of FA-2DG-D-Niclo dendrimer (25 or 50 mg/kg bw of niclosamide) or FA-2DG-D without niclosamide control was administered i.p. in ELL-induced mice at Day 7, as shown in **Figure 8A**. Then, the efficacy of dendrimers was examined 7 (Day 14) and 14 (Day 21) days after dendrimer administration. We first confirmed the suppressing capability of FRβ^+^ macrophages in the lesions and peritoneal cavity by FA-2DG-D-Niclo dendrimer (**Figure 8B-C**). Single dosing of FA-2DG-D-Niclo (50 mg/kg bw) significantly reduced FRβ^+^ macrophages in the lesions at Day 21, although the reduction of FRβ^+^ macrophages was not observed at Day 14 (**Figure 8B**). Peritoneal FRβ^+^ macrophage population was reduced by both doses (25 and 50 mg/kg bw) of FA-2DG-D-Niclo (**Figure 8C**). The results indicate that 7 days after dendrimer administration may not be sufficient to reduce FRβ^+^ macrophages in both lesions and the peritoneum. Indeed, lesion number and volume were also significantly reduced at Day 21 by FA-2DG-D-Niclo at both doses, but there were no significant differences in lesion number or volume at Day 14 (**Figure 9A**). However, we realized that three mice treated with both doses and examined at Day 14 did not have lesions. Furthermore, four and five mice, at doses of 25 and 50 mg/kg bw of niclosamide, respectively, did not have lesions by Day 21, either. Thus, a single dose of FA-2DG-D-Niclo administration completely inhibited lesion growth and progression in 37.5-62.5% of mice, whereas all control mice developed lesions on Days 14 and 21. While reduced FRβ^+^ macrophages in the remaining lesions were observed only at Day 21, a few mice at Day 14 lacked lesions for examination, suggesting that FA-2DG-D-Niclo was likely effective for lesion growth 7 days after administration. In support of this implication, abdominal and hindpaw hypersensitivities were already improved at Day 14 (**Figure 9B**). The abdominal and hindpaw sensitivity 7 days after lesion induction was high compared to before lesion induction (Day -1). At Day 14, FA-2DG-D-Niclo at a dose of 50 mg/kg bw of niclosamide significantly reduced the abdominal sensitivity, which is considered indicative of peripheral visceral pain. Both doses of FA-2DG-D-Niclo improved hindpaw sensitivity, which is likely be affected by peripheral and central sensitization. By Day 21, FA-2DG-D-Niclo at both doses further improved abdominal and hindpaw sensitivities. The results clearly showed that a single dose of FA-2DG-D-Niclo inhibited disease progression and endometriosis-associated hyperalgesia. Notably, a single dose of FA-2DG-D-Niclo remained effective for 2 weeks after administration. The current FDA-approved niclosamide treatment requires a high dose (2g/day) due to its low water solubility and is limited to oral administration for 7 days. Niclosamide treatment for patients who have endometriosis would need to be longer due to the chronic nature of the disease. Long-term systemic treatment with niclosamide could cause off-target actions. Therefore, the 2DG-D-based delivery of niclosamide and targeting of disease-specific FRβ^+^ macrophages is highly translational, as FA-2DG-D can deliver niclosamide intracellularly into FRβ^+^ macrophages, thereby enhancing efficacy, reducing side effects, and treating chronic diseases such as endometriosis.

**Figure 8.**
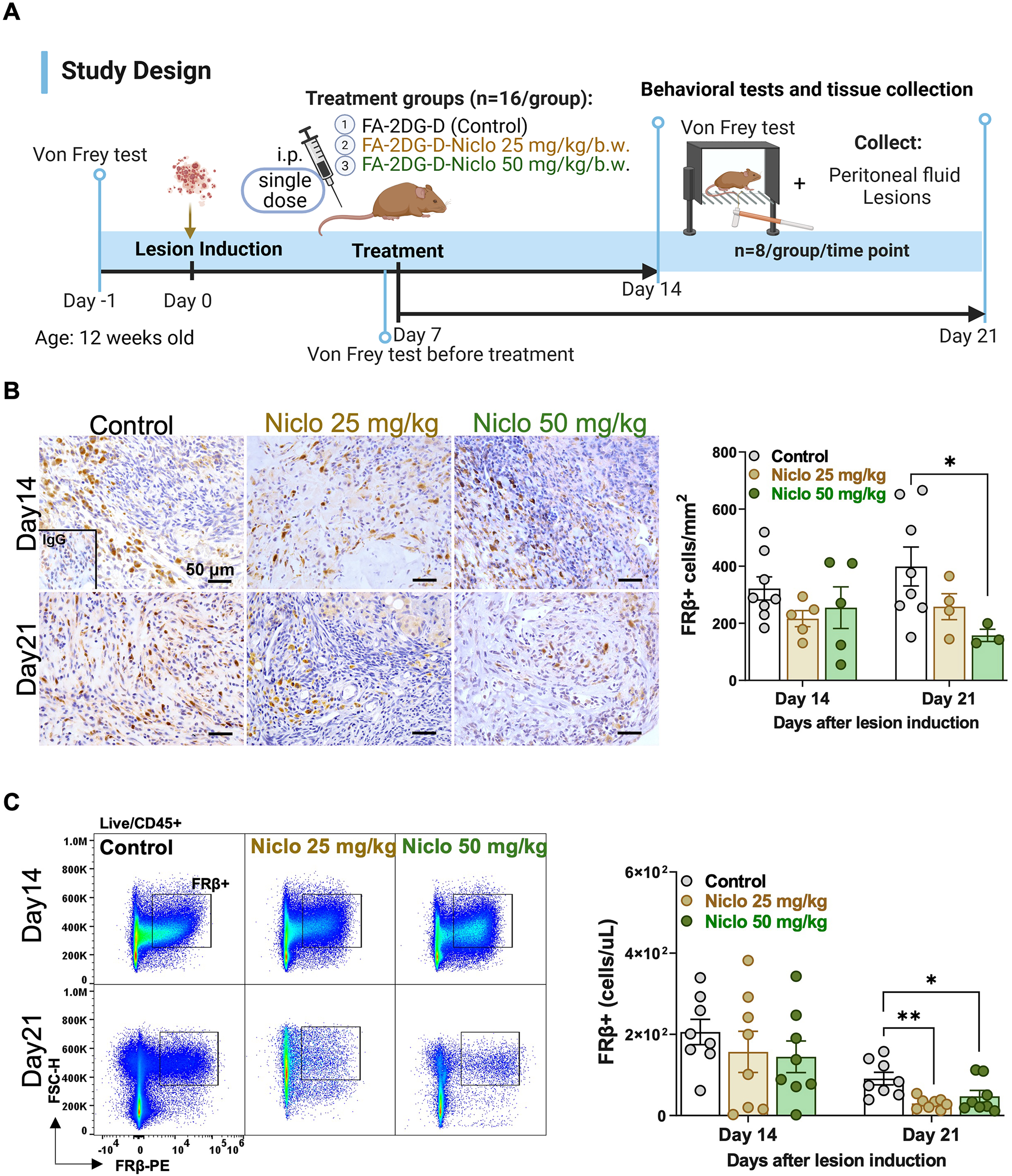
Quantitative analysis of FRβ^+^ cells in the lesion and peritoneum following dendrimer treatments. (**A**) Study design as described in Methods. (**B**) Representative immunohistochemical images (left) and quantification of FRβ^+^ cells (right) in lesions at 14 or 21 days after lesion induction (n=3-8, as dendrimer-Niclo inhibited lesion development). One-way ANOVA followed by the Tukey multiple-comparison test was used to assess differences among groups. **(C)** Representative flow plots (left) and quantification of peritoneal FRβ^+^ cells (right). Differences in peritoneal FRβ^+^ cells among groups at each time point were assessed by the Kruskal-Wallis test. Data in all figures are shown as the mean ± SEM (n=8). p < 0.05 (*), or p < 0.01 (**).

**Figure 9.**
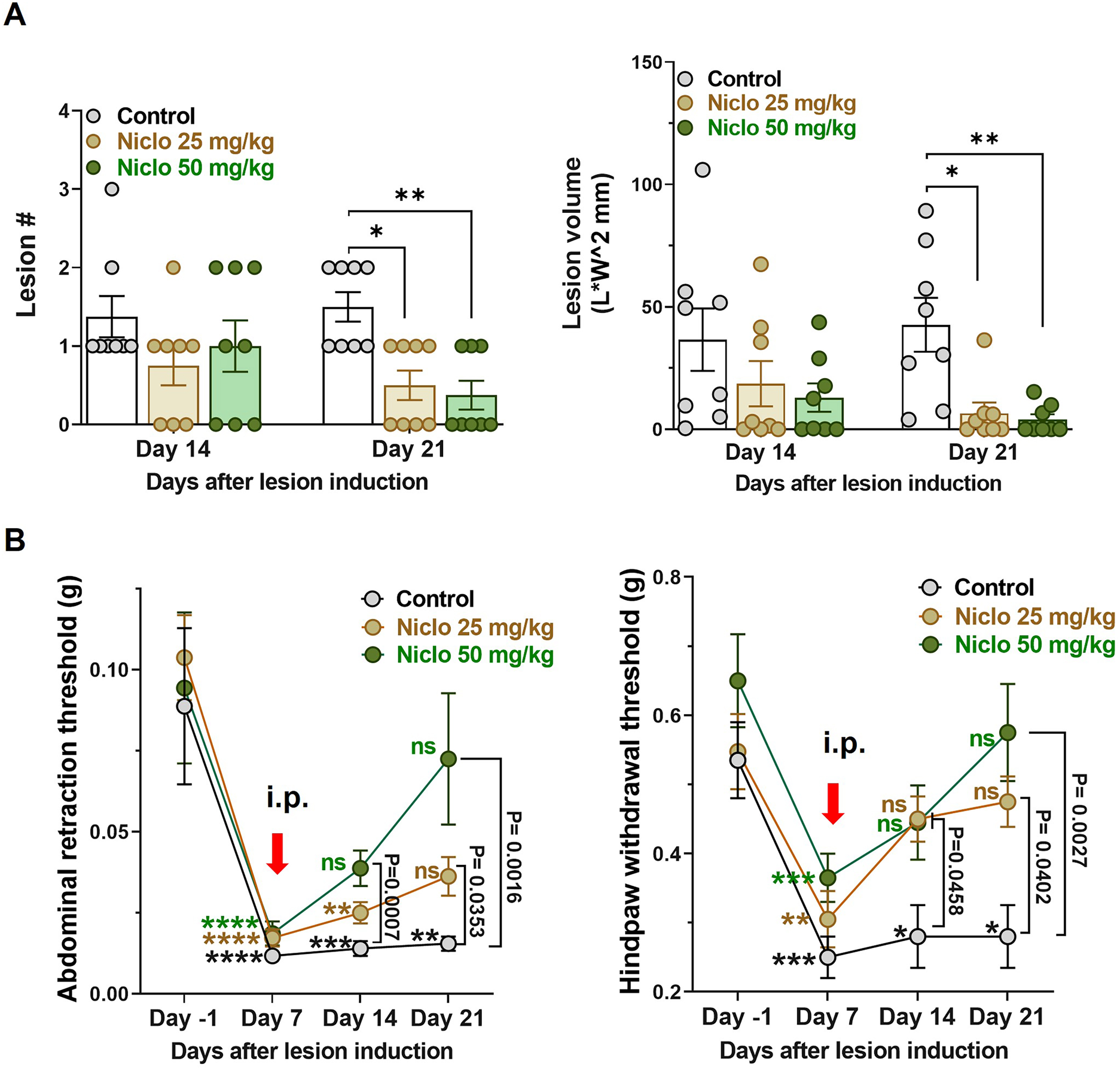
Efficacy of FA-2DG-D-Niclo on treating endometriosis. (**A**) Quantification of lesion numbers and volumes in mice at Days 14 and 21 after lesion induction. Lesion volumes were calculated as length × width^2^. Differences in lesion number and volume among groups at each time point were assessed by the Kruskal-Wallis test. (**B**) Evaluation of endometriosis-associated hyperalgesia. Abdominal (left) and hindpaw (right) withdrawal thresholds were assessed by the von Frey test. Data represent the group mean threshold at the corresponding time point (n=16 at Day -1 and Day 7, n=8 at Day 14 and Day 21). Red arrows show the time of i.p. administration of FA-2DG-D or FA-2DG-D-Niclo. The time course differences within each treatment group were compared using a Kruskal-Wallis test followed by the Dunnett multiple-comparison test. The colored stars or ns (non-significance) represent the differences at the corresponding time point relative to Day -1 within the treatment group. To compare dosing effects of niclosamide at each time point among groups, the Kruskal-Wallis test was used. Data are shown as mean ± SEM (n=8). p < 0.05 (*), p < 0.01 (**), p < 0.001 (***), or p < 0.0001 (****).

## 4. Conclusions

With integrated single-cell transcriptomic analysis, we further identified a significant recruitment of pro-disease FRβ⁺ macrophages within human endometriotic lesions and the pelvic cavity, mirroring our observations in the mouse model. Importantly, overexpression of FRβ on these macrophages provides a ligand-guided entry point, enabling receptor-specific targeting for both imaging and therapeutic interventions. Leveraging this receptor overexpression, we engineered a folic acid-conjugated dendrimer (FA-2DG-D) to exploit FRβ-mediated uptake for targeted imaging and drug delivery in endometriotic lesions. FA-2DG-D offers several critical advantages for translational development. The dendrimer architecture enables precise, reproducible conjugation of targeting ligands and therapeutic payloads via click chemistry, resulting in well-defined nanostructures, excellent aqueous solubility, and robust batch-to-batch consistency. Conjugation of niclosamide to generate FA-2DG-D-Niclo markedly enhanced the solubility of this poorly water-soluble drug, enabled controlled, intracellularly triggered release via a disulfide linker, and minimized premature systemic exposure. Importantly, the formulation demonstrated high stability under physiological and storage conditions. In a mouse model of endometriosis, the Cy5-labelled platform demonstrated lesion and peritoneal cell-specific accumulation and intracellular internalization along with a favorable biodistribution with minimal off-target organ accumulation. FA-2DG-D-Niclo enabled targeted delivery to disease-driving macrophages. Administration of FA-2DG-D-Niclo led to significant reductions in lesion number and volume, and improved hyperalgesia, highlighting both therapeutic efficacy and immunomodulatory activity. This work demonstrates the potential of FA-2DG-D platform to target FRβ⁺ macrophages for the treatment of endometriosis. By addressing key limitations of current hormonal and systemic treatments, this approach establishes a non-hormonal, FRβ⁺ macrophage-focused strategy for the treatment of endometriosis.

## Supporting Information

The chemistry experimental details and supplementary figures related to characterization, stability studies, and bioditribution are provided in supporting information.

## Data Availability Statement

The raw data of scRNA sequencing have been deposited to NCBI/SRA (PRJNA1418167). Additional data or materials related to the findings of this study are available from the corresponding author upon reasonable request.

## Authorship Contribution Statement

A.S. and K.H. designed the research; A.D., A.S., and R.S. designed and synthesized the dendrimers; M.S., M.H., and T.P. performed the animal experiments and/or analyzed the data; A.D., K.J.G., and A.I.D. performed quantitative biodistribution analysis; N.S.M., V.K.S., and P.S.C. collected/or prepared the human samples; M.S. performed single-cell RNA sequencing and analysis; A.D., M.S., R.S., A.S., and K.H. wrote the paper; all authors read, reviewed, edited, and approved the manuscript.

