## Supplementary Information for "2-Deoxyglucose dendrimer-enabled niclosamide delivery to FRβ-expressing macrophages alleviates endometriosis progression and associated hyperalgesia"

*Correspondence:

**Supplementary Information**

**Contents:**

**1. Chemistry Experimental Section**

- **1.1. Materials and Reagents**
- **1.2. Instruments**
- **1.3. Synthetic Procedures**
- **1.4. In Vitro Release Study of FA-2DG-D-Niclo**
- **1.5. Formulation Stability Study of FA-2DG-D-Niclo**

**2. Supplementary Figures**

### *1. Chemistry Experimental Section*

#### *1.1. Materials and Reagents*

All chemicals and reagents were purchased from Sigma-Aldrich or Merck and used as received unless otherwise specified. Analytical-grade solvents were employed for all syntheses. Reactions requiring anhydrous conditions were carried out under an inert atmosphere, with glassware oven-dried prior to use. Analytical thin-layer chromatography (TLC) was performed on silica gel 60 F254 aluminum-backed plates, and visualization was achieved under UV light or by appropriate staining. Purification of compounds was conducted using flash column chromatography on silica gel 60 (230–400 mesh). Dialysis membranes (Spectra/Por) were obtained from Repligen.

#### *1.2. Instruments*

Nuclear magnetic resonance (NMR) spectra were recorded on a Bruker 500 MHz high-resolution spectrometer using deuterated solvents, including chloroform-d ($CDCl_3$) or dimethyl sulfoxide-$d_6$ (DMSO-$d_6$). Chemical shifts (δ) for $^1$H NMR are reported in parts per million (ppm) relative to the residual solvent signal. Coupling constants (J) are given in hertz (Hz). Signal multiplicities are denoted as follows: s = singlet, d = doublet, t = triplet, q = quartet, and m = multiplet. High-resolution mass spectrometry (HRMS) analyses were conducted on a Waters Q-Tof Premier mass spectrometer operated in positive-ion V-mode using electrospray ionization (ESI).

Particle size distribution and zeta potential were determined by dynamic light scattering (DLS) using a Malvern Zetasizer Nano 90 (Malvern Instruments, Westborough, MA) at 25°C. For DLS analysis, samples of FA-2DG-D and FA-2DG-D-Niclo were prepared in deionized water (18.2 MΩ cm) at a concentration of 0.5 mg/mL and filtered through 0.2 μm syringe filters (Pall Corporation, HT Tuffryn membrane) directly into UV-transparent disposable cuvettes (12.5 × 12.5 × 45 mm). Zeta potential measurements were performed at a concentration of 0.2 mg/mL in 10 mM NaCl using Malvern Nanoseries disposable folded capillary cells, following the same sample preparation procedure.

High-performance liquid chromatography (HPLC) was employed to determine the purity of small molecules, dendrimers, and dendrimer conjugates, and to monitor drug release profiles. Analyses were carried out on a Waters Acquity Arc system (Milford, MA, USA) equipped with binary pumps, a 2998 photodiode array (PDA) detector, and a 2475 fluorescence detector, using Waters Empower software. Separations were achieved on a Waters Symmetry C18 column (5 μm, 4.6 × 250 mm) with a gradient elution. To assess the purity of the Cy5-conjugated dendrimer and to evaluate drug release profiles, HPLC analyses were performed using a gradient starting at 90:10 (Solvent A: 0.1% TFA in water; Solvent B: 0.1% TFA in acetonitrile), progressing to 50:50 (A:B) at 20 min, followed by 10:90 (A:B) at 38 min, and re-equilibrated to 90:10 (A:B) at 40 min, at a flow rate of 1 mL/min. For the niclosamide-conjugated dendrimer, the gradient commenced at 80:20 (Solvent A: 0.1% TFA in water; Solvent B: 0.1% TFA in acetonitrile), transitioned to 10:90 (A:B) at 10 min, was maintained at this ratio until 20 min, and re-equilibrated to 80:20 (A:B) at 30 min, with a flow rate of 1 mL/min. Dendrimers and dendrimer conjugates were monitored at 254 nm (aromatic absorption), 280 nm (folic acid absorption), 331 nm and 376 nm (Niclo absorption), while Cy5-labeled conjugate was analyzed at 650 nm (Cy5 absorption). Fluorescence measurements of 2DG-D-Cy5 and FA-2DG-D-Cy5 were performed using a Varioskan Lux Multimode Microplate Reader (Thermo Fisher). Fluorescence was recorded at Cy5 excitation λ_ex = 650 nm and emission λ_em = 670 nm.

#### *1.3. Synthetic Procedures*

***Synthesis of FA-Azide (3):*** Folic acid **(1)** (1.0 g, 2.26 mmol, 1.0 equiv) was dissolved in DMSO (20 mL) with gentle heating and sonication to ensure complete dissolution. DCC (700 mg, 3.40 mmol, 1.5 equiv) and NHS (400 mg, 3.47 mmol, 1.5 equiv) were then added, and the reaction mixture was stirred at room temperature for 16 hours. Subsequently, DIPEA (0.3 mL) was added to adjust the pH between 6 and 8. Azido-PEG5-Amine **(2)** (770 mg, 2.54 mmol, 1.2 equiv) was then added, and the reaction mixture was stirred overnight at room temperature. The reaction mixture was treated with cold diethyl ether (2 × 50 mL), and the precipitate was collected by decantation. The precipitate was washed with ethyl acetate (2 × 20 mL) and dichloromethane (2 × 20 mL). The remaining solvent was removed under reduced pressure to afford FA-Azide **(3)** as a solid in 70% yield.

$^{1}$H NMR (500 MHz, DMSO) δ 8.67 (s, 1H, Aromatic H), 8.09 (d, $J$ = 7.5 Hz, 1H, Amide H), 7.68 (d, $J$ = 8.5 Hz, 2H, Aromatic H), 7.31 (s, 1H, New Amide H), 6.99 (t, $J$ = 6.1 Hz, 1H, Amide H), 6.68 (d, $J$ = 8.5 Hz, 2H, Aromatic H), 4.51 (d, $J$ = 5.9 Hz, 2H, -$CH_2$), 4.37 – 4.29 (m, 1H, -CH), 3.72 – 3.53 (m, 20H, PEG H), 3.42 (t, $J$ = 4.9 Hz, 2H, PEG H), 3.00 (t, $J$ = 5.3 Hz, 2H, PEG H), 2.33 (t, $J$ = 7.5 Hz, 2H, -$CH_2$), 2.10 – 2.02 (m, 1H, -$CH_2$), 1.99 – 1.88 (m, 1H, -$CH_2$). (**Figure S1**)

$^{13}$C NMR (125 MHz, DMSO) δ 174.74, 174.65, 166.62, 161.68, 156.68, 154.68, 151.20, 148.96, 148.86, 129.35, 128.41, 121.91, 111.67, 70.26, 70.22, 70.20, 70.13, 70.11, 70.06, 69.70, 67.14, 52.64, 50.44, 46.38, 38.99, 31.22, 26.98. (**Figure S2**)

HPLC purity: 98.21%, retention time: 13.255 minutes. (**Figure S3**)

m/z *calculated* for $C_{31}H_{42}N_{11}O_{10}$ $[M+Na]^+$ = 750.32; *found* = 750.25 (**Figure S4**)

***Synthesis of 2DG-D-Hexyne (6):*** To a stirred solution of hexynoic acid **(5)** (6.1 mg, 0.054 mmol, 6.0 equiv) in anhydrous DMF (2 mL), EDC·HCl (21 mg, 0.11 mmol, 12.0 equiv) was added, and the mixture was allowed to stir for 15 minutes at room temperature. This activated solution was then added dropwise in a DMF solution (5 mL) containing 2DG-D **(4)** (200 mg, 0.009 mmol, 1.0 equiv) under continuous stirring. Subsequently, DMAP (3.5 mg, 0.027 mmol, 3.0 equiv) was added, and the reaction was maintained at room temperature for 12 hours. Reaction completion was verified by HPLC through observation of a distinct shift in retention time. The crude material was purified by dialysis (1 kDa membrane), initially in DMF for 12 hours and then in deionized water for another 12 hours. The dialyzed solution was freeze-dried to afford 2DG-D-Hexyne **(6)** as a viscous solid in 80% yield.

$^{1}$H NMR (500 MHz, DMSO) δ 8.50 (s, 8H, Amide H), 8.14 – 7.81 (m, 50H, Amide and Triazole H), 7.22 (s, 16H, Aromatic H), 5.18 (d, $J$ = 5.5 Hz, 6H, Sugar H), 4.93 – 4.81 (m, 38H, Sugar H),

4.77 (d, $J$ = 4.9 Hz, 12H, Sugar H), 4.62 – 4.42 (m, 117H, Sugar H), 4.35 – 4.30 (m, 7H, Sugar H), 4.21 – 4.05 (m, 54H, Sugar H), 3.91 – 3.72 (m, 97H, Sugar and PEG H), 3.72 – 3.43 (m, 914H, PEG H), 3.19 – 2.98 (m, 57H, Core and PEG H), 2.78 – 2.86 (m, 10H, Core H), 2.74 – 2.58 (m, 28H, Core and Hexyne H), 2.46 – 2.34 (m, 8H, Hexyne H), 2.24 – 2.13 (m, 28H, Core and Sugar H), 1.97 – 1.79 (m, 35H, Core and Sugar H), 1.75 – 1.68 (m, 9H, Hexyne H), 1.52 – 1.40 (m, 20H, Sugar H). (**Figure S5**)

$^{13}$C NMR (125 MHz, DMSO) δ 166.01, 152.22, 144.24, 140.35, 129.69, 124.71, 106.71, 97.22, 97.12, 84.14, 73.56, 72.32, 72.18, 72.11, 72.04, 70.48, 70.40, 70.30, 70.25, 70.23, 70.21, 70.13, 70.07, 70.01, 69.98, 69.49, 69.41, 69.38, 69.23, 69.16, 68.80, 68.42, 68.22, 66.24, 66.05, 64.15, 63.96, 61.47, 60.66, 49.75, 49.63, 38.79, 38.33, 38.25, 32.79, 25.61, 25.11, 23.98, 17.53. (**Figure S6**)

HPLC purity: 99.87%, retention time: 19.667 minutes. (**Figure S7**)

***Synthesis of FA-2DG-D (7):*** FA-Azide **(3)** (8 mg, 0.011 mmol, 2.5 equiv) was added to a stirred solution of 2DG-D-Hexyne **(6)** (100 mg, 0.0043 mmol, 1.0 eq) in 2 mL DMF. It was followed by addition of $CuSO_4·5H_2O$ (10 mol% per alkyne) dissolved in deionized water (0.1 mL). After 2 min of stirring, sodium ascorbate (15 mol% relative to alkyne) dissolved in deionized water (0.1 mL) was added, and the reaction mixture was stirred at 60 °C for 10 hours. Upon completion, the crude mixture was purified by dialysis (1 kDa membrane) in DMF for 12 hours, followed by dialysis in deionized water for 12 hours. The resulting solution was lyophilized to yield FA-2DG-D **(7)** as a solid in 84% yield.

$^{1}$H NMR (500 MHz, DMSO) δ 8.69 (s, 2H, *FA*-Aromatic H), 8.50 (s, 8H, *Dendrimer*-Amide H), 8.18 – 7.79 (m, 52H, *Dendrimer*-Amide and Triazole H plus *FA*-Amide H), 7.68 (d, $J$ = 8.4 Hz, 4H, *FA*-Aromatic H), 7.22 (s, 16H, *Dendrimer*-Aromatic H), 7.04 (d, $J$ = 5.5 Hz, 2H, *FA*-Amide H), 6.98 (t, $J$ = 6.1 Hz, 2H, *FA*-Amide), 6.68 (d, $J$ = 8.4 Hz, 4H, *FA*-Aromatic H ), 5.18 (d, $J$ = 5.4 Hz, 6H, *Dendrimer*-Sugar H), 4.93 – 4.79 (m, 40H, *Dendrimer*-Sugar H), 4.77 (d, $J$ = 4.9 Hz, 12H, *Dendrimer*-Sugar H), 4.56 – 4.45 (m, 124H, *Dendrimer*-Sugar H and PEG H plus *FA*-$CH_2$), 4.37 – 4.31 (m, 10H, *Dendrimer*-Sugar H plus *FA*-$CH_2$), 4.19 – 4.07 (m, 64H, *Dendrimer*-Sugar H), 3.89 – 3.74 (m, 111H, *Dendrimer*-Sugar H and PEG H), 3.73 – 3.50 (m, 1025H, *Dendrimer*-PEG H plus *FA*-PEG H), 3.19 – 2.98 (m, 50H, *Dendrimer*-Core and PEG H), 2.85 – 2.78 (m, 12H, *Dendrimer*-Core H), 2.73 – 2.59 (m, 34H, *Dendrimer*-Core and Hexyne H), 2.47 – 2.41 (m, 8H,

*Dendrimer*-Hexyne H), 2.27 – 2.10 (m, 42H, *Dendrimer*-Core and Sugar H), 2.04 (t, *J* = 7.6 Hz, 4H, *FA*-$CH_2$), 1.95 – 1.78 (m, 34H, *Dendrimer*-Core and Sugar H plus *FA*-$CH_2$), 1.75 – 1.69 (m, 11H, *Dendrimer*- Hexyne H), 1.55 – 1.42 (m, 20H, *Dendrimer*-Sugar H). (**Figure S8**)

$^{13}$C NMR (125 MHz, DMSO) δ 172.76, 152.22, 144.24, 124.71, 106.71, 97.22, 97.12, 73.56, 72.32, 72.19, 72.11, 72.04, 70.40, 70.30, 70.25, 70.23, 70.21, 70.13, 70.07, 70.01, 69.98, 69.41, 69.38, 69.16, 68.80, 68.42, 68.21, 66.05, 64.15, 63.96, 61.47, 49.75, 38.79, 38.34, 32.79, 25.61, 23.98, 17.53. (**Figure S9**)

HPLC purity: 96.22%, retention time: 16.150 minutes. (**Figure S10**)

***Synthesis of FA-2DG-D-Cy5 (9):*** A solution of Cy5-azide **(8)** (3.0 mg, 0.003 mmol, 2.5 equiv) in DMF (0.1 mL) was added to a solution of FA-2DG-D **(7)** (27 mg, 0.0011 mmol, 1.0 equiv) in DMF (0.9 mL). It was followed by addition of $CuSO_4 \cdot 5H_2O$ (10 mol% per alkyne) dissolved in deionized water (0.1 mL). After 2 min of stirring, sodium ascorbate (15 mol% relative to alkyne) dissolved in deionized water (0.1 mL) was added, and the reaction mixture was stirred at 40 °C for 6 hours. Upon completion, the crude mixture was purified by dialysis (1 kDa membrane) in DMF for 12 hours, followed by dialysis in deionized water for 12 hours. The resulting solution was lyophilized to yield FA-2DG-D-Cy5 **(9)** as a blue solid in 88% yield.

$^{1}$H NMR (500 MHz, DMSO) δ 8.69 (s, 2H, *FA*-Aromatic H), 8.50 (s, 8H, *Dendrimer*-Amide H), 8.41 (t, *J* = 12.9 Hz, 4H, *Cy5*-Aromatic H), 8.17 (d, *J* = 7.7 Hz, 4H, *Cy5*-Aromatic H), 8.11 – 7.80 (m, 55H, *Dendrimer*-Amide and Triazole H plus *FA*-Amide H), 7.72 – 7.64 (m, 6H, *FA*-Aromatic H plus *Cy5*-Aromatic H), 7.38 – 7.34 (m, 2H, *Cy5*-Aromatic H), 7.22 (s, 16H, *Dendrimer*-Aromatic H), 7.04 (d, *J* = 5.6 Hz, 2H, *FA*-Amide H), 6.99 (t, *J* = 6.1 Hz, 2H, *FA*-Amide H), 6.68 (d, *J* = 8.4 Hz, 4H, *FA*-Aromatic H), 6.62 (t, *J* = 12.4 Hz, 2H, *Cy5*-Aromatic H), 6.37 – 6.30 (m, 2H, *Cy5*-Aromatic H), 5.18 (d, *J* = 5.4 Hz, 8H, *Dendrimer*-Sugar H), 4.93 – 4.83 (m, 41H, *Dendrimer*-Sugar H), 4.77 (d, *J* = 4.9 Hz, 11H, *Dendrimer*-Sugar H), 4.55 – 4.44 (m, 142H, *Dendrimer*-Sugar H and PEG H plus *FA*-$CH_2$), 4.37 – 4.30 (m, 10H, *Dendrimer*-Sugar H plus *FA*-$CH_2$), 4.18 – 4.07 (m, 70H, *Dendrimer*-Sugar H), 3.87 – 3.73 (m, 110H, *Dendrimer*-Sugar H and PEG H), 3.70 – 3.51 (m, 1109H, *Dendrimer*-PEG H plus *FA*-PEG H), 3.13 – 3.02 (m, 50H, *Dendrimer*-Core and PEG H), 2.86 – 2.80 (m, 12H, *Dendrimer*-Core H), 2.74 – 2.59 (m, 32H, *Dendrimer*-Core and Hexyne H), 2.44 – 2.40 (m, 12H, *Dendrimer*-Core H), 2.35 (t, *J* = 7.4 Hz, 8H, *Dendrimer*-Hexyne H), 2.26 – 2.19 (m, 28H, *Dendrimer*-Core and Sugar H), 2.16 – 2.10 (m,

8H), 2.09 – 2.05 (m, 4H, *Dendrimer*-Hexyne H), 1.95 – 1.80 (m, 40H, *Dendrimer*-Core and Sugar H plus *FA*-$CH_2$), 1.75 – 1.70 (m, 29H, *Dendrimer*- Hexyne H plus *Cy5*-$CH_2$), 1.51 – 1.42 (m, 21H, *Dendrimer*-Sugar H), 1.30 – 1.24 (m, 14H, *Cy5*-$CH_2$). (**Figure S13**)

HPLC purity: 99.13%, retention time: 17.660 minutes. (**Figure S14**)

***Synthesis of tert-butyl (2-((2-azidoethyl)disulfaneyl)ethyl)(methyl)carbamate (10):*** A solution of sodium methoxide (87.6 mL, 1 M in MeOH) was added dropwise to a stirred mixture of **S-(2-((tert-butoxycarbonyl)(methyl)amino)ethyl) ethanethioate** (10.2 g, 43.8 mmol, 1.0 equiv) and **1,2-bis(2-azidoethyl)disulfane** (8.95 g, 43.8 mmol, 1.0 equiv) in anhydrous methanol (90 mL) at 0°C. The reaction mixture was stirred at room temperature for 3 hours under an argon atmosphere. Upon completion, the solvent was removed under reduced pressure, and the residue was extracted with water (100 mL) and ethyl acetate (3 × 100 mL). The combined organic extracts were dried over anhydrous sodium sulfate, filtered, and concentrated under reduced pressure. Purification by column chromatography afforded compound **10** as a pale-yellow liquid in 35% yield.

$^1H$ NMR (500 MHz, $CDCl_3$) δ 3.58 (t, $J$ = 6.8 Hz, 2H, $N_3$-$CH_2$), 3.53 – 3.45 (m, 2H, *N*-$CH_2$), 2.92 – 2.78 (m, 7H, *S-S*-$CH_2$ and *N*-$CH_3$), 1.45 (s, 9H, *Boc*-$CH_3$). (**Figure S17**)

$^{13}C$ NMR (125 MHz, $CDCl_3$) δ 154.57, 81.42, 79.03, 53.73, 52.21, 29.15, 27.26. (**Figure S18**)

***Synthesis of 2-((2-azidoethyl)disulfaneyl)-N-methylethan-1-aminium chloride (11):*** A solution of compound **10** (15.9 g, 54.3 mmol, 1.0 equiv) in dichloromethane was treated with 4 M HCl in dioxane (68.5 mL) at 0°C. The reaction mixture was stirred at room temperature for 3 hours, after which the solvent was removed under reduced pressure. The resulting residue was suspended in hexane (100 mL) and stirred for 30 min. The supernatant was decanted, and the process was repeated once. The resulting sticky solid was dried under high vacuum to yield compound **11**, which was used in the subsequent step without further purification.

***Synthesis of (2-((2-azidoethyl)disulfaneyl)ethyl)(methyl)carbamic chloride (12):*** To a suspension of compound **11** (4.5 g, 15.4 mmol, 1.0 equiv) in dry DCM (300 mL) at 0°C was added TEA (8.5 mL, 61.5 mmol, 4.0 equiv). The resulting mixture was then introduced dropwise into a precooled solution of triphosgene (4.56 g, 15.4 mmol, 1.0 equiv) in dry DCM maintained at -25°C. After stirring at -25°C for 1 hour, the reaction was allowed to warm to room temperature and stirred for an additional 30 minutes. The solvent was removed under reduced pressure, and the residue was co-distilled with DCM (10 mL × 3) and dried under high vacuum to afford the crude compound **12**, which was used in the subsequent step without further purification.

***Synthesis of Niclo-Azide (14):*** Crude compound **12** (4.5 g, 15.4 mmol, 1.0 equiv) was dissolved in dry DCM (150 mL), and a solution of niclosamide **(13)** (5.7 g, 17.66 mmol, 1.2 equiv) and TEA (5 mL, 35.3 mmol, 3.0 equiv) in dry DCM (110 mL) was added dropwise at room temperature under an inert atmosphere. Catalytic DMAP (400 mg) was then introduced, and the reaction mixture was stirred at room temperature for 16 hours. Upon completion, the solvent was removed under reduced pressure, and the crude product was purified by column chromatography using 30% ethyl acetate in hexane as the eluent. The purified material was suspended in hexane and stirred for 30 minutes, the supernatant was decanted, and the process was repeated twice more. On the third iteration, the solid was collected by filtration, washed with hexane, and dried to afford Niclo-Azide **(14)** as a pale brown solid in 49% yield.

$^1$H NMR (500 MHz, DMSO) δ 10.44 (d, $J$ = 19.7 Hz, 1H, Amide H), 8.44 (d, $J$ = 2.6 Hz, 1H, Aromatic H), 8.34 – 8.29 (m, 1H, Aromatic H), 8.14 (dd, $J$ = 25.7, 9.0 Hz, 1H, Aromatic H), 7.82 (d, $J$ = 2.7 Hz, 1H, Aromatic H), 7.73 – 7.68 (m, 1H, Aromatic H), 7.40 – 7.34 (m, 1H, Aromatic H), 3.69 (t, $J$ = 7.1 Hz, 1H, $N_3$-$CH_2$), 3.60 (t, $J$ = 6.5 Hz, 2H, $N$-$CH_2$), 3.54 (q, $J$ = 5.8 Hz, 1H, $N_3$-$CH_2$), 3.02 (t, $J$ = 7.0 Hz, 1H, $S$-$CH_2$), 2.98 – 2.87 (m, 6H, $S$-$CH_2$ and $N$-$CH_3$). (**Figure S19**)

$^{13}$C NMR (125 MHz, DMSO) δ 163.91, 163.86, 153.66, 153.41, 147.99, 144.86, 144.73, 141.27, 132.19, 132.15, 130.82, 130.78, 129.85, 129.37, 129.33, 127.48, 127.17, 126.28, 125.89, 125.79, 125.77, 125.48, 123.57, 123.53, 50.78, 49.59, 48.49, 48.28, 38.10, 37.18, 37.12, 35.49, 35.46, 35.30, 35.26, 30.90. (**Figure S20**)

HPLC purity: 95.94%, retention time: 14.691 minutes. (**Figure S21**)

m/z *calculated* for $C_{19}H_{18}Cl_2N_6O_5S_2$ $[M+H]^+$ = 543.02; *found* = 543.30. (**Figure S22**)

***Synthesis of 2DG-D-Hexyne' (15):*** To a stirred solution of hexynoic acid **(5)** (12 mg, 0.1 mmol, 12.0 equiv) in anhydrous DMF (4 mL), EDC·HCl (42 mg, 0.22 mmol, 24.0 equiv) was added, and the mixture was allowed to stir for 15 minutes at room temperature. This activated solution was then added dropwise in a DMF solution (5 mL) containing 2DG-D **(4)** (200 mg, 0.009 mmol, 1.0 equiv) under continuous stirring. Subsequently, DMAP (7 mg, 0.054 mmol, 6.0 equiv) was added, and the reaction was maintained at room temperature for 12 hours. Reaction completion was verified by HPLC through observation of a distinct shift in retention time. The crude material was purified by dialysis (1 kDa membrane), initially in DMF for 12 hours and then in deionized water for another 12 hours. The dialyzed solution was freeze-dried to afford 2DG-D-Hexyne' **(21)** as a viscous solid in 82% yield.

$^1$H NMR (500 MHz, DMSO) δ 8.50 (s, 8H, Amide H), 7.97 (m, 48H, Amide and Triazole H), 8.17 – 7.77 (s, 16H, Aromatic H), 5.18 (d, *J* = 5.3 Hz, 6H, Sugar H), 4.95 – 4.82 (m, 38H, Sugar H), 4.77 (d, *J* = 4.9 Hz, 12H, Sugar H), 4.56 – 4.46 (m, 125H, Sugar H), 4.36 – 4.28 (m, 9H, Sugar H), 4.20 – 4.02 (m, 59H, Sugar H), 3.85 – 3.77 (m, 99H, Sugar and PEG H), 3.69 – 3.51 (m, 983H, PEG H), 3.16 – 3.01 (m, 45H, Core and PEG H), 2.87 – 4.79 (m, 11H, Core H), 2.73 – 2.57 (m, 34H, Core H), 2.46 – 2.37 (m, 24H, Hexyne H), 2.27 – 2.11 (m, 42H, Core and Hexyne H), 1.96 – 1.78 (m, 34H, Core and Sugar H), 1.79 – 1.67 (m, 23H, Hexyne H), 1.54 – 1.41 (m, 19H, Sugar H). (**Figure S23**)

$^{13}$C NMR (125 MHz, DMSO) δ 162.78, 152.22, 144.24, 124.71, 106.71, 97.12, 73.56, 72.32, 72.18, 72.11, 72.05, 70.40, 70.29, 70.25, 70.23, 70.21, 70.13, 70.07, 69.98, 69.49, 69.41, 69.38, 69.16, 68.81, 68.42, 66.05, 64.15, 63.96, 61.47, 49.75, 38.33, 37.42, 36.25, 34.56, 31.23, 25.61, 23.97, 17.53, 16.15. (**Figure S24**)

HPLC purity: 97.68%, retention time: 10.363 minutes. (**Figure S25**)

***Synthesis of FA-2DG-D' (16):*** FA-Azide **(3)** (8 mg, 0.011 mmol, 2.5 equiv) was added to a stirred solution of 2DG-D-Hexyne' **(15)** (100 mg, 0.004 mmol, 1.0 eq) in 2 mL DMF. It was followed by addition of $CuSO_4 \cdot 5H_2O$ (10 mol% per alkyne) dissolved in deionized water (0.1 mL). After 2 min of stirring, sodium ascorbate (15 mol% relative to alkyne) dissolved in deionized water (0.1 mL) was added, and the reaction mixture was stirred at 60 °C for 10 hours. Upon completion, the crude mixture was purified by dialysis (1 kDa membrane) in DMF for 12 hours, followed by dialysis in deionized water for 12 hours. The resulting solution was lyophilized to yield FA-2DG-D' **(16)** as a solid in 85% yield.

$^1$H NMR (500 MHz, DMSO) δ 8.69 (s, 2H, *FA*-Aromatic H), 8.13 – 7.83 (m, 8H, *Dendrimer*-Amide H), 8.11 – 7.81 (m, 55H, *Dendrimer*-Amide and Triazole H plus *FA*-Amide H), 7.68 (d, *J* = 8.4 Hz, 4H, *FA*-Aromatic H), 7.22 (s, 16H, *Dendrimer*-Aromatic H), 7.04 (d, *J* = 5.5 Hz, 2H, *FA*-Aromatic H), 6.98 (t, *J* = 6.1 Hz, 2H, *FA*-Amide H), 6.68 (d, *J* = 8.4 Hz, 4H, *FA*-Aromatic H),

5.22 – 5.13 (m, 8H, *Dendrimer*-Sugar H), 4.91 – 4.84 (m, 41H, *Dendrimer*-Sugar H), 4.77 (d, *J* = 5.0 Hz, 12H, *Dendrimer*-Sugar H), 4.55 – 4.46 (m, 138H, *Dendrimer*-Sugar H and PEG H plus *FA*-$CH_2$), 4.37 – 4.29 (m, 10H, *Dendrimer*-Sugar H plus *FA*-$CH_2$), 4.17 – 4.08 (m, 65H, *Dendrimer*-Sugar H), 3.86 – 3.78 (m, 108H, *Dendrimer*-Sugar H and PEG H), 3.69 – 3.51 (m, 1037H, *Dendrimer*-PEG H plus *FA*-PEG H), 3.15 – 3.03 (m, 50H, *Dendrimer*-Core and PEG H), 2.86 - 2.80 (m, 12H, *Dendrimer*-Core H), 2.69 – 2.59 (m, 32H, *Dendrimer*-Core and Hexyne H), 2.46 – 2.38 (m, 24H, *Dendrimer*-Hexyne H), 2.26 – 2.11 (m, 49H, m, 42H, *Dendrimer*-Core and Sugar H), 2.02 – 1.98 (m, 4H, *FA*-$CH_2$), 1.96 – 1.80 (m, 40H, *Dendrimer*-Core and Sugar H plus *FA*-$CH_2$), 1.76 – 1.67 (m, 24H, *Dendrimer*- Hexyne H), 1.52 – 1.41 (m, 21H, *Dendrimer*-Sugar H). (**Figure S26**)

$^{13}$C NMR (125 MHz, DMSO) δ 152.22, 144.24, 124.71, 106.71, 97.22, 97.12, 73.56, 72.32, 72.19, 72.11, 72.04, 70.40, 70.30, 70.25, 70.23, 70.21, 70.13, 70.07, 70.01, 69.98, 69.41, 69.38, 69.16, 68.80, 68.42, 68.21, 66.05, 64.15, 63.96, 61.47, 49.75, 38.79, 38.34, 32.79, 25.61, 23.98, 17.53. (**Figure S27**)

HPLC purity: 99.29%, retention time: 9.456 minutes. (**Figure S28**)

***Synthesis of FA-2DG-D-Niclo (17):*** A solution of FA-2DG-D' **(16)** (27 mg, 0.001 mmol, 1.0 equiv) in deionized water (1 mL) was stirred in a round bottom flask. Niclo-Azide **(14)** (7.56 mg, 0.014 mmol, 10.5 equiv) dissolved in DMF (1 mL) was added to the flask, followed by $CuSO_4 \cdot 5H_2O$ (10 mol% per alkyne) dissolved in deionized water (0.1 mL). After 2 min, sodium ascorbate (15 mol% per alkyne) dissolved in deionized water (0.1 mL) was added, and the reaction mixture was stirred at 40 °C for 10 hours. Upon completion, the crude mixture was purified by dialysis (1 kDa membrane) in DMF for 12 hours, followed by dialysis in deionized water for 12 hours. The resulting solution was lyophilized to afford FA-2DG-D-Niclo **(17)** as a solid in 75% yield.

$^{1}$H NMR (500 MHz, DMSO) δ 10.43 (d, *J* = 14.0 Hz, 10H, *Niclo-Azide*-Amide H), 8.69 (s, 2H, *FA*-Aromatic H), 8.55 – 8.47 (m, 8H, *Dendrimer*-Amide H), 8.43 (s, 10H, *Niclo-Azide*-Aromatic H), 8.30 (d, *J* = 8.9 Hz, 10H, *Niclo-Azide*-Aromatic H), 8.18 – 7.79 (m, 84H, *Dendrimer*-Amide and Triazole H plus *FA*-Amide H plus *Niclo-Azide*-Aromatic H), 7.71 – 7.64 (m, 14H, *FA*-Aromatic H plus *Niclo-Azide*-Aromatic H), 7.41 – 7.33 (m, 10H, *Niclo-Azide*-Aromatic H), 7.21 (s, 16H, *Dendrimer*-Aromatic H), 7.06 – 6.94 (m, 4H, *FA*-Amide H), 6.68 (d, *J* = 8.3 Hz, 4H, *FA*-Aromatic

H), 5.22 – 5.13 (m, 8H, *Dendrimer*-Sugar H), 4.95 – 4.80 (m, 40H, *Dendrimer*-Sugar H), 4.77 (d, $J$ = 4.9 Hz, 12H, *Dendrimer*-Sugar H), 4.62 – 4.46 (m, 160H, *Dendrimer*-Sugar H and PEG H), 4.35 – 4.28 (d, $J$ = 11.5 Hz, 10H, *Dendrimer*-Sugar H plus *FA*-$CH_2$), 4.18 – 4.06 (m, 66H, *Dendrimer*-Sugar H), 3.85 – 3.76 (m, 112H, *Dendrimer*-Sugar H and PEG H), 3.67 – 3.48 (m, 1123H, *Dendrimer*-PEG H plus *FA*-PEG H plus *Niclo-Azide*-$CH_2$), 3.23 – 3.18 (m, 20H, *Dendrimer*-Core and PEG H), 3.13 – 2.99 (m, 70H, *Dendrimer*-Core and PEG H plus *Niclo-Azide*-$CH_2$), 2.97 – 2.81 (m, 43H, *Dendrimer*-Core H plus *Niclo-Azide*-$CH_2$ and -$CH_3$), 2.80 – 2.74 (m, 11H, *Dendrimer*-Core H), 2.67 – 2.61 (m, 39H, *Dendrimer*-Core and Hexyne H), 2.42 – 2.33 (m, 24H, *Dendrimer*-Hexyne H), 2.26 – 2.11 (m, 25H, *Dendrimer*-Core and Sugar H), 2.05 – 2.01 (m, 4H, *FA*-$CH_2$), 2.00 – 1.77 (m, 64H, *Dendrimer*-Core and Sugar H plus *FA*-$CH_2$), 1.53 – 1.41 (m, 18H, *Dendrimer*-Sugar H). (**Figure S29**)

$^{13}$C NMR (125 MHz, DMSO) δ 173.03, 166.02, 152.22, 144.24, 141.26, 140.35, 132.14, 125.48, 124.71, 123.52, 122.66, 106.70, 97.12, 73.55, 72.32, 72.11, 70.40, 70.29, 70.25, 70.21, 70.13, 70.07, 69.98, 69.40, 69.23, 69.16, 68.80, 68.42, 66.05, 63.95, 61.48, 49.75, 48.39, 38.33, 24.78. (**Figure S30**)

HPLC purity: 97.65%, retention time: 11.166 minutes. (**Figure S31**)

#### *1.4. In Vitro Release Study of FA-2DG-D-Niclo*

In vitro drug release studies were performed under three different conditions: (i) PBS (pH 7.4) containing 0.1 mM GSH, (ii) 100% human plasma, and (iii) intracellular-mimicking conditions (citrate buffer, pH 5.5, containing 10 mM GSH). FA-2DG-D-Niclo was dissolved at a concentration of 1 mg/mL in each medium. All samples were incubated at 37°C with continuous shaking to simulate physiological conditions. At predetermined time intervals, aliquots were withdrawn, immediately mixed with an equal volume of methanol, and stored at -20°C until analysis. The released drug was quantified by HPLC, and the cumulative release was calculated based on a standard calibration curve of free Niclo.

#### *1.5. Formulation Stability Study of FA-2DG-D-Niclo*

FA-2DG-D-Niclo formulations (1 mg/mL in PBS) were sterile-filtered using 0.4 µm filters and stored at 4°C, room temperature, and 40°C. Aliquots were collected at 1, 7, 15, and 30 days and analyzed by HPLC to assess purity and stability.

*2. Supplementary Figures*

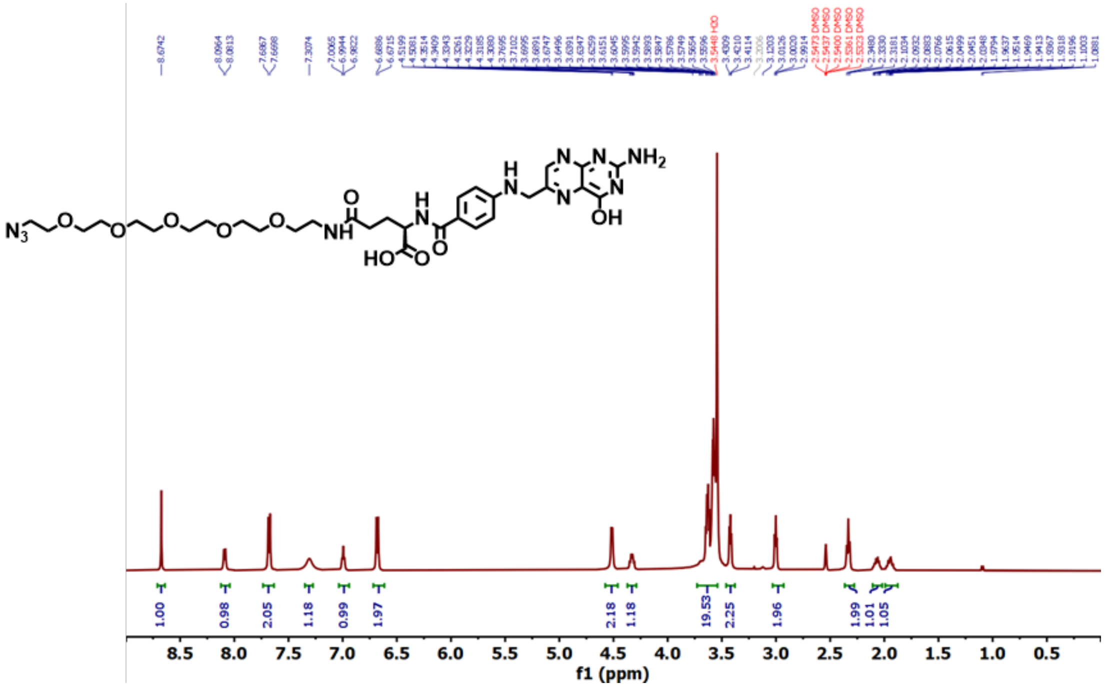


**Figure S1** $^{1}$H NMR of compound **3** in DMSO-$d_6$ (500 MHz).

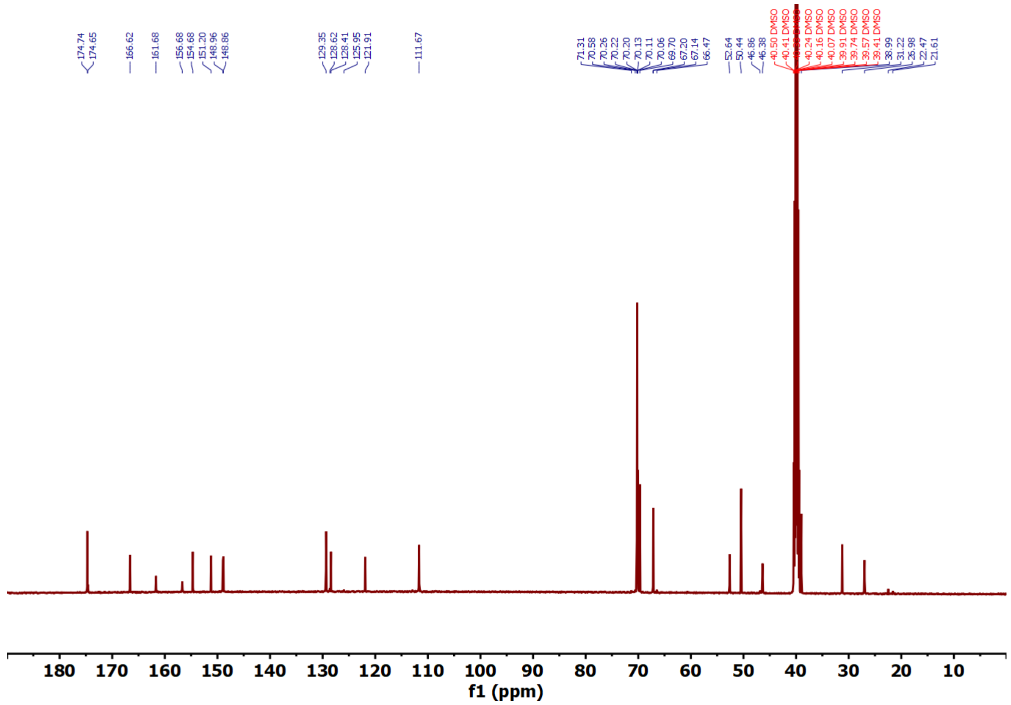


**Figure S2** $^{13}$C NMR of compound **3** in DMSO-$d_6$ (125 MHz).

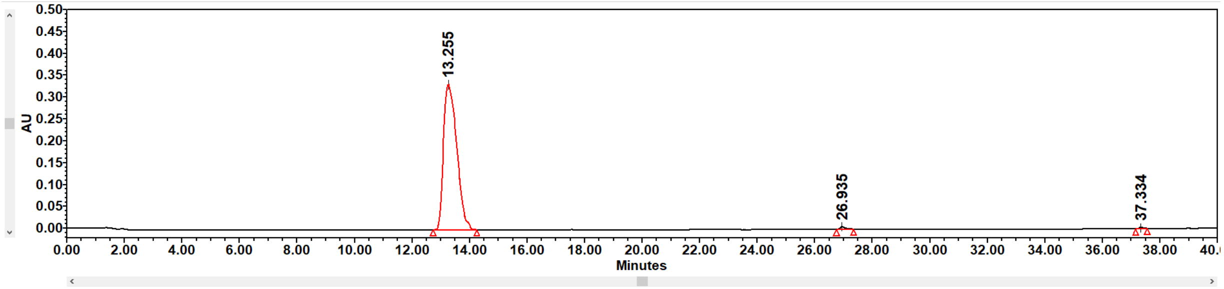


| | Name | Retention Time (min) | Area (µV*sec) | % Area | Height (µV) | Int Type | Amount | Units | Peak Type | Peak Codes |
|---|---|---|---|---|---|---|---|---|---|---|
| 1 | | 13.255 | 10680285 | 98.91 | 332933 | bb | | | Unknown | |
| 2 | | 26.935 | 87014 | 0.81 | 7007 | bb | | | Unknown | |
| 3 | | 37.334 | 30669 | 0.28 | 3199 | bb | | | Unknown | |

**Figure S3** HPLC chromatogram of compound **3** recorded at 280 nm.

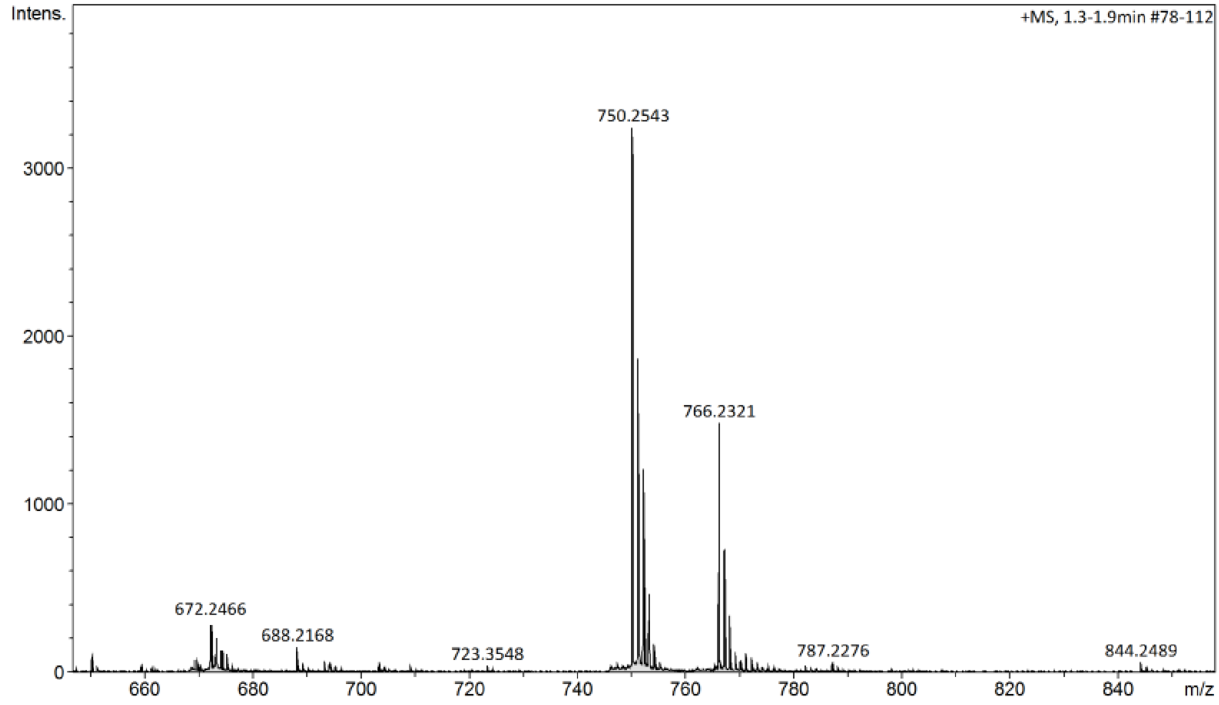


**Figure S4** Mass spectra of compound **3**.

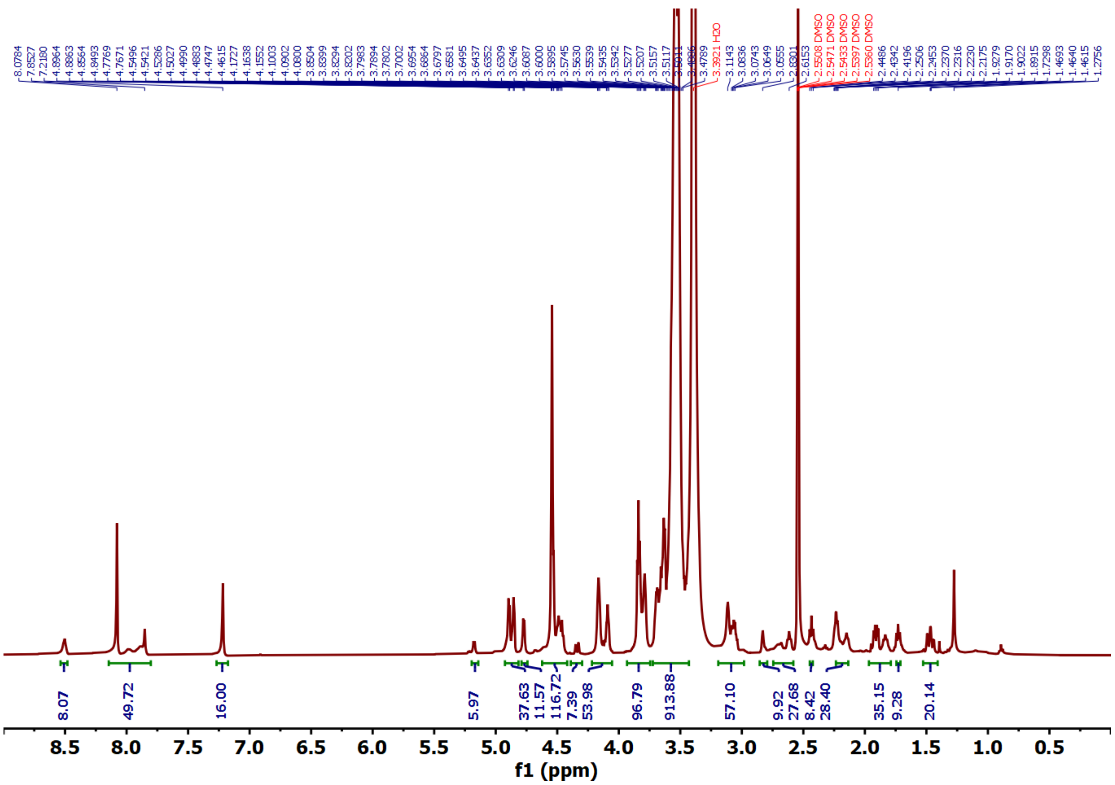

**Figure S5** $^{1}H$ NMR of compound **6** in DMSO-$d_6$ (500 MHz).

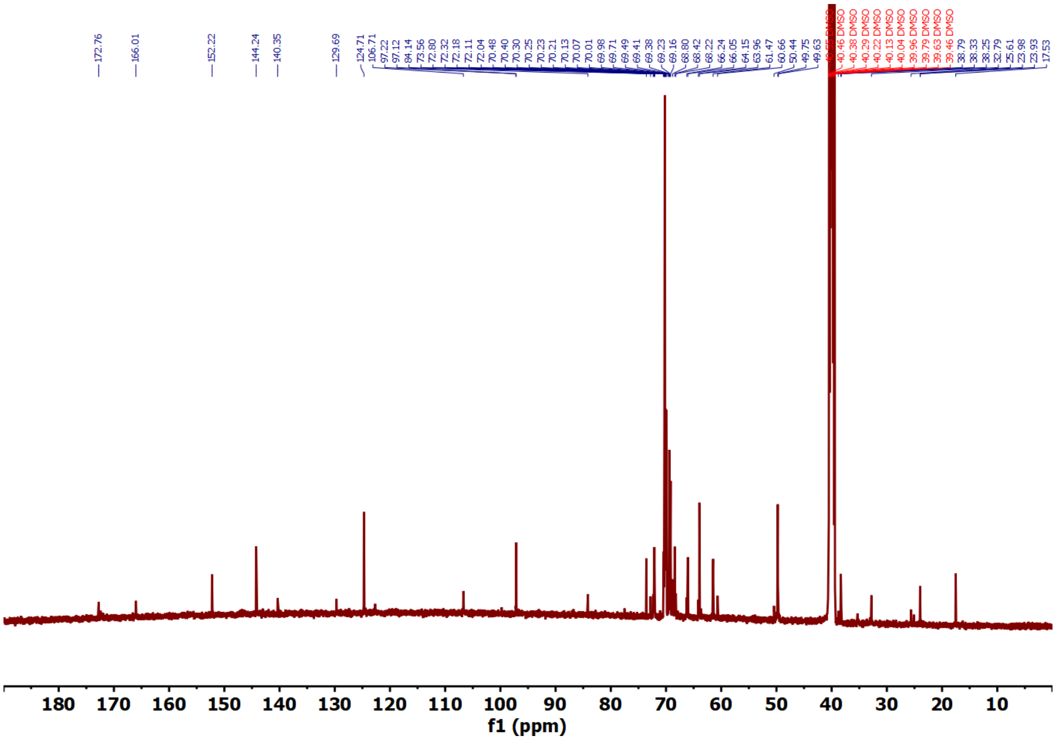

**Figure S6** $^{13}C$ NMR of compound **6** in DMSO-$d_6$ (125 MHz).

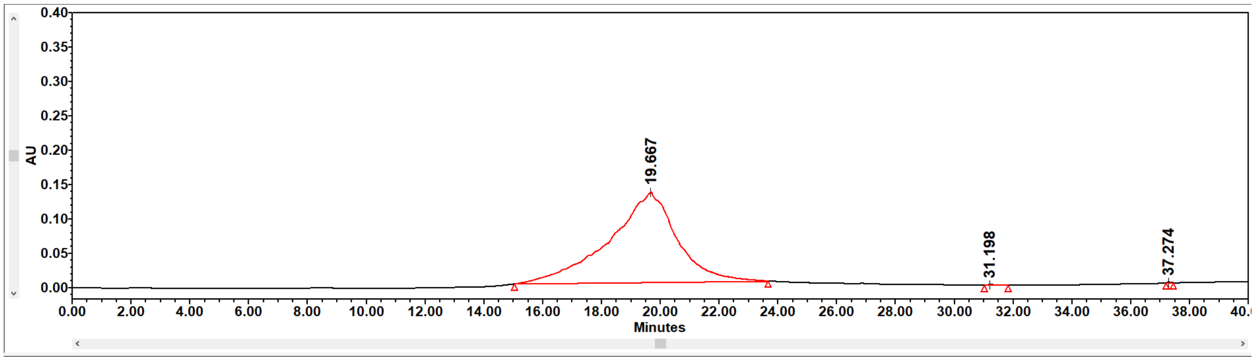


|  | Name | Retention Time (min) | Area (µV*sec) | % Area | Height (µV) | Int Type | Amount | Units | Peak Type | Peak Codes |
|---|---|---|---|---|---|---|---|---|---|---|
| 1 |  | 19.667 | 20784257 | 99.87 | 131592 | bb |  |  | Unknown |  |
| 2 |  | 31.198 | 17829 | 0.09 | 1097 | bb |  |  | Unknown |  |
| 3 |  | 37.274 | 9629 | 0.05 | 1296 | bb |  |  | Unknown |  |

**Figure S7** HPLC chromatogram of compound **6** recorded at 254 nm.

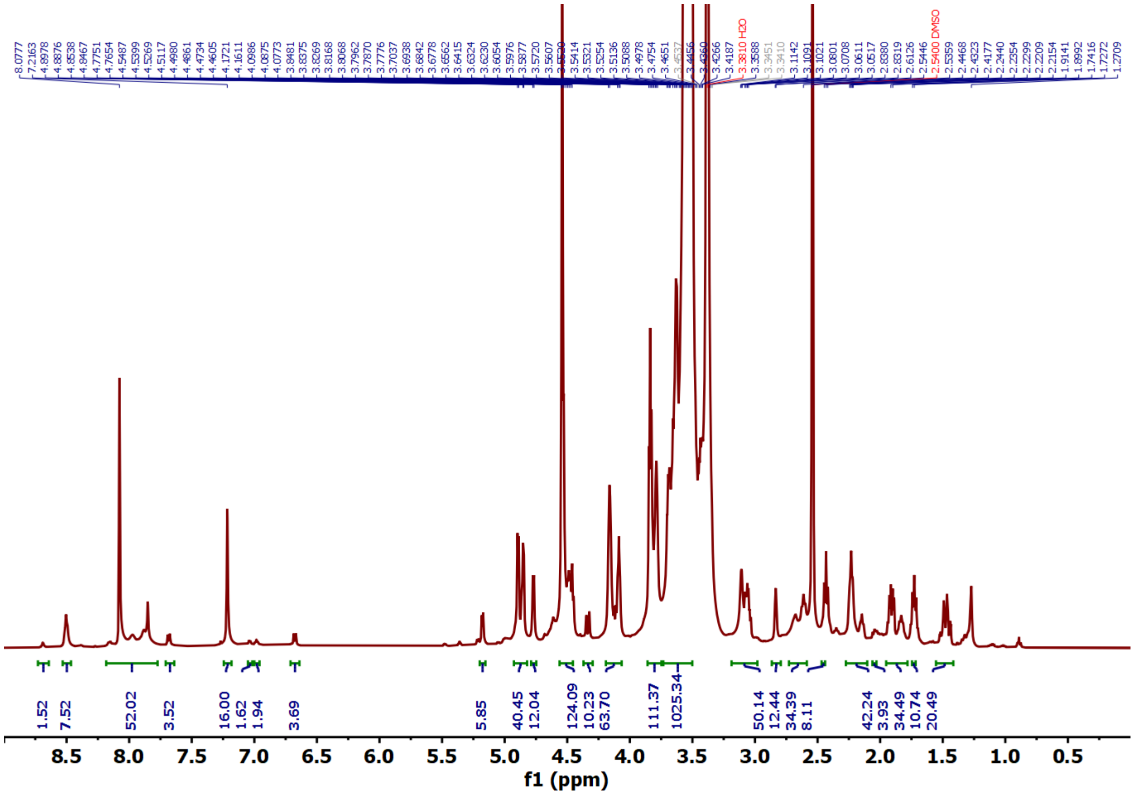


**Figure S8** $^1$H NMR of compound **7** in DMSO-$d_6$ (500 MHz).

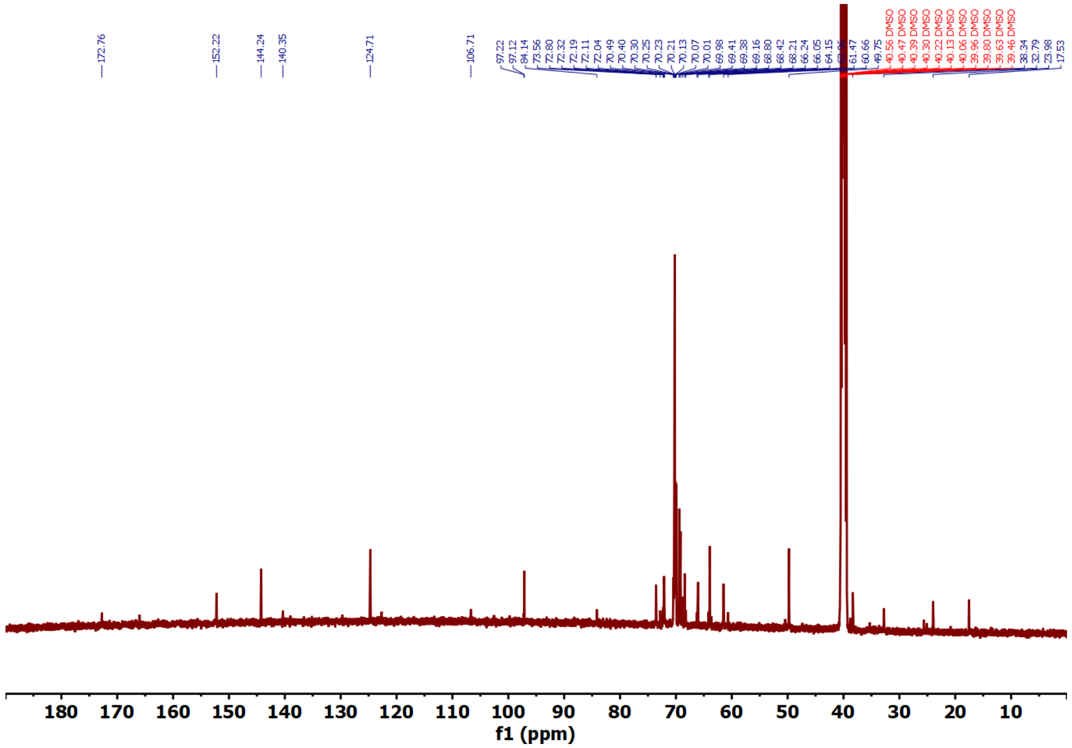


**Figure S9** $^{13}C$ NMR of compound **7** in DMSO-$d_6$ (125 MHz).

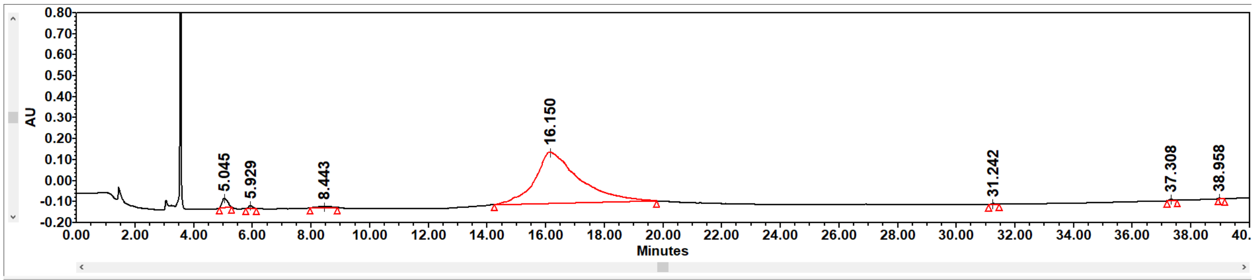


| | Name | Retention Time (min) | Area (µV*sec) | % Area | Height (µV) | Int Type | Amount | Units | Peak Type | Peak Codes |
|---|---|---|---|---|---|---|---|---|---|---|
| 1 | | 5.045 | 132747 | 2.14 | 10157 | bb | | | Unknown | |
| 2 | | 5.929 | 30371 | 0.49 | 3465 | bb | | | Unknown | |
| 3 | | 8.443 | 42533 | 0.68 | 1263 | bb | | | Unknown | |
| 4 | | 16.150 | 5980731 | 96.22 | 56598 | bb | | | Unknown | |
| 5 | | 31.242 | 10821 | 0.17 | 1020 | bb | | | Unknown | I08 |
| 6 | | 37.308 | 16230 | 0.26 | 1758 | bb | | | Unknown | I08 |
| 7 | | 38.958 | 2065 | 0.03 | 390 | bb | | | Unknown | |

**Figure S10** HPLC chromatogram of compound **7** recorded at 280 nm.

Statistics Table

| Name | Mean | Standard Deviation | RSD | Minimum | Maximum |
|---|---|---|---|---|---|
| Z-Average (nm) | 199.5 | 52.75 | 26.44 | 168.7 | 260.4 |
| Peak One Area by Number (%) | 99.97 | 0.05255 | 0.05256 | 99.91 | 100 |
| Peak One Mean by Number (nm) | 6.563 | 0.4248 | 6.473 | 6.157 | 7.005 |
| Peak One Width by Number (nm) | 1.545 | 0.3625 | 23.46 | 1.303 | 1.962 |
| Peak Two Area by Number (%) | 0.09127 | - | - | 0.09127 | 0.09127 |
| Peak Two Mean by Number (nm) | 22.37 | - | - | 22.37 | 22.37 |
| Peak Two Width by Number (nm) | 5.86 | - | - | 5.86 | 5.86 |

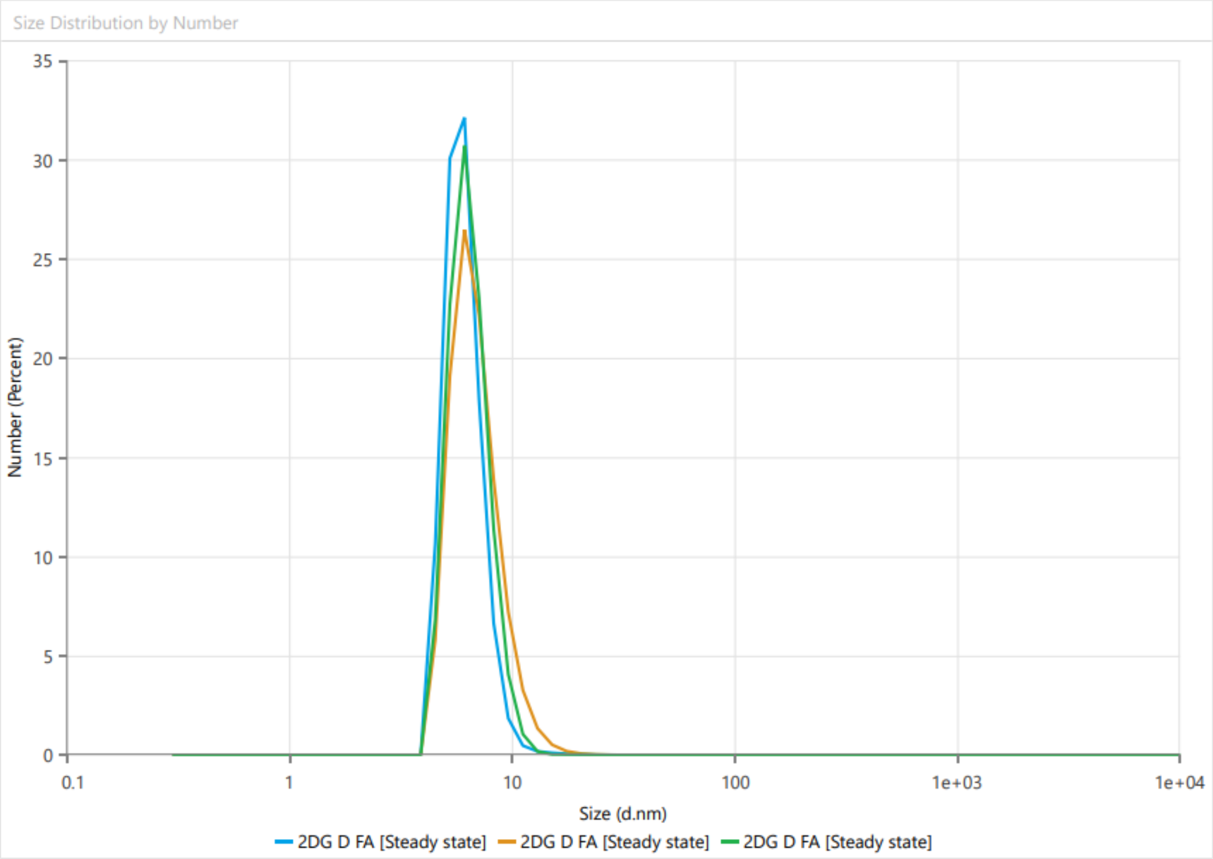


**Figure S11** Size report of compound **7** as measured by dynamic light scattering (DLS).

Statistics Table

| Name | Mean | Standard Deviation | RSD | Minimum | Maximum |
|---|---|---|---|---|---|
| Zeta Potential (mV) | -5.79 | 0.2625 | 4.534 | -6.064 | -5.54 |
| Conductivity (mS/cm) | 0.007917 | 0 | 0 | 0.007917 | 0.007917 |
| Zeta Deviation (mV) | 3.617 | 0.08599 | 2.377 | 3.555 | 3.715 |
| Wall Zeta Potential (mV) | -5.583 | 1.484 | 26.57 | -7.29 | -4.605 |
| Total Count Rate | 734.3 | 429.8 | 58.54 | 475.4 | 1230 |
| Mean Count Rate (kcps) | 271.6 | 0 | 0 | 271.6 | 271.6 |
| Zeta Peak One Area | 100 | 0 | 0 | 100 | 100 |
| Zeta Peak One Mean | -5.672 | 0.2469 | 4.354 | -5.899 | -5.409 |
| Zeta Peak One Mode | -5.607 | 0.6976 | 12.44 | -6.343 | -4.955 |
| Zeta Peak One Width | 3.559 | 0.1077 | 3.026 | 3.483 | 3.683 |

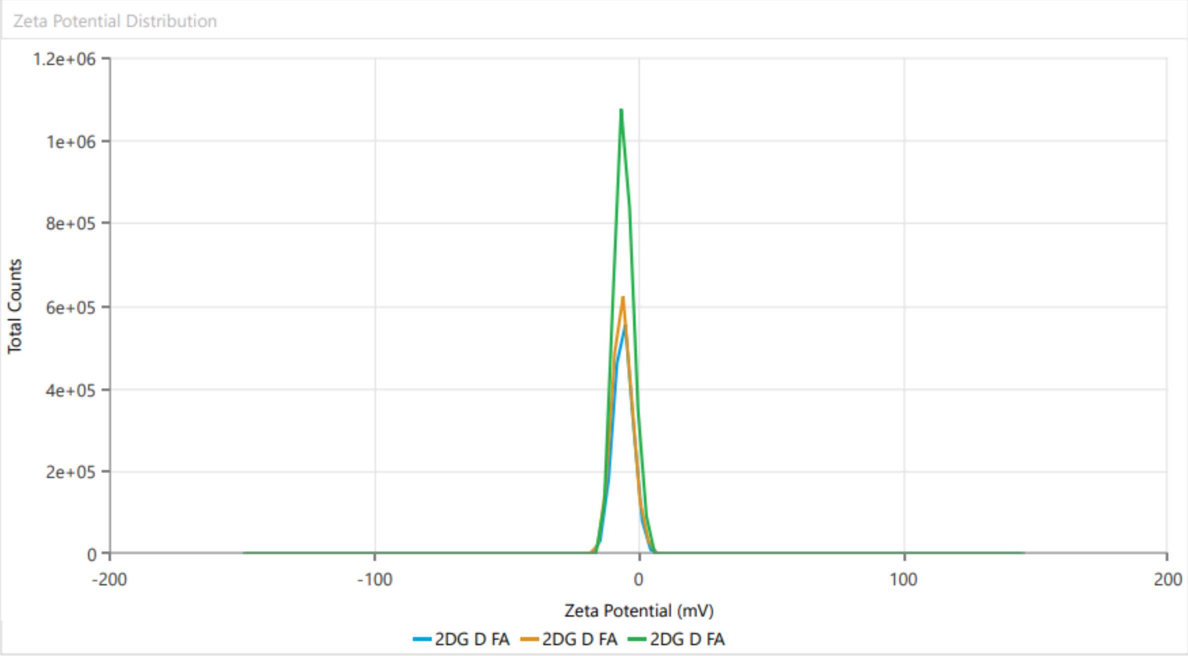


**Figure S12** Zeta potential report of compound **7**.

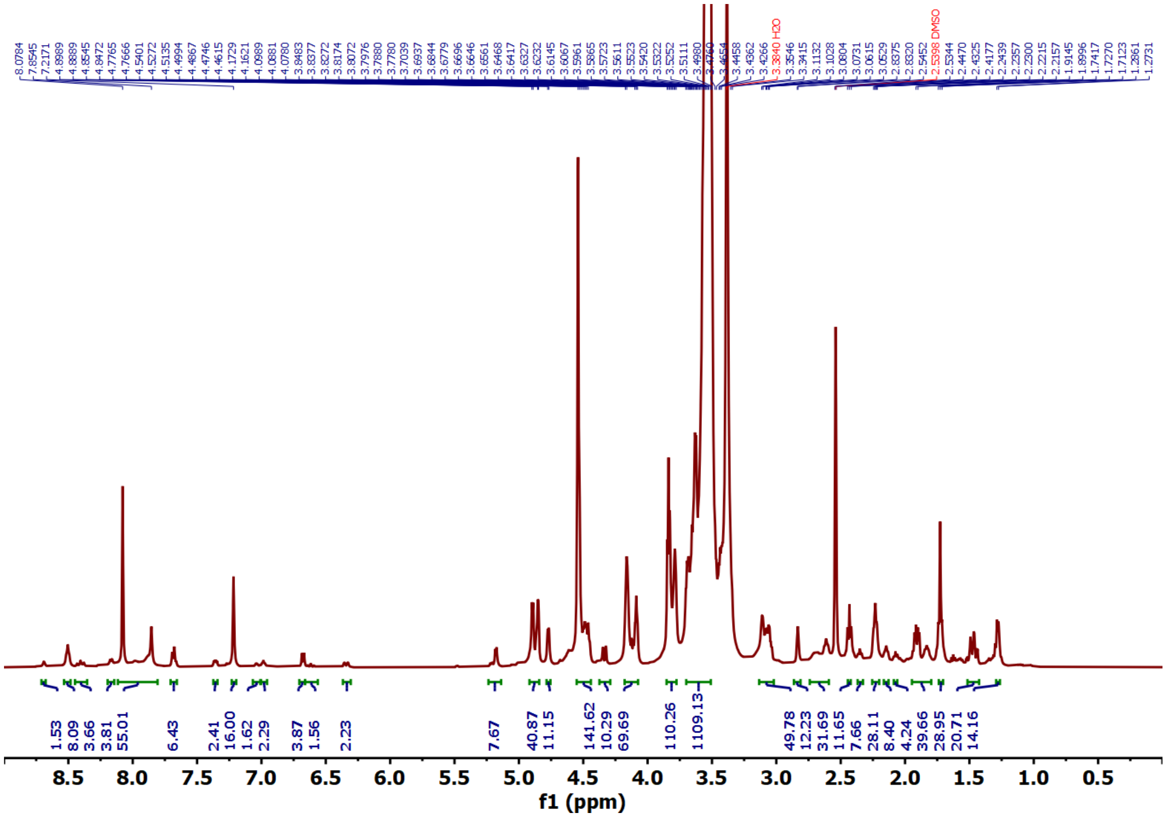


**Figure S13** $^{1}H$ NMR of compound **9** in DMSO-$d_6$ (500 MHz).

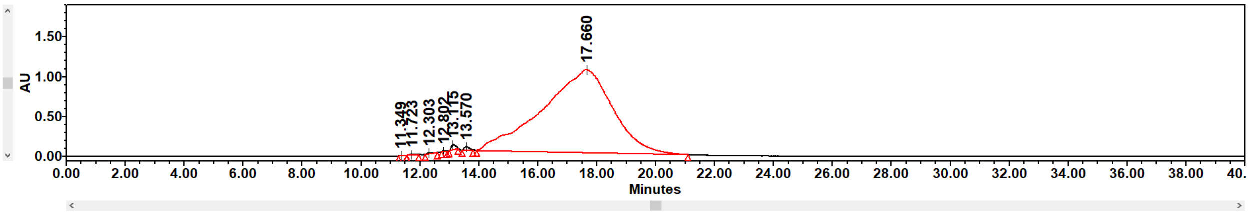


| | Name | Retention Time (min) | Area (µV*sec) | % Area | Height (µV) | Int Type | Amount | Units | Peak Type | Peak Codes |
|---|---|---|---|---|---|---|---|---|---|---|
| 1 | | 11.349 | 25656 | 0.01 | 3466 | bb | | | Unknown | |
| 2 | | 11.723 | 146275 | 0.08 | 11258 | bb | | | Unknown | |
| 3 | | 12.303 | 141611 | 0.08 | 12856 | bb | | | Unknown | |
| 4 | | 12.802 | 42696 | 0.02 | 6015 | bb | | | Unknown | |
| 5 | | 13.115 | 665206 | 0.39 | 63300 | bb | | | Unknown | I08 |
| 6 | | 13.570 | 478327 | 0.28 | 40028 | bb | | | Unknown | |
| 7 | | 17.660 | 170752669 | 99.13 | 1040924 | bb | | | Unknown | |

**Figure S14** HPLC chromatogram of compound **9** recorded at 650 nm.

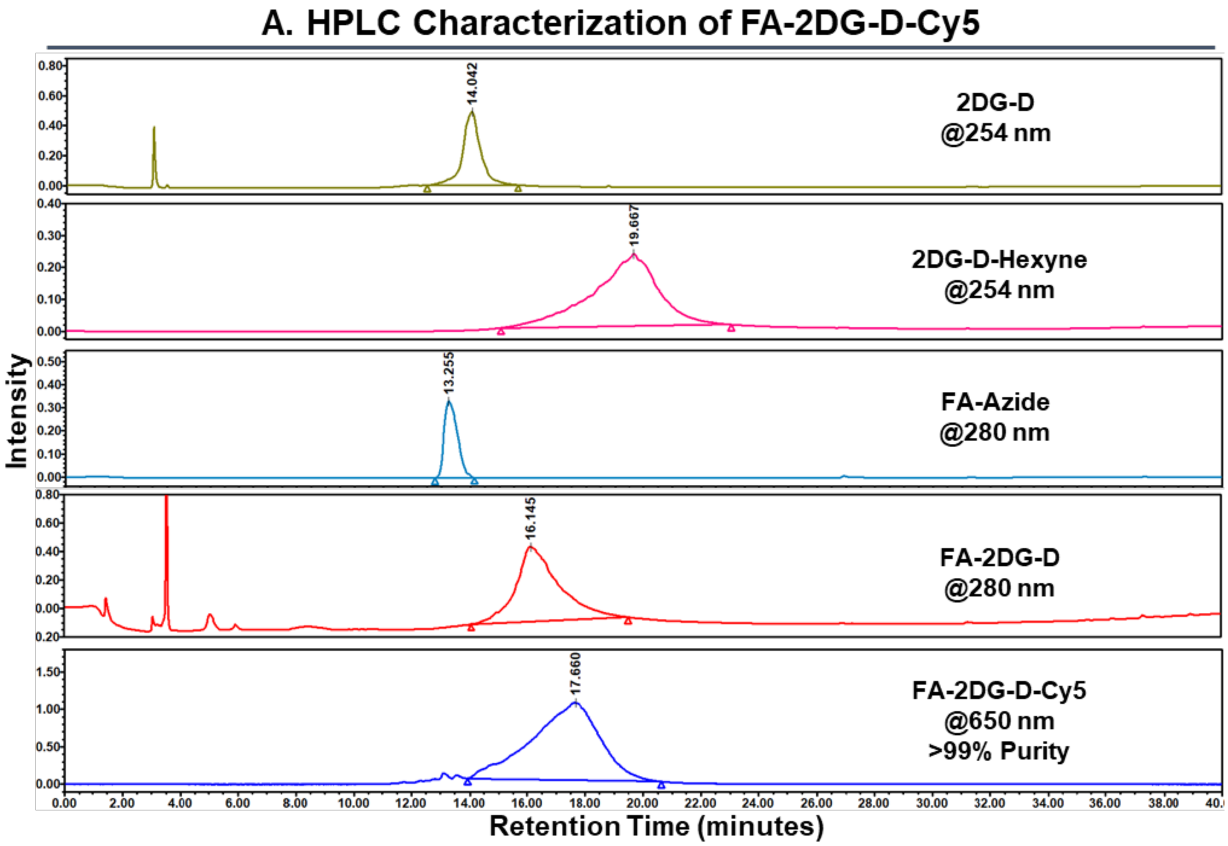


**Figure S15:** HPLC chromatograms of intermediates at various synthesis steps and percentage purity of FA-2DG-D-Cy5.

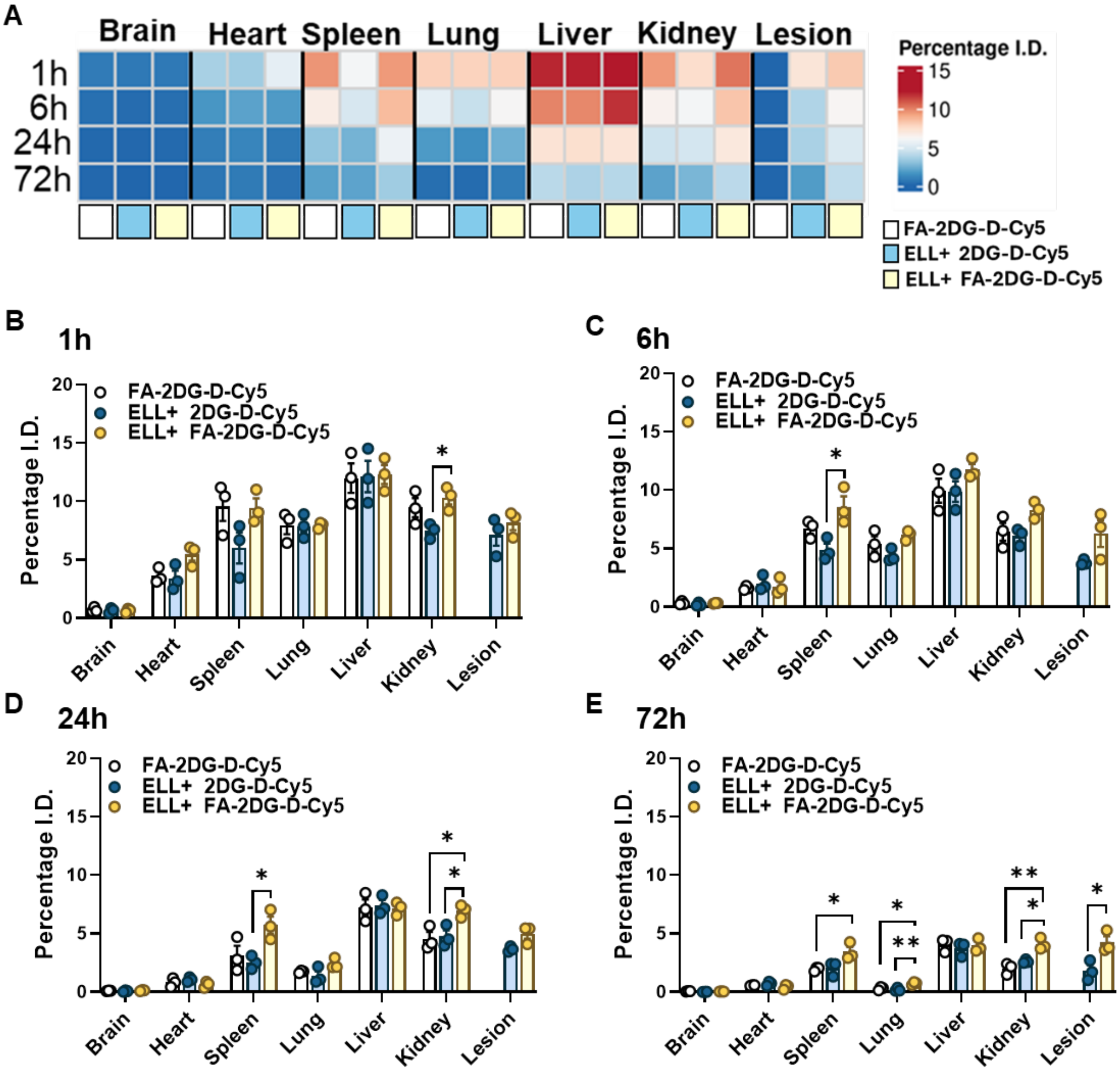


**Figure S16:** Quantitative biodistribution of FA-2DG-D-Cy5.(A) Heat map showing the mean percentage of injected dose in lesions and major organs at 1, 6, 24, and 72 hours after FA-2DG-D-Cy5 administration in sham controls, ELL-induced mice, and 2DG-D-Cy5 administered ELL-induced mice. Quantitative biodistribution of FA-2DG-D-Cy5 and 2DG-D-Cy5 in lesions and major organs at 1 (B), 6 (C), 24 (D), and 72 hour (E) time points in sham and ELL-induced mice (n = 3). Data were obtained via fluorescence spectroscopy of homogenized tissues and shown as a percentage of the injected dose per organ. Differences in each tissue between groups for data in (B), (C), (D), and (E) were analyzed by one-way ANOVA followed by Tukey multiple comparison tests. $p < 0.05$ (*), $p < 0.01$ (**). ELL, endometriosis-like lesion.

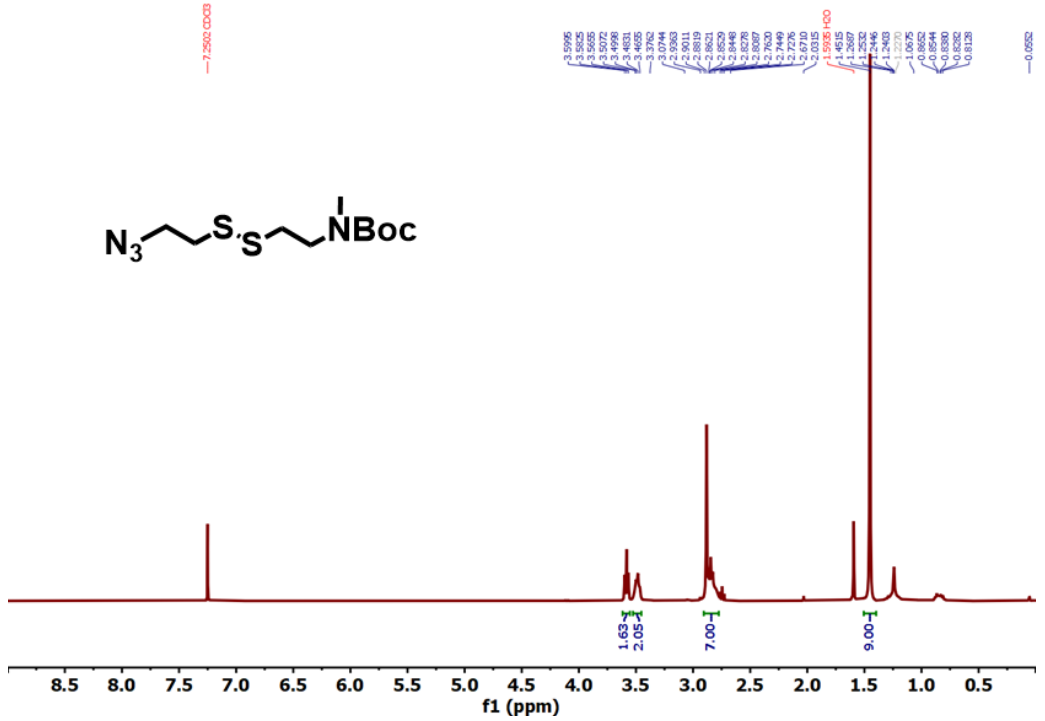


**Figure S17** $^{1}H$ NMR of compound **10** in $CDCl_3$ (500 MHz).

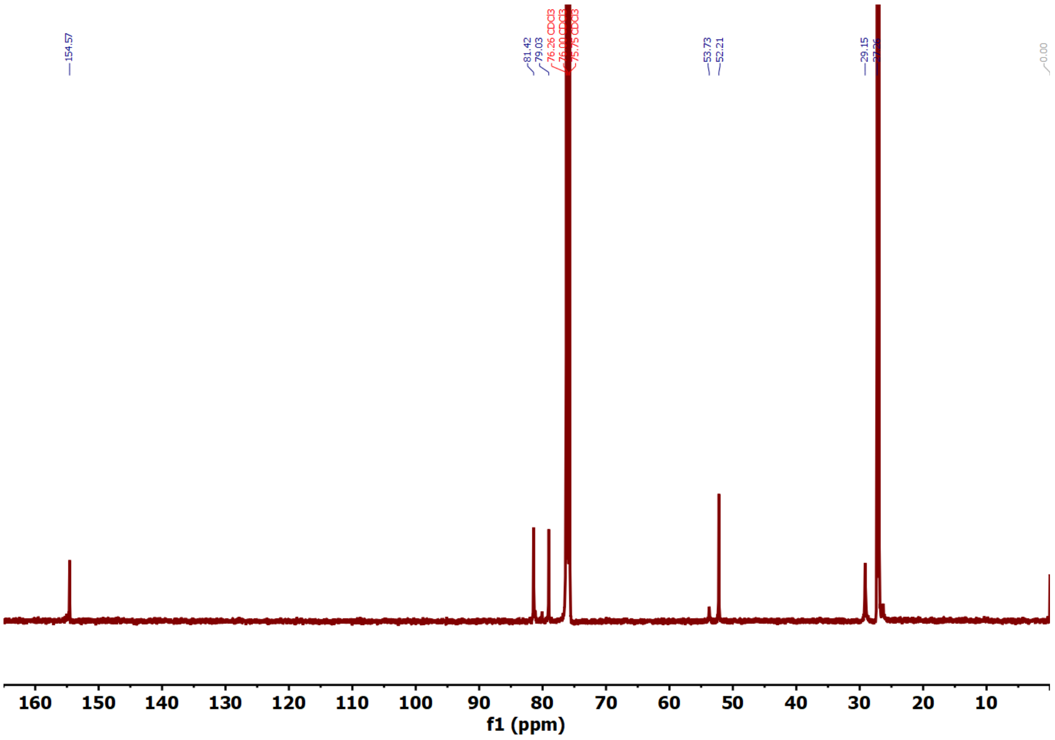


**Figure S18** $^{13}C$ NMR of compound **10** in $CDCl_3$ (125 MHz).

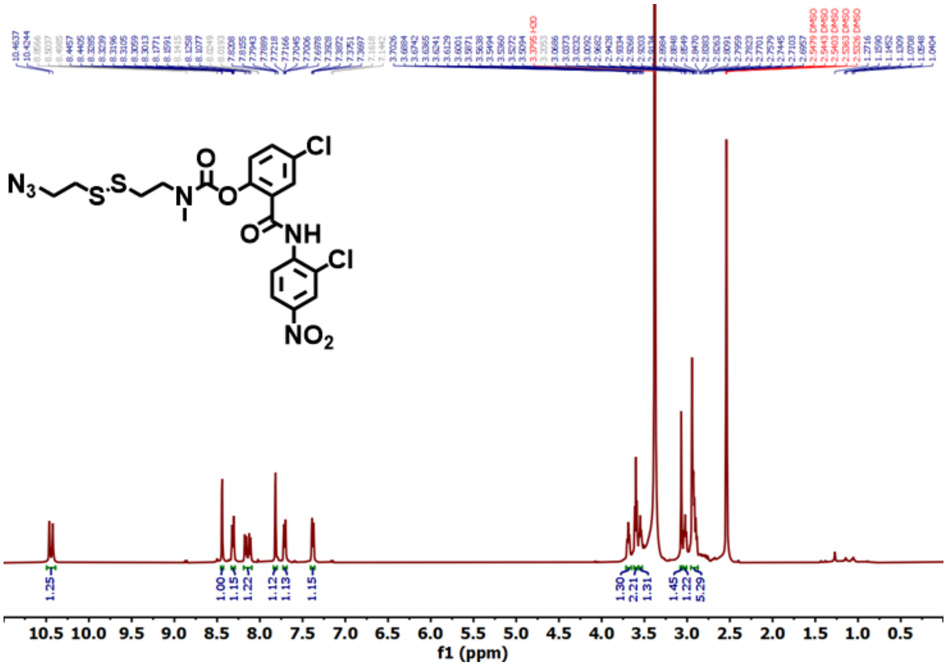


**Figure S19** $^{1}$H NMR of compound **14** in DMSO-$d_6$ (500 MHz).

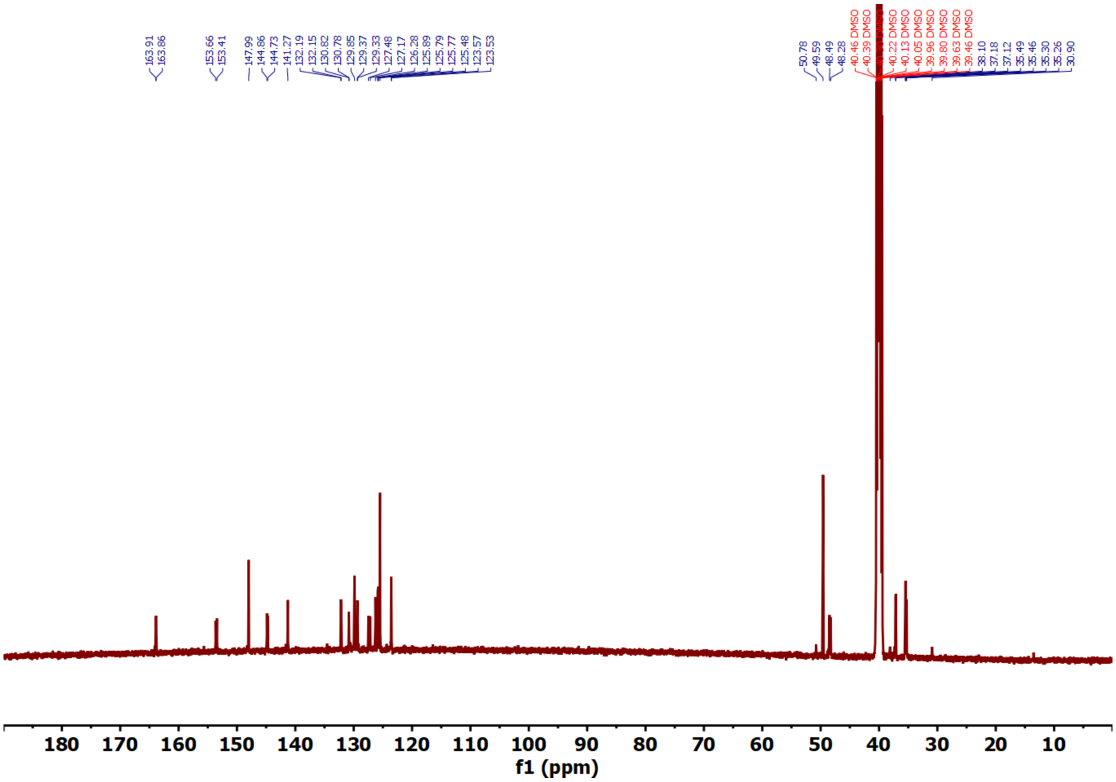


**Figure S20** $^{13}$C NMR of compound **14** in DMSO-$d_6$ (125 MHz).

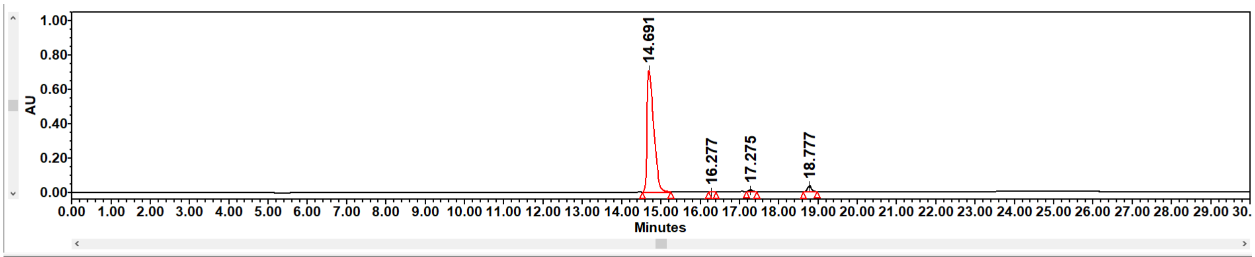


| | Name | Retention Time (min) | Area (µV*sec) | % Area | Height (µV) | Int Type | Amount | Units | Peak Type | Peak Codes |
|---|---|---|---|---|---|---|---|---|---|---|
| 1 | | 14.691 | 8448590 | 95.94 | 712289 | bb | | | Unknown | |
| 2 | | 16.277 | 6770 | 0.08 | 1190 | bb | | | Unknown | |
| 3 | | 17.275 | 84528 | 0.96 | 12119 | bb | | | Unknown | |
| 4 | | 18.777 | 265961 | 3.02 | 35699 | bb | | | Unknown | |

**Figure S21** HPLC chromatogram of compound **14** recorded at 331 nm.

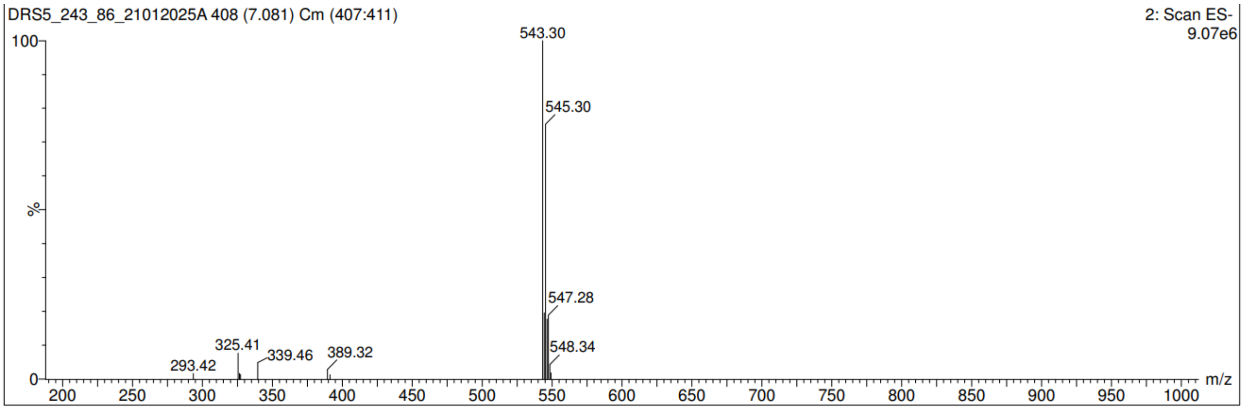


**Figure S22** Mass spectra of compound **14**.

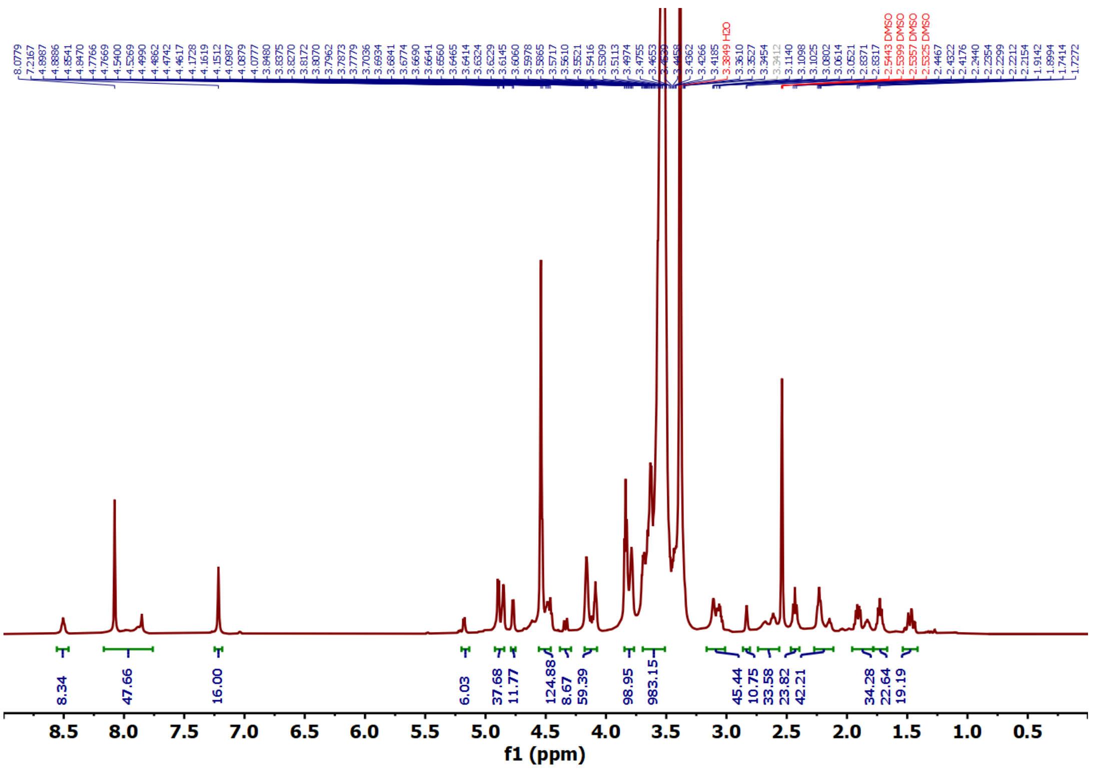

**Figure S23** $^{1}$H NMR of compound **15** in DMSO-d$_6$ (500 MHz).

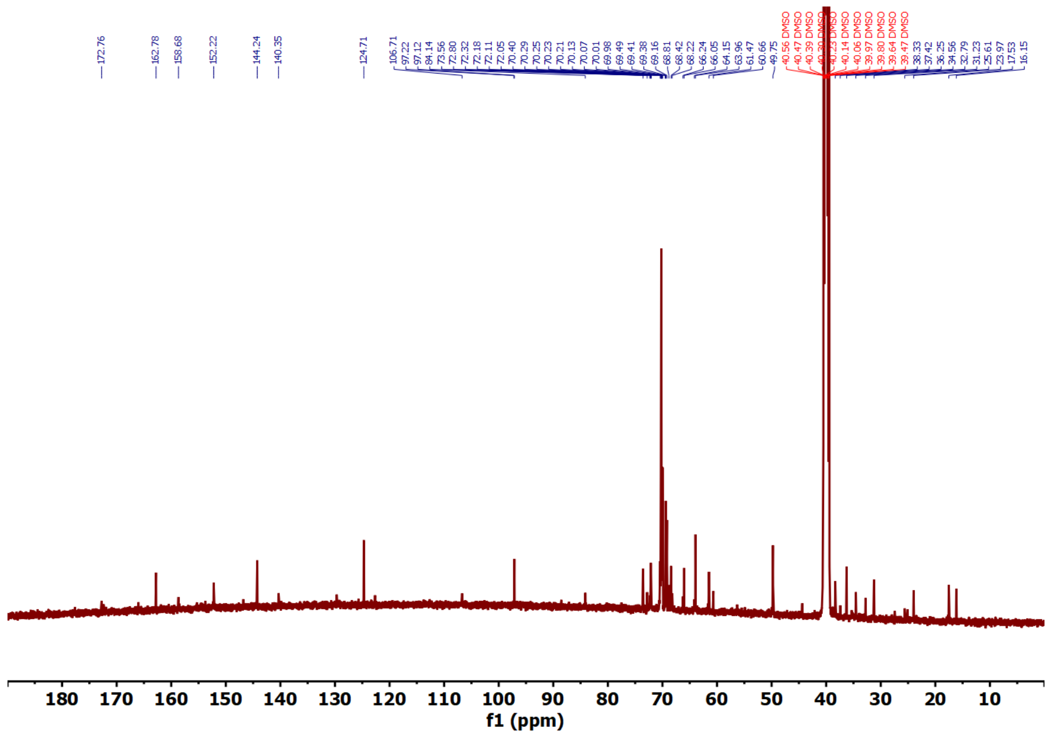

**Figure S24** $^{13}$C NMR of compound **15** in DMSO-d$_6$ (125 MHz).

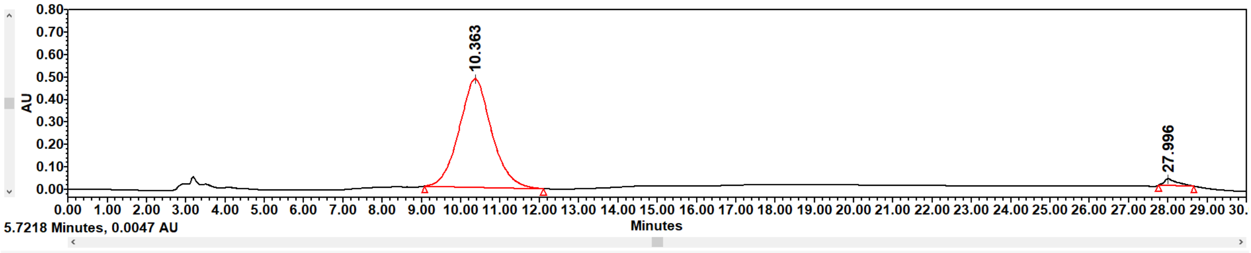


| | Name | Retention Time (min) | Area (μV*sec) | % Area | Height (μV) | Int Type | Amount | Units | Peak Type | Peak Codes |
|---|---|---|---|---|---|---|---|---|---|---|
| 1 | | 10.363 | 12203183 | 97.68 | 211542 | bb | | | Unknown | |
| 2 | | 27.996 | 290131 | 2.32 | 12600 | bb | | | Unknown | |

**Figure S25** HPLC chromatogram of compound **15** recorded at 254 nm.

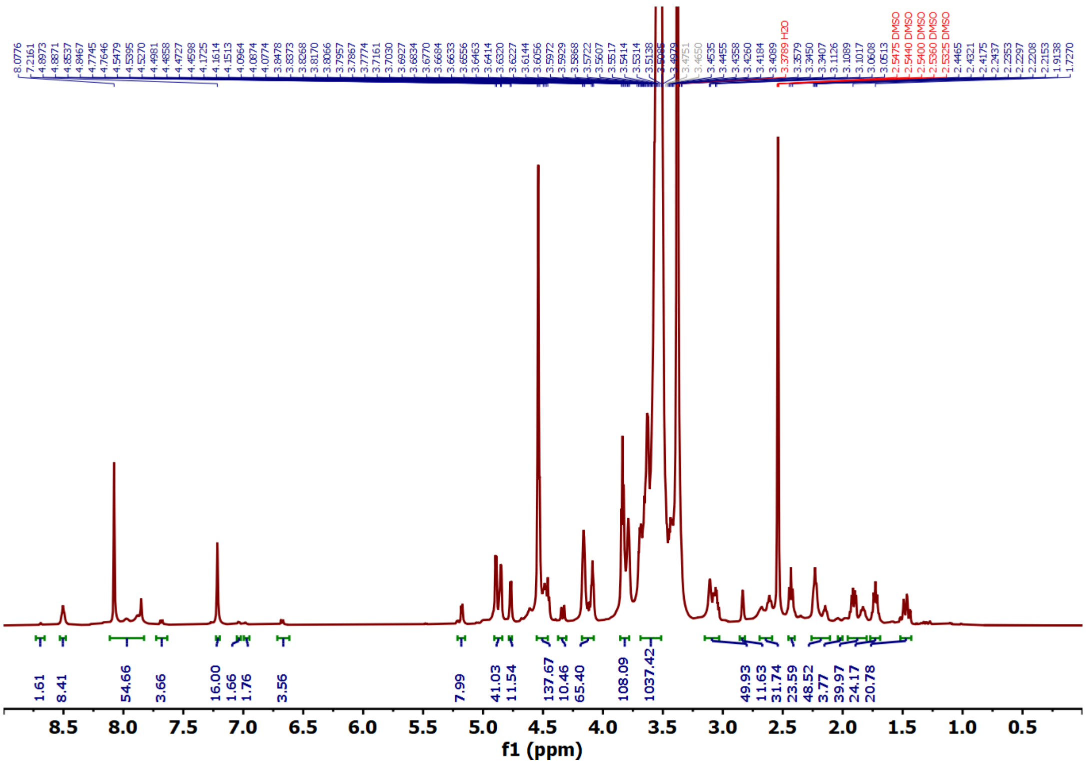


**Figure S26** $^{1}$H NMR of compound **16** in DMSO-$d_6$ (500 MHz).

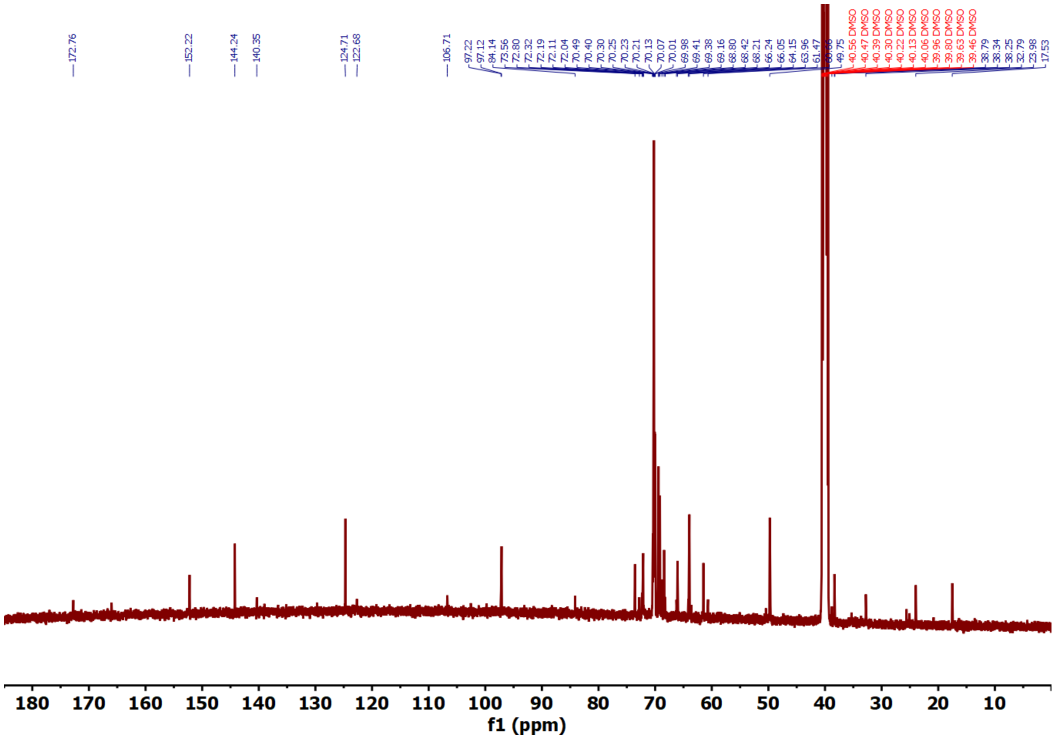


**Figure S27** $^{13}$C NMR of compound **16** in DMSO-d$_6$ (125 MHz).

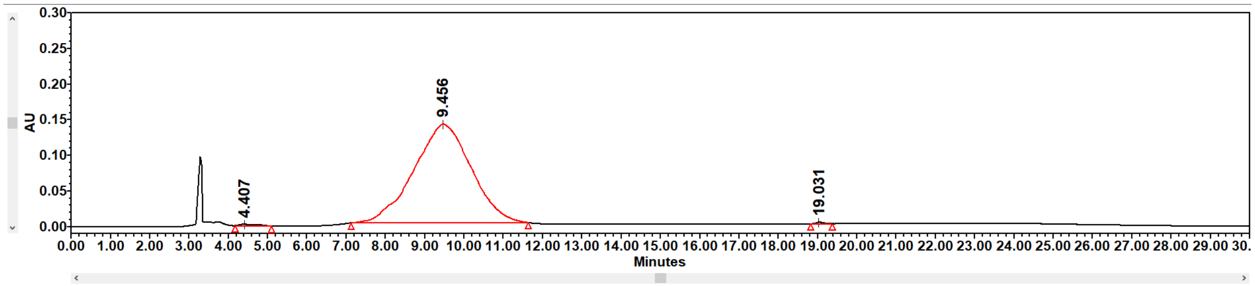


| | Name | Retention Time (min) | Area (µV*sec) | % Area | Height (µV) | Int Type | Amount | Units | Peak Type | Peak Codes |
|---|---|---|---|---|---|---|---|---|---|---|
| 1 | | 4.407 | 69403 | 0.49 | 2009 | bb | | | Unknown | |
| 2 | | 9.456 | 14065999 | 99.29 | 138460 | bb | | | Unknown | |
| 3 | | 19.031 | 31315 | 0.22 | 2227 | bb | | | Unknown | |

**Figure S28** HPLC chromatogram of compound **16** recorded at 254 nm.

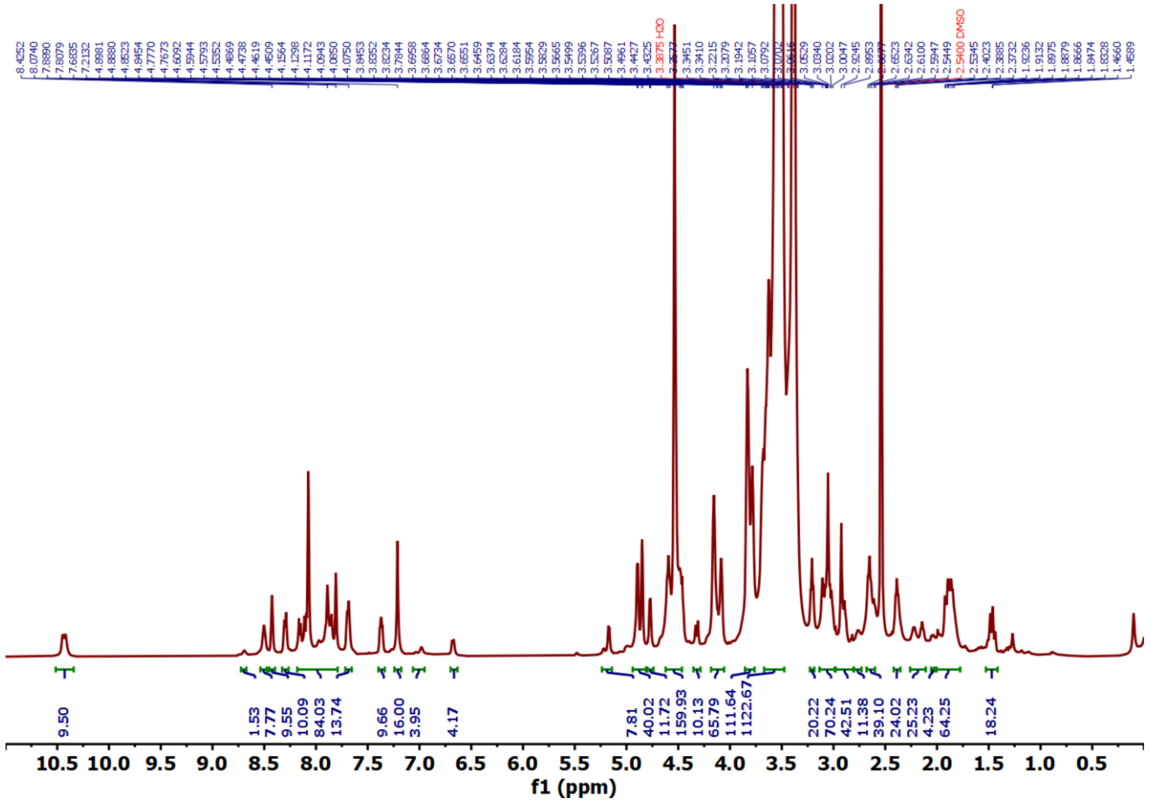

**Figure S29** $^{1}H$ NMR of compound **17** in DMSO-$d_6$ (500 MHz).

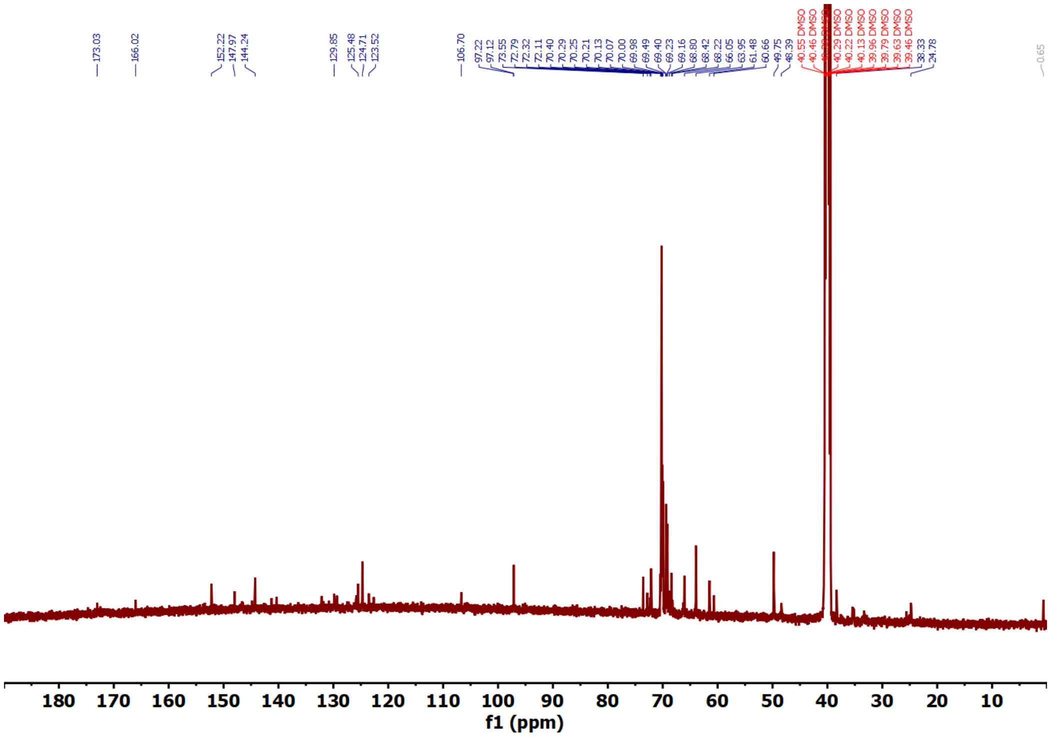

**Figure S30** $^{13}C$ NMR of compound **17** in DMSO-$d_6$ (125 MHz).

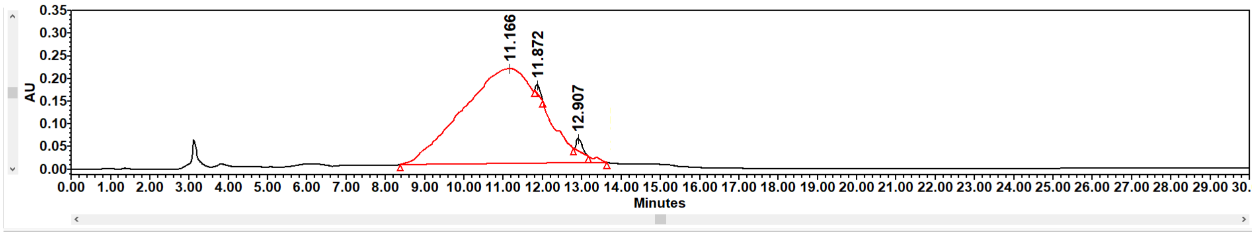


| | Name | Retention Time (min) | Area (µV*sec) | % Area | Height (µV) | Int Type | Amount | Units | Peak Type | Peak Codes |
|---|---|---|---|---|---|---|---|---|---|---|
| 1 | | 11.166 | 32981071 | 98.69 | 209027 | bb | | | Unknown | |
| 2 | | 11.872 | 158404 | 0.47 | 19643 | tt | | | Unknown | I08 |
| 3 | | 12.907 | 283412 | 0.84 | 26946 | tt | | | Unknown | I08 |

**Figure S31** HPLC chromatogram of compound **17** recorded at 331 nm.

Statistics Table

| Name | Mean | Standard Deviation | RSD | Minimum | Maximum |
|---|---|---|---|---|---|
| Z-Average (nm) | 327.9 | 321.1 | 97.93 | 66.08 | 686.2 |
| Peak One Area by Number (%) | 99.98 | 0.02977 | 0.02977 | 99.94 | 100 |
| Peak One Mean by Number (nm) | 7.601 | 0.5683 | 7.477 | 7.256 | 8.257 |
| Peak One Width by Number (nm) | 1.415 | 0.07306 | 5.163 | 1.346 | 1.491 |
| Peak Two Area by Number (%) | 0.03494 | 0.03427 | 98.08 | 0.01071 | 0.05917 |
| Peak Two Mean by Number (nm) | 32.7 | 12.23 | 37.41 | 24.05 | 41.35 |
| Peak Two Width by Number (nm) | 11.17 | 4.513 | 40.4 | 7.98 | 14.36 |

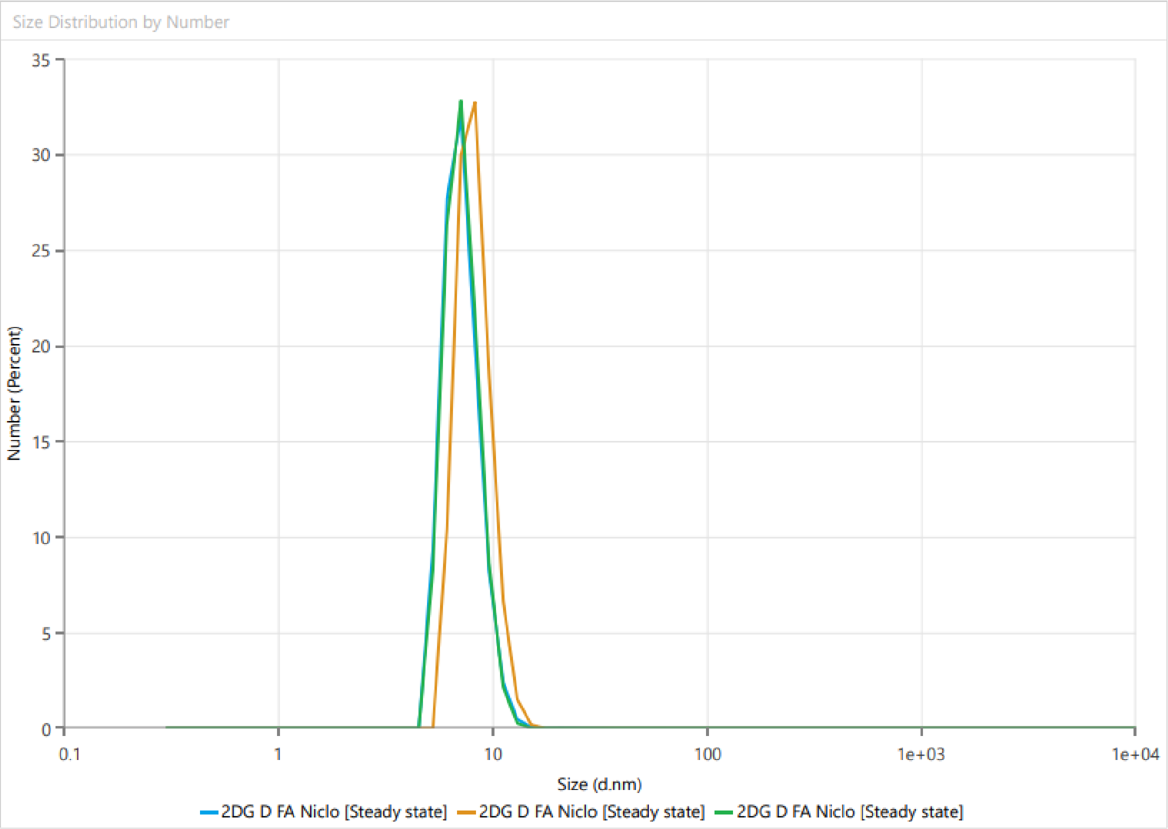


**Figure S32** Size report of compound **17** as measured by dynamic light scattering (DLS).

Statistics Table

| Name | Mean | Standard Deviation | RSD | Minimum | Maximum |
| --- | --- | --- | --- | --- | --- |
| Zeta Potential (mV) | -7.699 | 0.6212 | 8.068 | -8.371 | -7.146 |
| Conductivity (mS/cm) | 0.007801 | 0 | 0 | 0.007801 | 0.007801 |
| Zeta Deviation (mV) | 4.043 | 0.2027 | 5.013 | 3.855 | 4.258 |
| Wall Zeta Potential (mV) | -8.085 | 1.093 | 13.52 | -9.284 | -7.144 |
| Total Count Rate | 645.3 | 267.6 | 41.48 | 458.5 | 951.9 |
| Mean Count Rate (kcps) | 349.5 | 0 | 0 | 349.5 | 349.5 |
| Zeta Peak One Area | 100 | 0 | 0 | 100 | 100 |
| Zeta Peak One Mean | -7.689 | 0.7121 | 9.26 | -8.496 | -7.147 |
| Zeta Peak One Mode | -8.002 | 0.1767 | 2.208 | -8.13 | -7.8 |
| Zeta Peak One Width | 4.047 | 0.2041 | 5.043 | 3.861 | 4.265 |

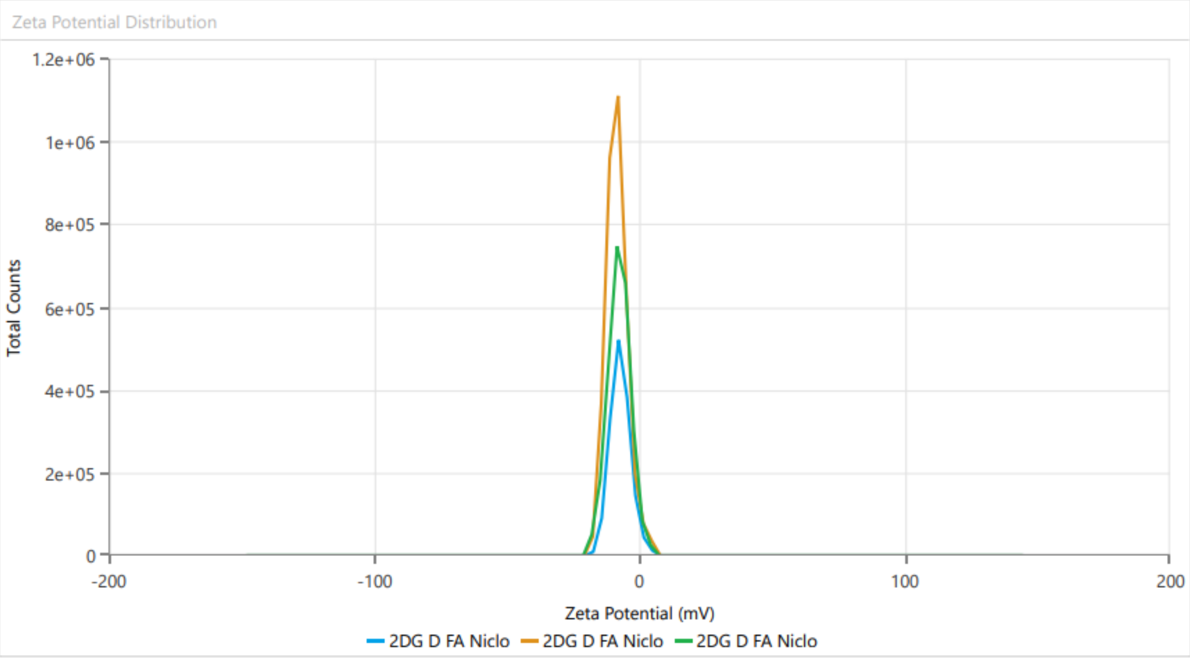


**Figure S33** Zeta potential report of compound **17**.

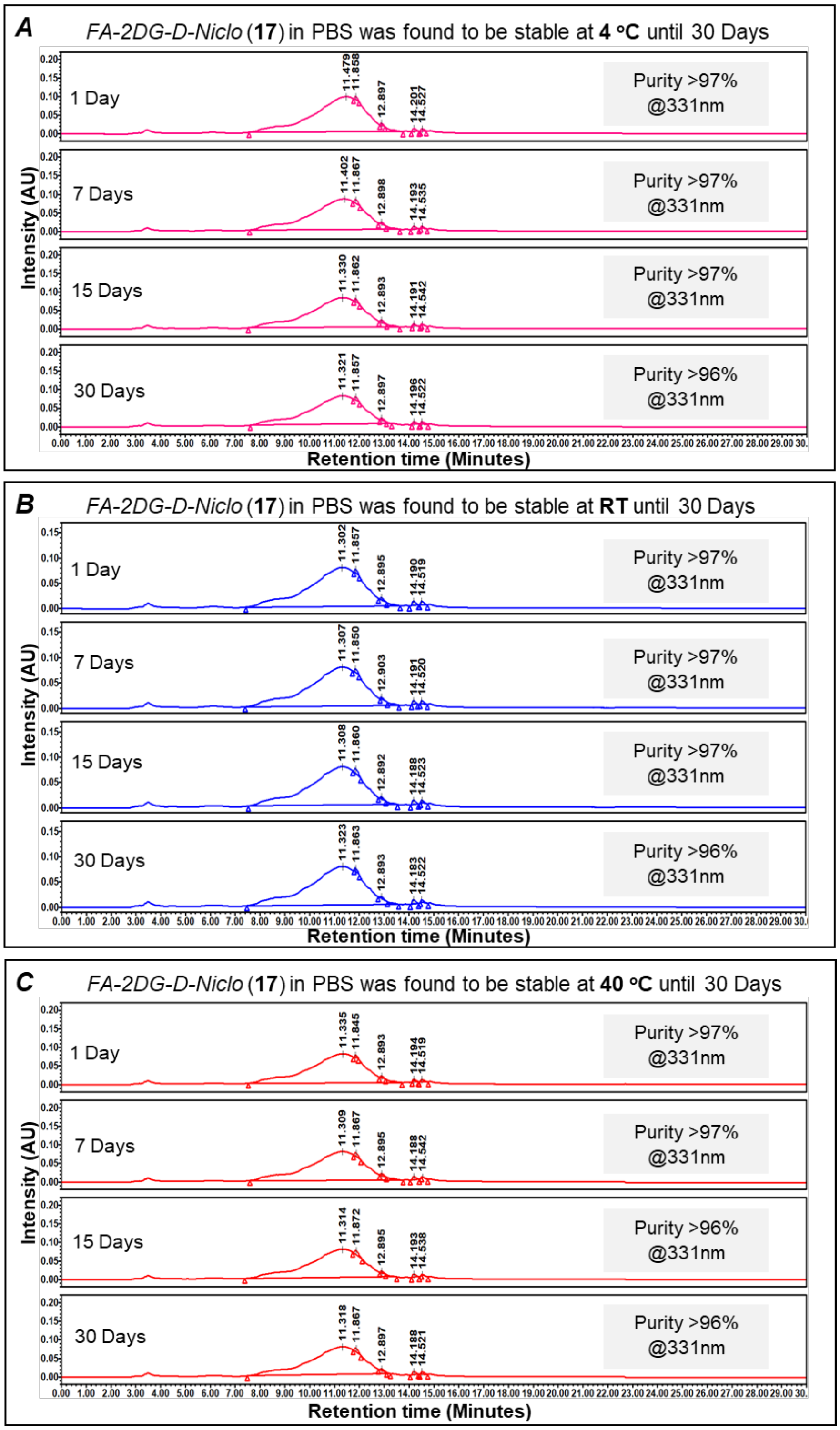


**Figure S34:** Formulation of FA-2DG-D-Niclo (**17**) in PBS was found to be stable at 4ºC (**A**), RT (**B**) and at 40ºC (**C**) until 30 days.
